# Measuring age-dependent viscoelastic properties of organelles, cells and organisms via Time-Shared Optical Tweezer Microrheology

**DOI:** 10.1101/2023.10.17.562595

**Authors:** Frederic Català-Castro, Santiago Ortiz-Vásquez, Carmen Martínez-Fernández, Fabio Pezzano, Carla Garcia-Cabau, Martín Fernández-Campo, Neus Sanfeliu-Cerdán, Senda Jiménez-Delgado, Xavier Salvatella, Verena Ruprecht, Paolo-Antonio Frigeri, Michael Krieg

**Affiliations:** ICFO - Institut de Ciències Fotòniques, Castelldefels, The Barcelona Institute of Science and Technology, Barcelona, Spain; Center for Genomic Regulation (CRG), The Barcelona Institute of Science and Technology, Barcelona, Spain; Institute for Research in Biomedicine (IRB Barcelona), The Barcelona Institute of Science and Technology, Barcelona, Spain; ICREA, Pg. Lluis Companys 23, Barcelona 08010, Spain; Universitat Pompeu Fabra (UPF), Barcelona, Spain; Impetux Optics, Spain

**Keywords:** Optical tweezers, microrheology, cell biophysics, mechanobiology, cell nucleus, LLPS

## Abstract

Recording the mechanical response of biological samples, the cell’s interior and complex fluids in general, would enable deeper understanding of cellular differentiation, ageing and drug discovery. Here, we present a time-shared optical tweezer microrheology (TimSOM) pipeline to determine the frequency- and age-dependent viscoelastic properties of biological materials. Our approach consists in splitting a single laser beam into two near-instantaneous time-shared optical traps to carry out simultaneous force and displacement measurements with sub-nanometer and sub-picoNewton accuracy during sinusoidal perturbations. Leveraging numerical and analytical models, we find solutions to commonly encountered deviations, to build a practical and robust nanorheometer. We demonstrate the versatility of the technique by 1) measuring the phase transitions of an ageing biomolecular condensate, 2) quantifying the complex viscoelastic properties of three intracellular compartments of zebrafish progenitor cells, and, 3) using *Caenorhabditis elegans*, we uncover how mutations causing nuclear envelopathies soften the cytosol of intestinal cells during organismal age. Together, our advances afford rapid phenotyping of material properties inside cells and proteins blends, opening avenues for biomedical and drug screening applications.

## 1 Main

Most if not all fluids, cells and their organelles in a human body consist primarily of water, sugar and proteins. The many weak interactions between the individual molecules give rise to complex mechanics, which are characterized by a frequency-dependent, viscoelastic response to self-generated and external forces. Such frequency-dependent mechanics are thus important for mechanotransduction, morphodynamic events and motor protein efficiency. How complex materials deform and flow in relation to their chemical composition is the science of rheology[1]. Rheological characterizations of bulk material become problematic when the sample is precious and little material is available, or is expected to demonstrate substantial heterogenity[2]. The correct understanding of the rheological properties, however, is important for chemical engineering of gels, blends and emulsion, food processing[3], drug delivery[4, 5] and health and disease[6]. The rheology of biological fluids is crucial for a variety of physiological and pathological processes[7, 8], including cell division[9], cell migration[10], mechanotransduction[11, 12, 13] and intracellular transport [14]. Recent data indicates that alterations in the way cells and their components react to mechanical forces are linked to various diseases, such as cancer and neurodegeneration[8, 15, 16, 17]. In particular, many phase-separated, liquid-like condensates exhibit age-dependent mechanical properties[18, 19, 11]. Their transition from a liquid-like to a gel or glassy state has been associated with a poor prognosis of many neurodegenerative disorders[18]. Therefore, obtaining precise characterizations of their time-dependent microrheological properties could yield valuable insights for drug development and diagnosis.

Many techniques have been put forward that afford the characterization of the cell’s rheological properties, including atomic force microscopy (AFM), Brillouin spectroscopy, and genetically encoded reporters for stress and strain[20]. Despite the wealth and versatility of these methods, many cannot simultaneously exert forces and measure their resulting effects on cell mechanics and mechanotransduction. Whereas AFM can measure a large range of forces in physiologically relevant pN-nN range and deformations in the nm-μm range, measurements are restricted to the cell surface and cannot report on the mechanics of specific cellular compartments. Brillouin microscopy has advantages as it can non-invasively measure mechanical moduli of intracellular compartments but neither be used to measure nor apply forces.

Optical tweezers are particularly well suited to derive material properties of biological materials as they operate in the pN range characteristic for molecular interactions and can measure the position of micron-sized objects with sub-nanometer accuracy in three dimensions using non-invasive infrared laser light[21]. The small scale of deformations also permits the analysis of the heterogeneity of the response, rather than providing a bulk-modulus from macroscopic measurements, a key point for experiments aimed to understand subcellular mechanical compartmentalization. Optical tweezer-based, active microrheology is a powerful technique for probing the mechanical properties of complex fluids in biological systems, such as the cytoplasm of living cells [9, 22, 23, 24, 15, 25], cancer spheroids [26], in living animals [27, 28, 29] and biomolecular condensates [19, 11, 30, 31]. This method involves the measurement of the motion of small particles suspended within a fluid (or embedded in a viscoelastic gel), due to an oscillating optical trap [32, 33]. The frequency-dependent probe displacement 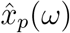 in response to the active force 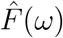 captures the elasticity and viscosity of the material over the observed rheological spectrum. Moreover, optical tweezers can stably trap small microspheres (with radius *r*_bead_ ≈ 10^−7^m; [34, 24]) making them particularly useful to measure their response to force embedded into very small volumes (e.g. *V*_cell_ = 10^−15^ vs. *V*_bead_ = 10^−19^ m^3^), e.g. in living cells, purified protein blends or novel synthetic materials and therefore afford applications where samples are precious. The small scale of deformations also permits the analysis of the heterogeneity of the response, rather than providing a bulk-modulus from macroscopic measurements, a key point for experiments aimed to understand subcellular mechanical compartmentalization.

The measurement of the mechanical and rheological properties requires the simultaneous recording of stress and strain. Using optical tweezers, this is equivalent to measuring the optical force acting onto an optically-trapped probe and its displacement resulting from such force. Often, active microrheology is implemented by stage driving while recording the force in a calibrated trap in situ[35, 36, 37]. Due to the induced sample motion, stage driving is incompatible with simultaneous optical observations by fluorescence microscopy. A solution to this is a setup with two lasers and two detection optics: one laser for driving displacements and one for measuring forces. Due to the complexity of this implementation, only a handful of setups exist worldwide[9, 38, 39, 40, 25].

Here, we present an active microrheology method with a single time-shared laser[41] generating two distinct optical traps - one that drives the active oscillation, while the second one acts as a static, displacement detection beam (Fig. 1a). We thus name them *driving* and *static* trap, respectively. We first theoretically established that a fast, 25 kHz beam sharing generates a quasi continuous trapping potential for simultaneous force and displacement measurements. Because the two traps appear intermittently and complementary to each other, we introduce corrections to compensate for the deviation of the raw measurements obtained by time-sharing compared to the expected result for two simultaneous traps.

**Fig. 1.**
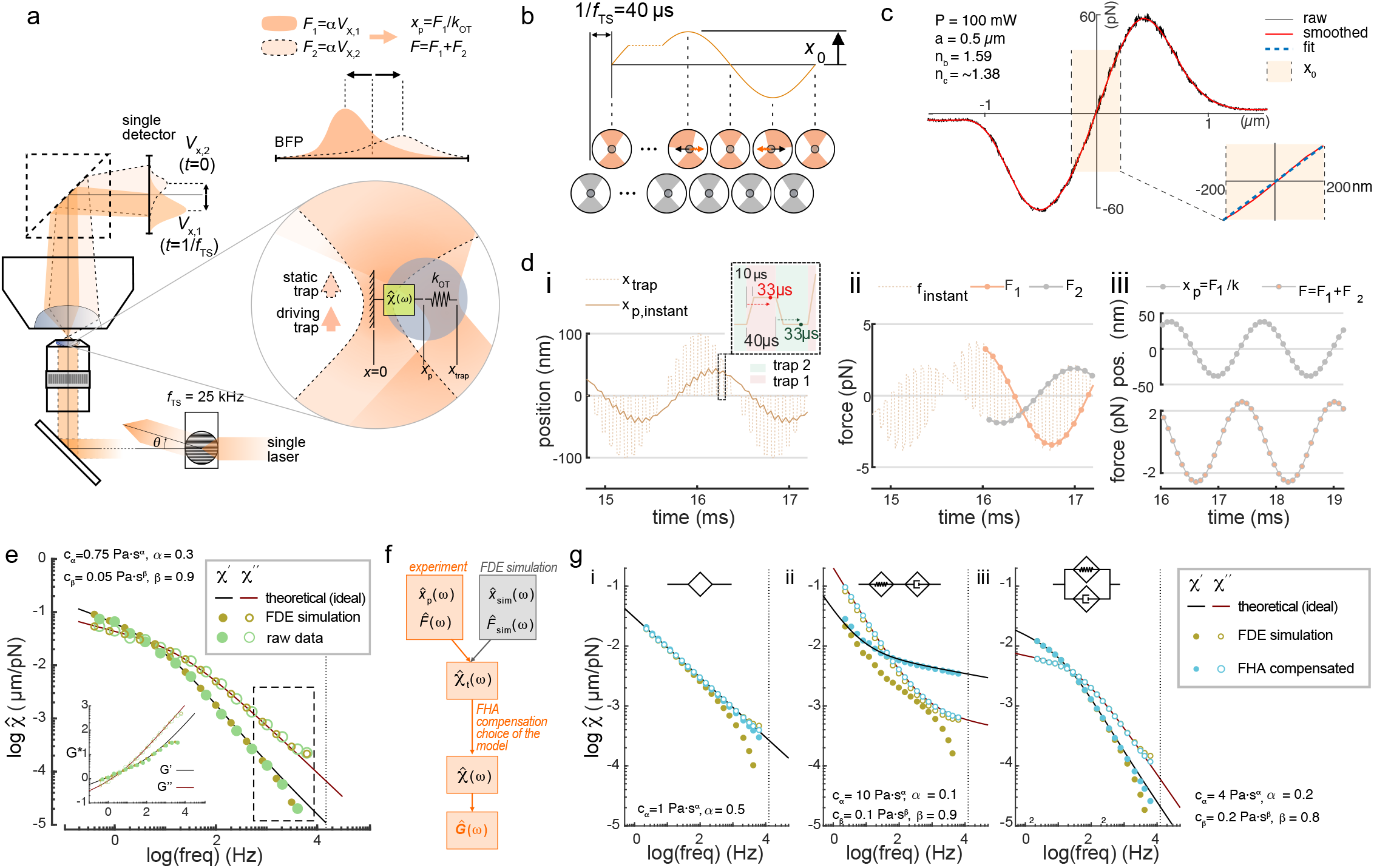
Time-shared optical tweezer microrheology. **a**, Schematic of time-shared optical tweezer microrheology with direct light momentum sensing of optical forces. A single 1064-nm laser beam is time-shared at 25 kHz between a *driving* (1) and a static *detection* (2) trap. The driving trap (orange) oscillates around the trapped particle, while the static trap (light orange, dashed line) monitors the particle position as *x*_*p*_ = *F*_2_*/k*. For clarity, only the spring for the driving trap was indicated, noting that both traps have the same spring constant. The optical force acting onto the probe particle corresponds to the addition of forces exerted by the two traps: *F* = *F*_1_ + *F*_2_, which are obtained as *F*_1,2_ = *αV*_1,2_, where *α* is here the volt-to-picoNewton conversion factor of a single, light-momentum direct force sensor. **b**, Time-sharing position and force measurement sequence. While trap 2 remains motionless at the optical axis in order to detect bead displacements through BFP interferometry, trap 1 applies an active sinusoidal perturbation with amplitude *x*_0_ at the time sharing frequency 1*/f*_*T S*_. The schematic on the bottom represents the deflection of the laser beam by the trapped particle for the driving (orange) and static trap (grey). **c**, Force profile acquired by sweeping the trap across a 1 μm polystyrene microsphere embedded in the cytoplasm of a zebrafish cell. The shaded area indicates that force is linear with displacement over the amplitude of the rheology routine. Blue dotted line is *F* = *k* · *x*. **d-g**, Quantitative description of bead motion in TimSOM. **d**, Simulation of the instantaneous position (i) of the probe particle in water using the *fractional derivative equation* method (*A* = 100 nm, *f* = 625 Hz, *k* = 50 pN/μm, water viscosity: *η* = 10^−3^ Pa·s) and the resulting instantaneous optical force acting onto the probe (ii). Interleaved force values for the static and driving traps, sampled at *f*_*T S*_*/*2 = 12.5 Hz with a delay of 33 μs (Supplementary Text 2), are indicated as thin dashed line. Inset of (i) shows the time sharing properties of the active (trap 1) and passive trap (trap 2) with the rise time of 10 μs. In ii, the probe position and in iii, the total force are shown. **e**, Response function derived from the FDE simulation and experimental data acquired in a zebrafish progenitor cell using the time-shared microrheology routine. Solid lines show the expected, ideal behavior of a fractional Kelvin-Voigt material. The inset shows the complex shear modulus. The dashed box indicates high frequencies with expected deviations due to the non-simultaneous measurement of stress and strain. **f**, Analytical pipeline to retrieve the G modulus from the deviated measurements and/or simulations. **g**, Response function (*χ*^′^, storage; *χ*^′′^, loss) of the ideal scenario (theoretical), the time-shared simulations (FDE) and the compensated data points (FHA) for a (i) single springpot; (ii) fractional Maxwell and (iii) fractional Kelvin Voigt model. Parameters used for simulation indicated in each panel. Legend is indicated on the right of the panel.

To demonstrate the versatility of the time-shared optical tweezer microrheology (TimSOM), we measured how the rheological spectrum ranging from 0.5 Hz to 2 · 10^3^ Hz changes with time, development and age, using biomolecular condensates, cells and animals as a model system. We chose zebrafish and *C. elegans* as examples for our technology, for their rich mechanobiology [42, 43, 44, 45] and to further enrich our knowledge of the importance of the material properties for morphogenesis and age. Lastly, to facilitate the use of TimSOM in the lab, we provide a detailed ‘benchtop’ protocol in the supplementary material.

## 2 Results

### 2.1 Concept, theory and experimental implementation of time-shared optical tweezer microrheology

To conceptualize a simplified microrheology experiment, we consider a *single* laser source to drive the probe and measure the resulting displacement. The laser is time-shared and generates two traps that alternate between two positions. In other words, half of the time the laser generates a trap on position 1 and the other half on position 2. Thus, time-sharing generates a discontinuous, but quasi-simultaneous stress/strain measurement (Fig. 1a, b). The advantage is a reduced complexity and alignment, and, as a consequence of the shared laser source, the traps have identical power, position sensitivity *β*_1_ = *β*_2_ ≡ *β* and trapping stiffness *k*_1_ = *k*_2_ ≡ *k/*2, where *k* is the sum of the stiffness of the two traps (Fig. 1a, SI Text 1.2).

We envision that at the beginning of the routine, both traps are centered on the microsphere (Fig. 1b). Trap 1, the *driving trap*, starts to oscillate at a given frequency *ω* and amplitude *x*_0_, i.e. *x*_1_(*t*) = *x*_0_ sin(*ωt*). Trap 2, the *static trap*, remains fixed at *x*_2_(*t*) = 0 (Figure 1b). The amplitude must be in the linear range of the trap (*x*_0_ = 200 nm; Fig. 1c) and therefore smaller than the radius of the a microsphere (*a* = 0.5 *μ*m). Thus, the probe particle will feel a force from both traps. Because the traps are of equal strength, we expect that the static laser exerts a force on the probe in opposite directions and back into the starting position. Further, because force and position measurements are interleaved, we reasoned that the measured microsphere trajectory deviates from the ideal case (SI Text 1.4).

To model and understand the effects due to time sharing, we first simulated the displacement of the particle under the influence of the two intermittent traps solving the equation motion under an external force in a viscoelastic matrix that is best described by power laws [46]. In particular, we consider the fractional Kelvin-Voigt or Maxwell models [47]. These are generalized viscoelastic models that extend the applicability of the classical Kelvin-Voigt, Maxwell and structural damping models. They have been used to describe the power-law behaviour of many biological samples and gels[47], including the intracellular cytoplasm[9, 48, 49] and protein condensates.

First, we numerically computed the probe trajectory using the fractional derivative equations (FDE) that describe motion in materials with power law rheology (SI Text 1.5). We found that each trap influences the instantaneous probe trajectory, *x*_p,instant_(*t*), leading to a sawtooth shape (Fig. 1d). We reasoned that this deviation from the instantaneous trajectory depends on the viscoelastic properties of the probed material, and the rate at which is deformed. We thus ask how does this generally affect the response obtained from the TimSOM measurement in viscoelastic media?

For viscoelastic materials, the response 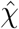 of the microsphere, as it transitions from the ‘current’ to the ‘new’ position under force depends on the stimulation frequency (*ω*). This is described with the complex response function 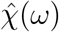 and the complex shear modulus *Ĝ*(*ω*), which consist of a real part and an imaginary part, typically denoted as *Ĝ* = *G*^′^ + *iG*^′′^. *G*^′^ is the storage modulus and *G*^′′^ is the loss modulus (see SI Text 1.2 for the derivation), which describe the elastic and the viscous behavior of the material, respectively.

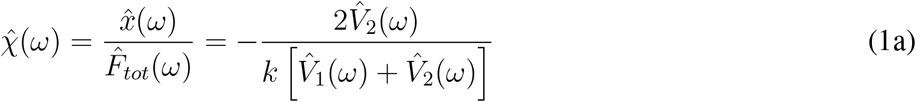

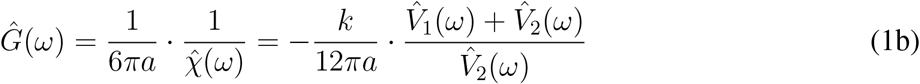

Interestingly, due to the symmetry between the two traps, equations 1 do not depend on the displacement sensitivity *β* (μm/V) (see SI Text 1.2). Thus, the accuracy of our method strictly depends on the ability to generate two identical traps with the same stiffness, *k* (pN/μm), from a single laser source, as well as the bandwidth of the measurement of the relative voltage change (SI Text 1.3). This is straight-forward in a setup modulated with fast-scanning acusto-optic deflectors (AODs) [50, 51] and equipped with direct light momentum force detection (Fig. 1a), where *V*_1_ and *V*_2_ are the voltage signals of a PSD placed at the back focal plane (BFP) of a direct force measurement sensor [52].

We used equation 1 to calculate the response of the microsphere over frequencies ranging from 0.1Hz up to the Nyquist frequency (6.25kHz), and we found that it indeed deviated significantly from the ideal response function at high frequencies (beige points vs solid line in Fig. 1e). At this point, we straightforwardly implemented TimSOM in our setup, with the aim to test this prediction. Indeed, when we performed a rheology routine in the viscoelastic cytoplasm of a living cell, we intriguingly observed the same deviation from the ideal fractional Kelvin-Voigt behavior, as predicted by the FDE simulation (Fig. 1e).

Next, to understand and eventually correct for this deviation, we obtained an analytical solution of the probe motion —and the resulting 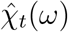 — through a first harmonic approximation (FHA) of the trap positioning sequence in the frequency domain (for a derivation see SI Text 2 and Ref. [53]). Our analytical function predicts the time-sharing response function from the ideal, 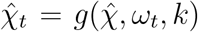, which also depends on the time-sharing frequency, *ω*_*t*_ = 2*πf*_*t*_, and trap stiffness, *k*. This function can then be inverted to retrieve the artefact-free response function after compensating the deviated measurement (Fig. 1f,g, SI Text 1.12).

To show that the compensation can be applied generally and indeed is able to retrieve the ideal, artefact-free response function (solid lines in Fig. 1g), we first simulated trajectories and the response function using the fractional derivative equations in various materials, such as power law and the fractional Maxwell and Kelvin Voigt material (beige points in Fig. 1g, SI Text 1.11). For viscoelastic liquids, e.g. Maxwell materials, the deviations strongly affect the real part of the response function, such that the solid behaviour at frequencies larger than the crossover frequency is not accessed for inadequately stiff traps: The bead jumps instantaneously between the two traps which creates a deviation from the expected viscoelastic behaviour in the high frequency range of materials where the elastic contributions dominate. Strikingly, when we applied the compensation routine, we recovered an almost perfect match (celeste point in Fig. 1g) to the ideal behavior. Subtle deviations remain at high frequencies which could be minimized by incorporating higher harmonics into the compensation. We therefore confidently conclude that time-sharing measurements correctly reflect the behaviour for viscoelastic solids, e.g. Kelvin-Voigt materials — up to 2000 Hz, deviations from ideal behaviour a less than 15%.

With this formalism in hand, we are now equipped to measure the mechanics of most common biological materials, including cells, tissues and protein blends.

### 2.2 TimSOM measures 0.1 Pa-100 kPa over five decades of frequencies

To experimentally validate the TimSOM scheme, we used three different materials with known rheological properties to extract the shear modulus through TimSOM over a frequency range from 2 Hz to 6.25 kHz. The viscosity of water and the different glycerol mixtures extracted from TimSOM were consistent with the values extracted from drag force measurements of a microsphere driven at different velocities (Extended Data Fig. 2a,b). In the Fourier domain, the *F*_1_ and *F*_2_ peaks corresponding to the active oscillations applied stand out by three orders of magnitude with respect to the noise baseline (Extended Data Fig. 2c i). We then fabricated different polyacrylamide gels ranging from 20Pa-100kPa (see Methods). We first characterized the soft gels with a standard method to extract the complex shear modulus using creep compliance *J*(*t*) measurements [54] (Extended Data Fig. 2e). When we performed TimSOM on the same gel, we recovered a frequency-dependent shear modulus that was described well by a fractional Kelvin-Voigt model (Extended Data Fig. 2e) in close agreement to the creep-compliance measurements. For the stiffest gels (20% acrylamide) we measured a plateau modulus of ≈30kPa that reached 100kPa at the short time scales. Together, due to its exceptional displacement sensitivity and force resolution, TimSOM measured the rheological properties of stiff gels over a wide band of frequencies to capture the rubber-glass transition that was previously reported[55]. Again, the frequency-domain active peaks were at least one standard deviation higher than the noise baseline, though a larger noise arose around 10 to 100 Hz due to mechanical coupling (Extended Data Fig. 2c ii).

Lastly, we also tested TimSOM on microspheres embedded in freshly prepared and uncured 10:0.1 PDMS prepolymer/curing agent mixtures (Extended Data Fig. 2f). Consistent with previous reports, the pre-gelation PDMS was best described as a fractional Maxwell viscoelastic gel[47] with a plateau modulus at high frequencies of ≈50kPa[56, 28]. This corresponded to a minimal displacement of the microsphere of less than 1 nm. Taken together, TimSOM is suitable to investigate the storage and loss moduli of viscous and viscoelastic materials with different properties, and retrieves viscoelastic moduli over five orders of magnitude over a five decade frequency range.

### 2.3 Deciphering the viscoelastic properties of biomolecular condensates

Biomolecular condensates (BMCs) ensue from liquid-liquid phase separation and are relevant for the organization of the cytoplasm of living cells[57, 58]. The correct localization and material properties of these BMCs are essential for proper cellular functions[11] and are thus under tight cellular control [59]. The unregulated formation and transition of condensates from liquid to solid aggregates often correlate with disease propagation where a higher degree of stiffening correlates with disease prognosis.

Recently, laser tweezer rheology revealed that many BMCs behave as ageing Maxwell liquids, with an age-dependent change in viscosity[19]. These experiments are commonly performed in a dual optical trap, in which a droplet is sandwiched between two trapped, active and passive microspheres (Fig. 2 and Supplementary Video S1 [19, 60]). The complex shear modulus and surface tension of the material is then extracted from the active force measurement and passive bead displacement, with the assumption that the microsphere radius is considerably smaller than the protein droplet and that the viscosity of the dilute phase is known [61]. To relax these assumptions and simplify the experiment with a single trap, we performed TimSOM on MEC-2 BMCs as a model for an age-dependent maturation process and compared these results to the dual trap assay[11] (Fig. 2 and Supplementary Video S2). After applying the compensation routines for a fractional Maxwell material, we found a similar frequency response in both configurations that suggested a significant and noticeable shift in the cross-over frequency to lower values after 24 h of droplet formation (Fig. 2a, b). Interestingly, the values for stiffness and viscosity were slightly higher in the single bead assay, which measured the bulk modulus in the center of the droplet as opposed to the interfacial moduli. These differences suggest that the maturation process does not happen homogenously across the BMC instantaneously but affects center and periphery differently[62]. To test if TimSOM can be applied to other BMCs, we tested BMCs composed of *cytoplasmic polyadenylation element binding protein 4* (CPEB4), an RNA-binding protein that regulates translation through cytoplasmic changes in poly(A) tail length, with links to idiopathic autism spectrum disorder [63] and with previously uncharacterized shear modulus[64]. In their ‘naive’ phase (within 2h after droplet formation *in vitro*), the mechanical response function confirmed their fractional Maxwell behavior (Extended Data Fig. 3a), and the BMCs displayed a ≈10x slower relaxation time scale an increased viscosity (Extended Data Fig. 3b) compared to MEC-2/UNC-89. Moreover, we found that the CPEB4 condensates stiffen very quickly after formation, visible by the appearance of fibers (Extended Data Fig. 3c), which prevented rheological characterization within the limit of the instrument. Taken together, TimSOM accurately resolved the (fractional) Maxwell behaviour of BMCs and uncovered a potential mechanism for the rigidity percolation transition in MEC-2 across the condensate.

**Fig. 2.**
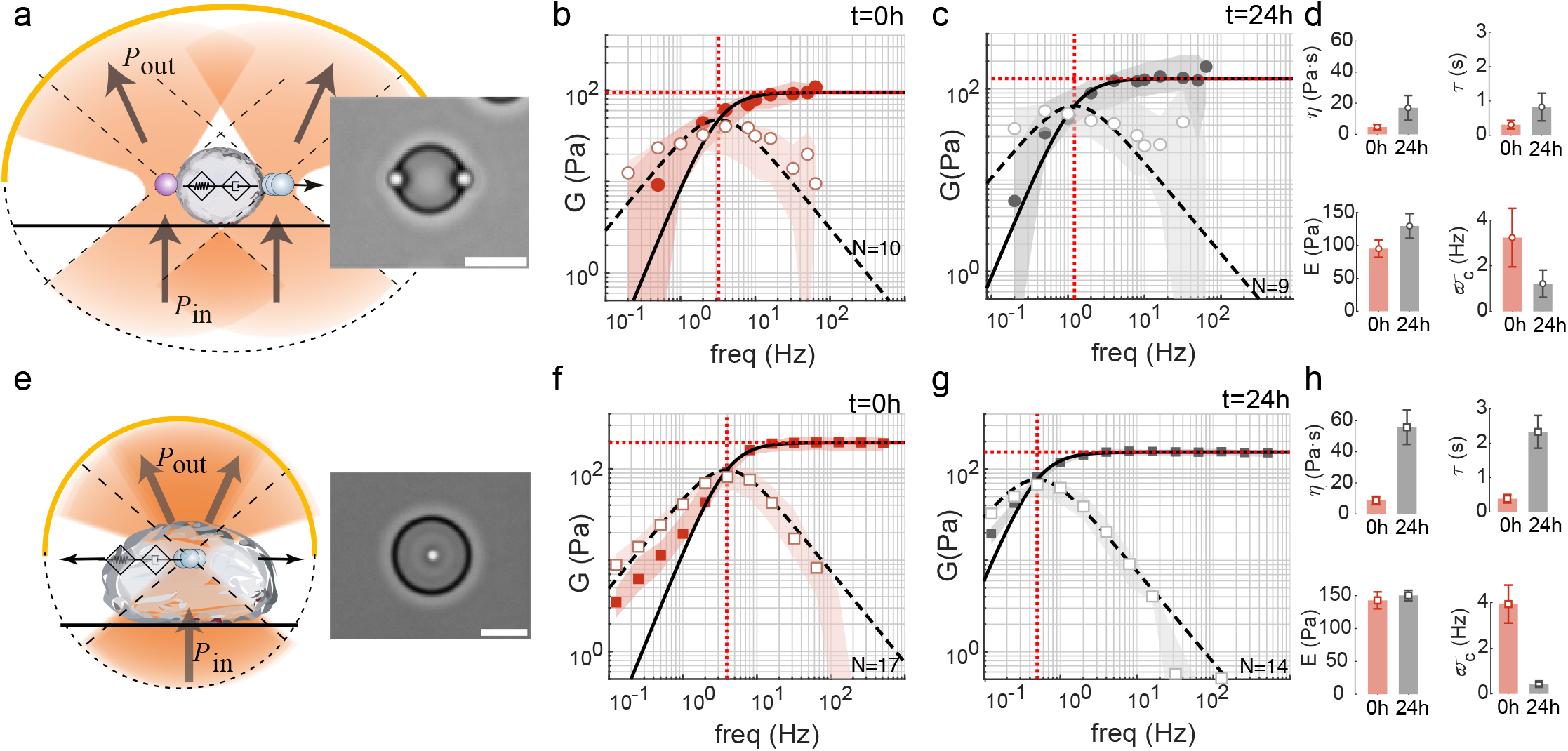
Viscoelastic properties of biomolecular condensates. **a-d** Schematic and snapshot of a MEC-2/UNC-89 protein droplet measured with a pair of optically trapped PEG-terminated microspheres in a dual optical trap. Scale bar = 10 μm. *P*_*in*_, *P*_*out*_ define the light momentum before and after interacting with the trapped microsphere. See also Supplementary Video 1. **b, c**, Storage (*G*^′^(*ω*), filled circles) and loss (*G*^′′^(*ω*), open circles) moduli measured in the dual optical trap at (b) *t* = 0h and (c) *t* = 24h after condensate formation. The solid and dashed lines are the real and imaginary parts of a non-fractional Maxwell G modulus fitted to the acquired data. Circles and shadows are the median and ±25% quantiles over N=10 (b) and N=9 (c) measurements. **d**, Variation of the fitting parameters showing the change in the dynamic viscosity, *η* (Pa·s), stiffness, *E* (Pa), time constant, *τ* = *η/E* (s) and crossover frequency, *ω*_*c*_ = 1*/τ* (Hz) over 24h condensate maturation in the dual optical trap. **e-h**, Schematic and snapshot of a TimSOM experiment on MEC-2/UNC-89 protein droplet with an embedded carboxylated microbead. See also Supplementary Video 2. **f, g**, Storage (*G*^′^(*ω*), filled squares) and loss (*G*^′′^(*ω*), open squares) moduli measured with TimSOM at (f) *t* = 0*h* and (g) *t* = 24*h* after condensate formation. The solid and dashed lines are the real and imaginary parts of a non-fractional Maxwell G modulus fitted to the acquired data. Squares and shadows are the median and ±25% quantiles over N=17 (f) and N=14 (g) measurements. **h**, Variation of the fitting parameters over 24h extracted from the TimSOM routine.

### 2.4 The nucleoplasm is a soft viscoelastic solid

Direct, momentum based optical force measurements have their greatest potential inside living cells, where traditional mechanics measurements are limited by time consuming and sample-variant *in situ* calibrations[65, 24]. However, extracting information about their mechanics relies on the simultaneous measurement of deformation using a second detector (see above), in addition to the force. Thus, Tim-SOM should facilitate intracellular rheology without sacrificing simplicity and throughput. To demon-strate TimSOM inside cells, we performed active microrheology routines of three different compartments inside zebrafish germ layer progenitor cells. Following methods described previously[51], we introduced microspheres into individual zebrafish embryos and extracted cells 4 hours-post-fertilization (hpf). We trapped individual microspheres to perform active microrheology routines in the cytoplasm and at the nuclear interface in the frequency range from 0.5 Hz-2 kHz using a 50-90 mW optical trap at the sample plane and calculated the complex shear modulus from the corresponding response function (see Supplementary Text). When performing the routines in the cytoplasm, we did not notice a significant history effect or non-linear mechanical response for the duration of the rheology routine, as we measured a constant phase lag between the trapping force and the probe oscillation throughout the measurement, and neither for repetitive measurements in the same cell (e.g. heating related to laser power)) (Extended Data Fig. 4). To compare the rheological properties between different cells and conditions, we used the fractional Kelvin-Voigt model which captures the frequency-dependent response of composite materials and describes the curves with 4 parameters ([9, 47], Supplementary Text 3). The frequency-dependent G modulus revealed a viscoelastic response with an elastic plateau at low frequencies (Fig. 4a) and a viscous response dominated at high frequencies. The cross-over frequency at which the dissipative forces dominate was ≈5 Hz and above, the cytoplasm fluidified as indicated by exponent *β* approaching 1 (Extended Data Fig. 5c). Thus, the cytoplasmic rheology is consistent with a shear-thinning viscoelastic fluid, with a rubber-glass transition at high frequencies. We found that the cytoplasm is stabilized by a synergistic effect of the actin and microtubule cytoskeleton: pharmacological perturbation of actin and tubulin polymerization lead to decreases in *C*_*α*_ and *C*_*β*_, indicating a reduction in the magnitude of the elastic and viscous modulus [48]. Both elements, however, have a different contribution to the low frequency response: Interestingly, depolymerization of the MT cytoskeleton decreases the low frequency power law exponent *α*, different to what was observed in dividing tissue culture cells[9], which points towards a partial solidification of the cytoplasm. This may be an indirect effect of MT on the actin cytoskeleton[66], as it is known that Rho-GEF are bound to the MT lattice which gets released upon MT depolymerization to exert its effect on actin dynamics by increasing GTPase activity[67]. Indeed, upon actin depolymerization, the effect of nocodazole on *α* is reversed and the cytoplasm is again more liquid-like (Extended Data Figure 6a). Taken together, the complex shear modulus was lower for all frequencies after depolymerization of the actin (Fig 3a and Extended Data Fig. 5d) and microtubule cytoskeleton (Extended Data Fig. 6a), but not after disrupting myosin II motor activity (Extended Data Fig. 6b), indicating that the cytoskeletal network integrity rather than contractility had the major effect on the rheological signature of the cytoplasm.

**Fig. 3.**
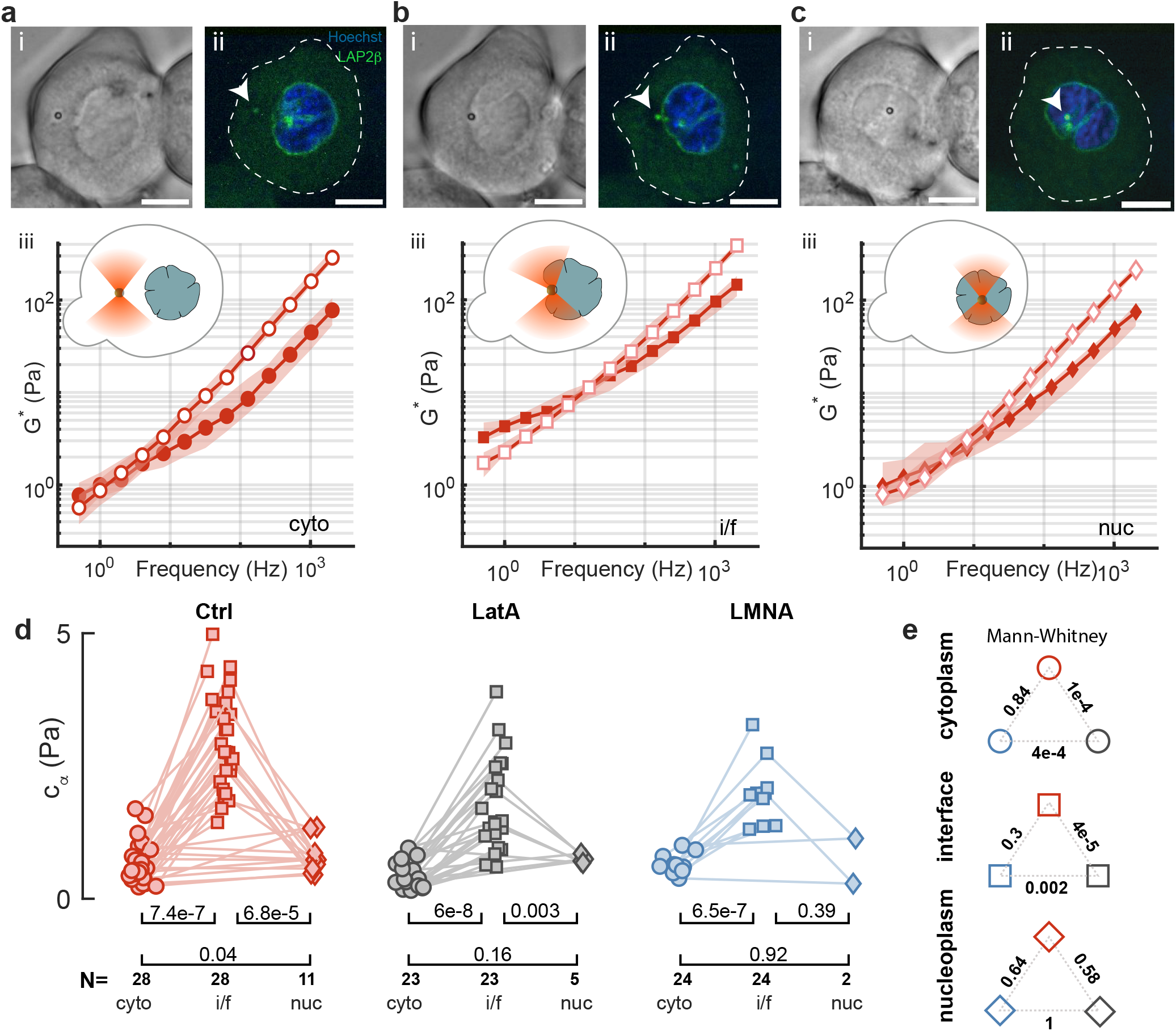
Cytoplasm versus nuclear rheology. **a-c**, Representative (i) brightfield and (ii) confocal imaging of a zebrafish progenitor cell stained with Hoechst (blue) to label the nucleus and expressing Lap2*β*:GFP (green) with a microsphere in its (a) cytoplasm (cyto), (b) nuclear interface (i/f) and (c) inside the nucleus (nuc). (iii) Frequency spectrum of the complex G modulus indicating the storage (closed symbols) and loss modulus (open symbols) of the three corresponding compartments. Solid line is a fit of the fractional Kelvin-Voigt model to the data. Scale bars = 10 μm. See also Supplementary Video 3 for the complete routine. **d**, Stiffness (*C*_*α*_) of the cytoplasm, nuclear interface and nucleoplasm for control and actin depolymerization (LatA) conditions as extracted from the fit of a fractional Kelvin-Voigt model to the rheological spectrum. Lines connect paired datapoints that were acquired in the same cell with the same microsphere. p-values above the brackets derived from a paired t-test. See Extended Data Fig. 5 and Supplementary Table 1 for the comparison of all other fit parameters and their p-values. N=number of cells used in the measurement. **e**, p-values of the indicated pairwise comparison using a 2-sided Mann-Whitney U-test for *C*_*α*_ of the cytoplasm, nuclear interface and the nucleoplasm in control, LatA treatment and Lamin A (LMNA) overexpression.

**Fig. 4.**
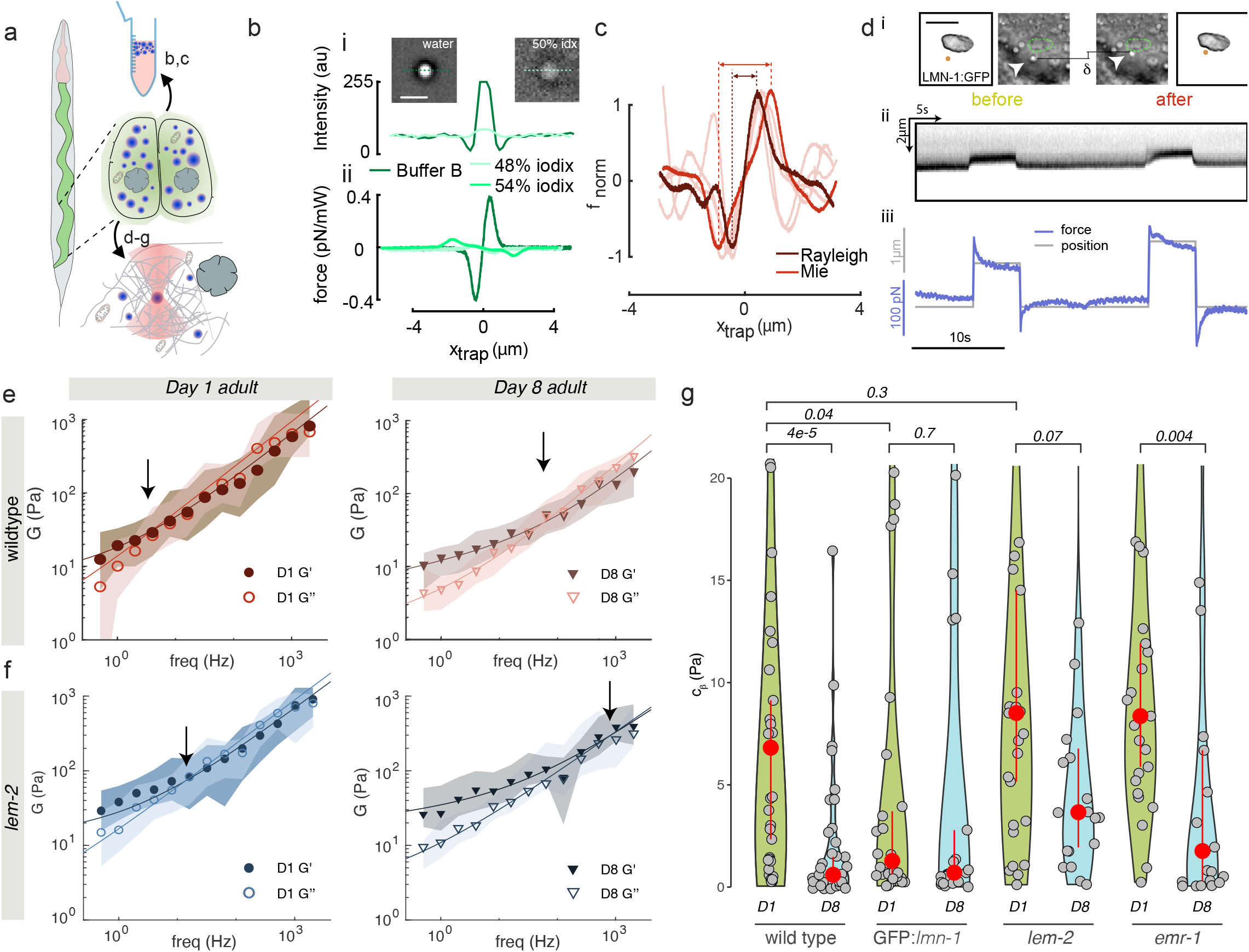
Longitudinal tissue microrheology *in vivo*. **a**, Sketch of an animal with the intestinal tissue highlighted in green and the pharynx in red. The close-up sketch shows a pair of posterior intestinal cells with lipid droplets in blue. Lipid droplets were isolated from adult animals, purified and tested under various conditions for their suitability as optical tweezer probes (b,c see Methods). For *in vivo*, individual droplets were trapped to measure the rheological response of the material in its vicinity (panels d-g). **b**, Refractive index matching with varying concentrations of iodixanol. (i) Brightfield micrograph of a lipid droplet in buffer B on the left (see Methods, representative for N=6 droplets) and in 48% of iodixanol (right, N=12). Graph shows the intensity profile along the dotted line indicated in the photograph. Scale bar = 2 μm. (ii) Force profile on a droplet in the matched conditions, for 0, 48% and 54% of iodixanol. **c**, Force scan across the lipid droplet for particle radius estimation (see Methods). Lipid droplets vary in size from the trapping force Rayleigh (dark red) and Mie (light red) limits. N=2 droplets, representative for all measurements. **d**, (i) Fluorescence of *GFP::lmn-1* and bright field images demonstrating nuclear deformation with a trapped lipid droplet upon contact during a tweezer experiment. *δ* indicates the deformation of the nucleus during the test, arrowhead points to the trapped lipid droplet. Scale bar = 2 μm. (ii) Kymograph of two consecutive step indentations of an intestinal nucleus using a lipid droplet as force probe. (iii) Force and displacement during the same step indentation protocol. **e, f**, Frequency dependent shear modulus for two different ages of (e) wildtype and (f) age-matched *lem-2* mutants. The median and ±25%quantiles are represented by lines and shadows, respectively. **g**, Viscosity (*C*_*β*_) of the cytoplasm as extracted from the high-frequency component derived from the fit of the fractional Kelvin-Voigt model to the rheological spectrum of day-1 and day-8 adults for four different genotypes as indicated. Red circle indicates median±bootstrapped 95% confidence interval. p=0.003 derived from a non-parametric Kruskal-Wallis test, followed by a pairwise comparison using a one-sided Dunn test as indicated above the horizontal brackets (for details on statistics and number of measurements see Supplementary Data Table 2 and Extended Data Fig. 10).

Surprisingly, we found that the cytoplasm was soft enough to re-position the same microsphere to the cytoplasmic/nuclear interface where we repeated the measurement with the aim to test the mechanical properties of the largest cellular organelle (Supplementary Video 3). In agreement with previous reports[68, 69], we found that the nuclear interface is stiffer than the cytoplasm (Fig. 4). In order to unravel the molecules responsible for determining the shear modulus of the interface, we conducted experiments on cells that overexpress Lamin A, a key protein found in the nuclear envelope. In parallel, we also treated cells with with Latrunculin A (LatA) to perturb the actin network. The complex shear modulus of the nuclear interface of ectopic Lamin A expressing cells was indistinguishable from control cells which do not express high levels of Lamin A endogenously[70, 71] (Extended Data Fig. 5e). We also performed a creep compliance test to measure the resistance of the nucleus to large deformations (Extended Data Fig. 7a-d). Even though the nuclear envelope was clearly more resistant on short and long time scales, its initial stiffness was unaffected by Lamin A overexpression (Extended Data Fig. 7f- j). Actin around the nucleus was previously known to dampen and transmit mechanical forces[72] and was observed in actin stainings of isolated gastrulating zebrafish progenitor cells (Extended Data Fig. 8a,b and ref [73]). We thus speculated that actin has a noticeable effect on the mechanics of the nuclear interface. Indeed, after treating the cells with 0.5 μM LatA as previously described[12], cells immediately ceased blebbing and we already noticed a significant reduction in the initial stiffness and resistance to deformation when we performed the large-strain creep-compliance test (Extended Data Fig. 7e-j). Moreover, the complex shear modulus of the nuclear interface was significantly reduced in the presence of LatA compared to control cells. However, even without a functional actin network, it remained significantly higher than that of the cytoplasm in presence of LatA (Fig. 4d,e, Extended Data Fig. 5d). As the nuclear envelope originates from the endoplasmatic reticulum (ER), we asked whether or not our measurements at the nuclear interface are affected by the properties of the ER. We first stained the ER and the nucleus and found that the microsphere is indeed in close contact with the nuclear envelope, without large accumulations of ER in between (Extended Data Fig. 8c,d). We then applied Brefeldin A, a drug that perturbs ER morphology and mechanics, to the cells at low concentrations that were used before to induce morphological alterations and mechanical changes of the ER[74]. However, compared to the actin cytoskeleton, the ER had little influence on our measurements (Extended Data Fig. 8e). Together, these data show that actin increases the shear modulus of the nuclear/cytoplasmic interface which is unaffected by Lamin A expression at the inner nuclear membrane.

To better understand the rheological properties of the nucleus itself, we mechanically inserted microspheres into the nucleoplasm (Extended Data Fig. 7). Despite the relatively large shear moduli of the nuclear interface compared to the cytoplasm (Fig. 4a,b), the envelope was very flexible and deformed easily under modest forces exerted by the optical trap. When we applied a constant force of 100-150 pN onto the nuclear envelope, the microsphere entered into the nucleus which afforded the possibility to independently measure the rheology of the nucleoplasm (Fig. 4c, Supplementary Video 4). Importantly, the nuclei retained their integrity and functionality after microsphere insertion, as we did not observe leakage of DNA or soluble GFP (Supplementary Video 5, Supplementary Text 3.2) from the nucleus into the cytosol, and cell division proceeded indistinguishably from untreated cells with correct nuclear envelope breakdown and reassembly (Supplementary Video 6 and 7), with absence of the apoptotic marker Annexin V up to 1h after the experiment (not shown). The large deformation of the nuclear envelope during the insertion did not activate known mechanotransduction pathways as judged by the absence of cortical myosin II relocalization [12, 75]. Likewise, the rheological characteristics of both the cytoplasm and the nuclear interface remained largely unchanged after the insertion, mirroring their state prior to the indentation, suggesting that the nuclear deformation does not incur cytoskeletal softening [76] (Supplementary Text 3.2). Our rheology measurement of the nucleoplasm revealed a comparably soft material similar to the cytoplasm. The complex shear moduli, however, did not change in presence of LatA and the expression of Lamin A (Extended Data Fig. 5), indicating that the mechanical properties of the nucleoplasm is not influenced by the actin cytoskeleton and the nuclear envelope. Together, this indicates that with the same microsphere, we are able to measure the frequency dependent mechanical response of three cellular compartments and that the nucleoplasm is composed of a soft, viscoelastic material.

### 2.5 Premature ageing affects cytoplasmic viscoelasticity

Next, we applied TimSOM in specific tissues of *C. elegans*, a model system to understand the factors for organismal age. Ageing is a multifactorial process under genetic control[77], but how organismal age affects the mechanical properties of cells and tissues, however, is poorly understood. As a proof of principle, we specifically asked if the cytoplasm displays changes in rheological properties during the first eight days of age. Due to the difficulty in introducing microspheres into adult tissues without affecting animal physiology, we first established the use of endogenous lipid droplets as mechanical stress probes. Recent data demonstrates that cellular lipid droplets resist deformation when subjected to forces[78], cause itself nuclear indentations[78, 79] and activation of mechanotransduction pathways leading to hepatocyte differentiation[79]. Such droplets are abundant in various *C. elegans* tissues, including the intestinal epithelium (Fig. 4a) and the epidermis, where they were observed to mechanically rupture the nucleus without being deformed, and thus have a great potential to serve as endogenous force probes[80].

To establish these lipid droplets as *in vivo* force probes for rheology in *C. elegans*, we first confirmed that 1) their refractive index (*n*) is significantly higher than the surrounding cytoplasm (Fig. 4b), 2) they are ≈1μm in diameter to maximize trapping stiffness[34, 24] (Fig. 4c) and, 3) they are sufficiently stiff to be able to indent the desired target [78, 80]. We thus isolated lipid droplets from *C. elegans* using established procedures[81] and used iodixanol (idx) as index matching media [82] to determine their refractive index. At an idx concentration of 48%, the droplets were indistinguishable from the surrounding medium, which corresponds to n=1.42 (Fig. 4b). Thus, the lipid droplet’s *n* was significantly larger than the cytoplasm(n=1.33-1.37, [83]). The diameter of the droplets was measured by scanning the optical trap across and applying a diameter look-up-table correction obtained from numerical simulations based on the T-matrix method [84] (Fig. 4c, Extended Data Fig. 9). In order to confirm the suitability of lipid droplets as indenters, we first confirmed that they can be trapped and moved *in situ* and can cause a visible deformation of the nuclear envelope when they are moved against the nucleus with up to 200 pN of contact force (Fig. 4d). We next used purified droplets as rheological probes *in vitro* and embedded them into PA gels together with polystyrene microspheres as control probes. We then compared the G modulus of the PA gel obtained through TimSOM with both probes. The G modulus of PA gels measured with lipid droplets coincided with that obtained from embedded polystyrene microspheres (Extended Data Fig. 9). Together, this indicates that endogenous lipid droplets are sufficiently stiffer than other organelles, as also can be seen in their higher longitudinal modulus[85, 86], and are suitable as nano-indenters for optical force measurements.

We then trapped a single lipid droplet in the cytoplasm of intestinal epithelial cells and performed TimSOM from 0.5 Hz to 2 kHz. The rheological spectrum of day one adults fit well to a fractional Kelvin-Voigt model, indicative for the behaviour of a viscoelastic solid. Generally, the cytoplasm was 10 times stiffer than the cytoplasm of zebrafish progenitor cells (compare *C*_*α*_, *C*_*β*_; Fig. 3 with Extended Data Fig. 10, and Fig. 4 with Extended Data Fig. 5), which prevented us to move the droplet within the cytoplasm to probe multiple compartments with the same trapped object. This solid-like signature was significantly reduced in old animals (day eight adults), suggesting that viscoelastic properties of the cytoplasm change during age (Fig. 4e). In attempt to define the genetic factors that contribute to cytoplasmic ageing, we performed TimSOM in mutants known to display premature ageing phenotypes in *C. elegans*, such as mutants of the nuclear envelope[87].

We specifically tested a dominant negative *GFP::lmn-1*/Lamin A construct, and mutations in *emr-1* Emerin and *lem-2* LEMD2 whose human homologs are implicated in muscle waste disorders and progerias[88, 89, 90]. These proteins localize to the inner nuclear membrane and are involved in coupling heterochromatin to the nuclear envelope with proposed roles in nuclear mechanics[91, 92, 93]. Their role in regulating cytoplasmic rheology, however, is not understood. When we tested *GFP:lmn-1/Lamin A*, we found generally lower storage and loss moduli over all frequencies (Extended Data Fig 10a) in young adults, which was unchanged old animal. Fitting the fractional Kelvin-Voigt model to the data revealed a significantly reduced cytoplasmic viscosity (embodied in parameter *C*_*β*_ Fig. 4g) and a slight but insignificant increase in its elastic response visible in the reduction of the low-frequency exponent *α* (Extended Data Fig 10f). This suggests that wildtype Lamin function is required for cytoplasmic viscosity and that age may affect viscosity through changes of the nuclear envelope.

To further test the role of the nuclear envelope proteins in regulating cytoplasmic viscosity during age, we tested *lem-2* and *emr-1* mutant animals. To our surprise, we did not observe major differences in cytoplasmic viscosity *C*_*β*_ compared to control cells, neither in young nor old animals (Fig. 4e). Emerin *emr-1* mutants, however, had a significantly higher exponent *α* and *β*, indicative for a more fluid-like material (Extended Data Fig. 10f,g). Together, this suggests a general softening and fluidification of the cytoplasm due to the mutations of the nuclear envelope (Extended Data Fig. 10, Supplementary Text 3) and, our data show that mutations of the nuclear envelope sensitize the cytoplasm to age-related changes in viscoelasticity.

## 3 Discussion

Here, we introduced an active microrheology procedure based on optical gradient traps, optimized for small sample volumes that massively increases experimental throughput by reducing material expenditure, instrumentation complexity and measurement time. The direct, momentum-based force measurements enables these measurements in turbid media with varying refractive index and unknown probe morphology[94], which affords its application inside cells without time consuming calibration routines. Overall, these factors reduce costs and experimental complexities to provide a rich repertoire of parameters for drug screening and model testing.

Our method has significant advantages over existing active rheology routines based on two separate driving and detection lasers[27, 48] or separating one laser beam into two beams with orthogonal polarization[95]. Even though this would fulfil the condition that forces and deformations are measured instantaneously (for equations 1 to be valid without further compensation), this comes with the complexity that requires to align two optical paths and two detectors at the BFP interferometry collecting system. Further, the alignment of two laser beams is not constant over the whole field-of-view of the optical trap, limiting the execution of dynamic trapping experiments at various locations [27, 96] and the need of a strong trapping laser induces uncertainty in position detection[48]. Moreover, depolarization for light rays propagating at large angles would generate massive cross-talk between the two traps [97, 98], hence the measurements would be affected by depolarization in such a highly focused laser beam [97].

Despite the promised versatility, our approach has some short-comings, that arise from the violation of the time-continuity intrinsic in an AOD-based time-sharing configuration. Foremost, the response of the probe at high frequencies is underestimated if the elastic contribution of the material is too large. Thus, this deviation depends on the type of material that is being probed. We developed a detailed protocol and a compensation algorithm to correct for this deviation, which require basic assumption on the underlying material properties of the sample, e.g. if it behaves primarily like viscoelastic Maxwell liquid (such as biomolecular condensates[61, 19, 11]) or viscoelastic solid (Kelvin-Voigt). As shown here (Fig. 3 and Fig. 4), and by others[49, 9, 48, 47, 99] that cell primarily behave as a fractional Kelvin-Voigt material, which we found are less affected by this deviation. The time-sharing measurements cannot unequivocally determine the real-part of the response function in the frequency spectrum of pure and ideal Maxwell liquids in an arbitrarily stiff trap. Thus, it is important to make the measurements with a trap that is less rigid than the material behaviour at the Nyquist frequency. That is, one must choose a 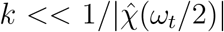 or equivalently *G*_0_ *<<* |*Ĝ*(*ω*_*t*_*/*2)|, such that it may be necessary to make several iterations to find the optimal stiffness of the trap *k* with which to make the measurements. Lastly, TimSOM, like other optical tweezer experiments inside cells requires a probe that can be manipulated and used to measure the mechanics[24]. They either need to be injected into cells or embryos[51, 43, 23], optoporated[100], phagocytosed[9, 15, 101] or biolistically delivered[102]. In samples where ectopic delivery of microsphere is not possible, protein aggregates[103] and cytoplasmic or nuclear lipid droplets[104, 53, 78, 80] and other internal structures[105, 106, 107] may conveniently be used to apply intracellular forces and measure mechanics.

Our examples illustrate how a single trap can be used to measure various materials, ranging from simple fluids to the complex response of viscoelastic fluid-like protein condensates and different organelles inside living cells and whole animals. TimSOM correctly measured the previously described Maxwell-like behaviour of biomolecular condensates and their time-dependent rigidity transition with a single trial. To do this, we used fractional rheology formalism in order to compensate the time-sharing response function to obtain the frequency-dependent G modulus of the condensates. Even though this rigidity transition can be identified by the fiber formation in 20% of all droplets[11], microrheology can determine the rigidity transition, even without any visible changes in droplet morphology. In contrast to the dual trap rheology assay, we should note that bulk rheology is independent of surface tension. Consequently, the low frequency range is not limited by surface effects and can thus be extended, in principle, to ultra-long timescales. We should point out at this point that the measured surface tension of the BMC is irrelevant, as the chemical environment inside a cell is different than that of the buffer *in vitro*. Thus, values for surface tension (*γ*) derived from dual trap experiments *in vitro* should be interpreted with caution. Likewise, the experimental routine is much simpler and does not require perfect alignment of the microspheres in the equatorial plane of the droplet. These important simplifications set the base for future implementations of high-throughput rheology measurements during research and drug discovery aimed to identify regulators of condensate ageing processes[108, 109].

A complete understanding of the adaptive response to mechanical stress requires knowledge of the cell cortex mechanics and ion channel dynamics[110] and the subcellular mechanical properties[111, 112]. We find that the mechanics of the nucleus is dominated by the cytoplasmic/nuclear interface rather than the nucleoplasm. This result coincides with recent observations made with Brillouin microscopy of cells in developing ascidian embryo, where a higher Brillouin shift was observed in perinuclear regions, indicative for a higher longitudinal modulus of the interface, than both, the nucleus and the cytoplasm[113]. Likewise, observations that the nucleus has a lower mass density and longitudinal modulus than the surrounding cytoplasm[114, 85] support our data that nucleoplasm is a soft, viscoelastic solid. This observation is consistent with the ‘open’ state of euchromatin of undifferentiated progenitor cells[115] which contrasts nucleoplasm rheology in differentiated cells [76]. Whereas the lamin nucleoskeleton did not seem to influence the complex shear modulus of the interface probed at small deformations, we found that Lamin A increased the resistance of the nucleus to large deformations applied our creep-compliance test. This is consistent with observations that Lamin B affects the stiffness and Lamin A the viscous resistance of the nuclear envelope[116]. Recently, it was shown that rheological measurements may be affected by the distance of the probe to the cell membrane, an effect that increases as the probe radius approaches the radius of the cell[117]. As the probe size is an order of magnitude smaller than the cell size, while displacements are small, we estimate that the wall effect remain negligible in our experimental system (Supplementary Text 5.4). Together, our results add to the previous observations that cells primarily respond as fractional Kelvin-Voigt materials[9, 48, 49]. This might have important consequences on the way how cells store and transmit mechanical energy over long time scales and distance, especially for morphodynamic events[12] and mechanosensation[118, 44].

Finally we investigated how cytoplasmic rheology changes with age in a living animal and provided evidence how mutations of the nuclear envelope affect the cytoplasmic rheology during age. The findings that the nucleus influences cytoplasmic stiffness is similar to glioma cells, in which the stiffness of the cytoplasm correlated with decreased level Lamin A/C[15]. The question arises how the envelope coordinates such cytoplasmic changes? It was proposed that the density of ribosomes induce crowding and scale cytoplasmic viscosity[119]. It is conceivable that age reduces the density of ribosomes as the synthesis demand of an animal decreases[120]. Alternatively, mitochondrial morphodynamics and number decrease during age[121, 122], posing a plausible explanation for the age-related decline in viscoelasticity and an exciting novel research direction of the involvement of mitochondria in cell mechanics. Future work needs to address how a change in cytoplasmic viscoelasticity affects cellular function during age and more general, how genetic factors influence rheological parameters and viscoelasticity.

## Acknowledgements

We would like to thank the PRBB fish facility for animal maintenance, *Caenorhabditis elegans* center (CGC, supported by the National Institutes of Health, P40 OD010440) for reagents and Neus Sanfeliu-Cerdan for help with sample preparation and comments on the first draft. All authors thank Arnau Farré, Erik Schäffer, the NMSB and Cell Dynamics lab members for discussion. We thank Peter Askjaer for *C. elegans* strains and the Single Molecule Biophotonics lab at ICFO for sharing chemicals.

## Funding

MK acknowledges financial support from the ERC (MechanoSystems, 715243), Human Frontiers Science Program (RGP021/2023), MCIN/ AEI/10.13039/501100011033/ FEDER “A way to make Europe” (PID2021-123812OB-I00, CNS2022-135906), “Severo Ochoa” program for Centres of Excellence in R&D (CEX2019-000910-S), from Fundació Privada Cellex, Fundació Mir-Puig, and from Generalitat de Catalunya through the CERCA and Research program. V.R. acknowledges financial support from the Ministerio de Ciencia y Innovacion through the Plan Nacional (PID2020-117011GB-I00) and funding from the European Union’s Horizon EIC-ESMEA Pathfinder program (101046620, BREAKDANCE 101072123). XS acknowledges funding from AGAUR (2017 SGR 324), MINECO (BIO2015-70092-R and PID2019-110198RB-I00), and the European Research Council (CONCERT, contract number 648201). CGC acknowledges a graduate fellowship from MINECO (PRE2018-084684). IRB Barcelona and ICFO are the recipient of a Severo Ochoa Award of Excellence from MINECO (Government of Spain).

## Author contributions statement

FCC performed *in silico* and experimental characterization of Tim-SOM, optical trapping simulations and biomolecular condensate microrheology with assistance from NSC and MF. SOV did experimental characterization of TimSOM, zebrafish microinjection, intracellular and intranuclear microrheology. CMF performed strain generation through CRISPR-Cas9, *in vitro* and *in vivo* analysis of *C. elegans* lipid droplets and intestinal microrheology. FP and SJ generated zebrafish strains and performed the cloning for mRNA used for microinjection. CGC established CPEB4 and MEC-2::UNC-89 co-condensates for *in vitro* characterizations of biomolecular condensates. XS, VR and MK were responsible for funding acquisition. PAF developed the theoretical compensation pipeline, performed simulations and experiments on model materials. FCC, PAF and MK conceived the idea and wrote the manuscript, with input from all authors.

## Data and Code availability

All numerical data is available in the source data document. Some data wa submitted to zenodo.org. The code to perform the TimSOM FDE simulation has been deposited to https://gitlab.icfo.net/rheo/Tweezers/timsom

## Competing Interest

PAF is holder of the patent #WO/2022/171898 protecting the time-shared optical tweezer microrheology technique[123]. XS is cofounder of Nuage Therapeutics. All other authors do not declare any conflicts of interests.

## 4 Extended Data Figures

**Extended Data Figure 1.**
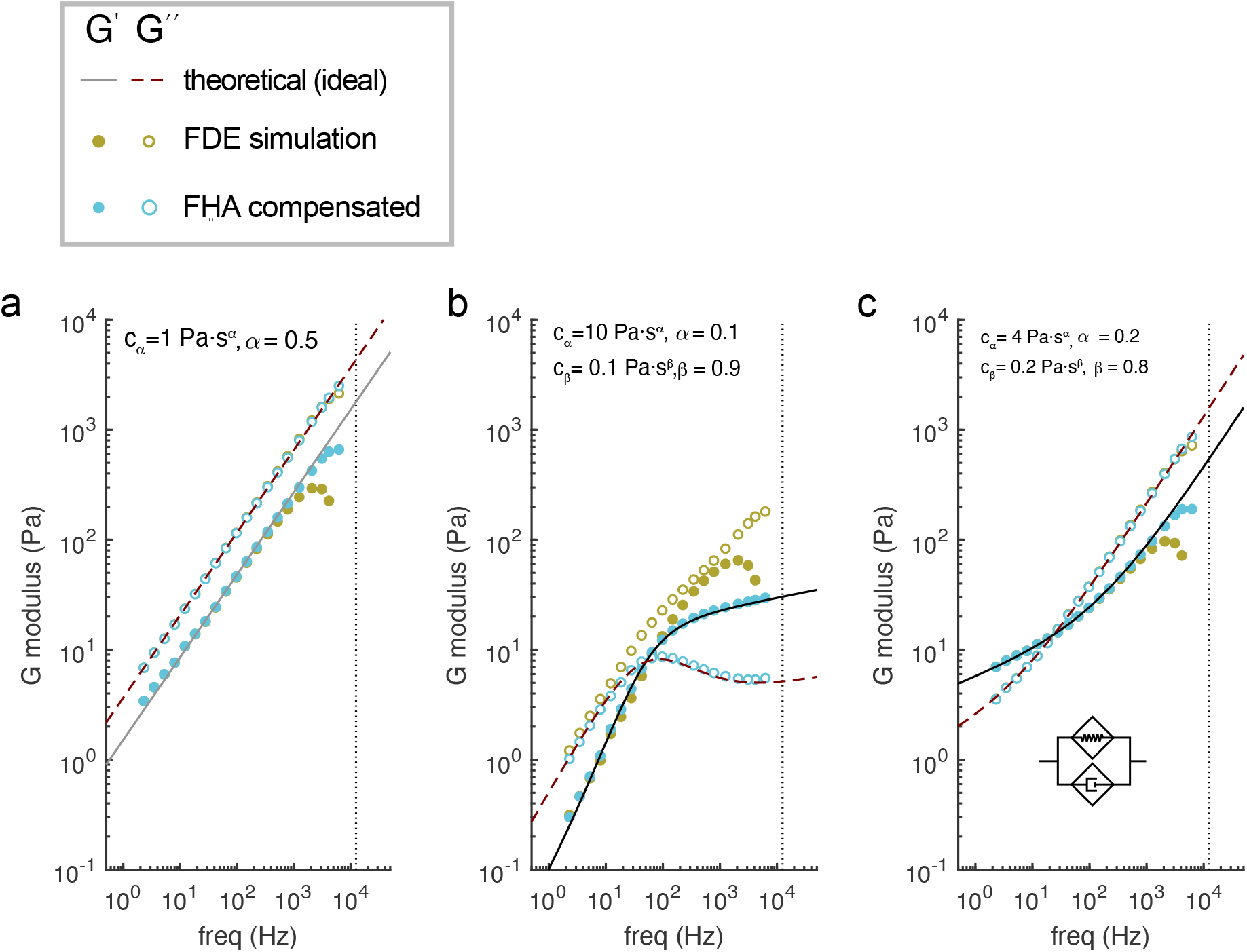
Theoretical analysis of the compensation of the G modulus extracted from TimSOM. **a-c**, G moduli calculated from the response function (Eq. 1), 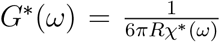, simulated through the FDE method for the (a) single springpot model, (b) the fractional Maxwell and the (c) fractional Kelvin-Voigt model. Parameters shown in the panels were used to during the FDE simulation.

**Extended Data Figure 2.**
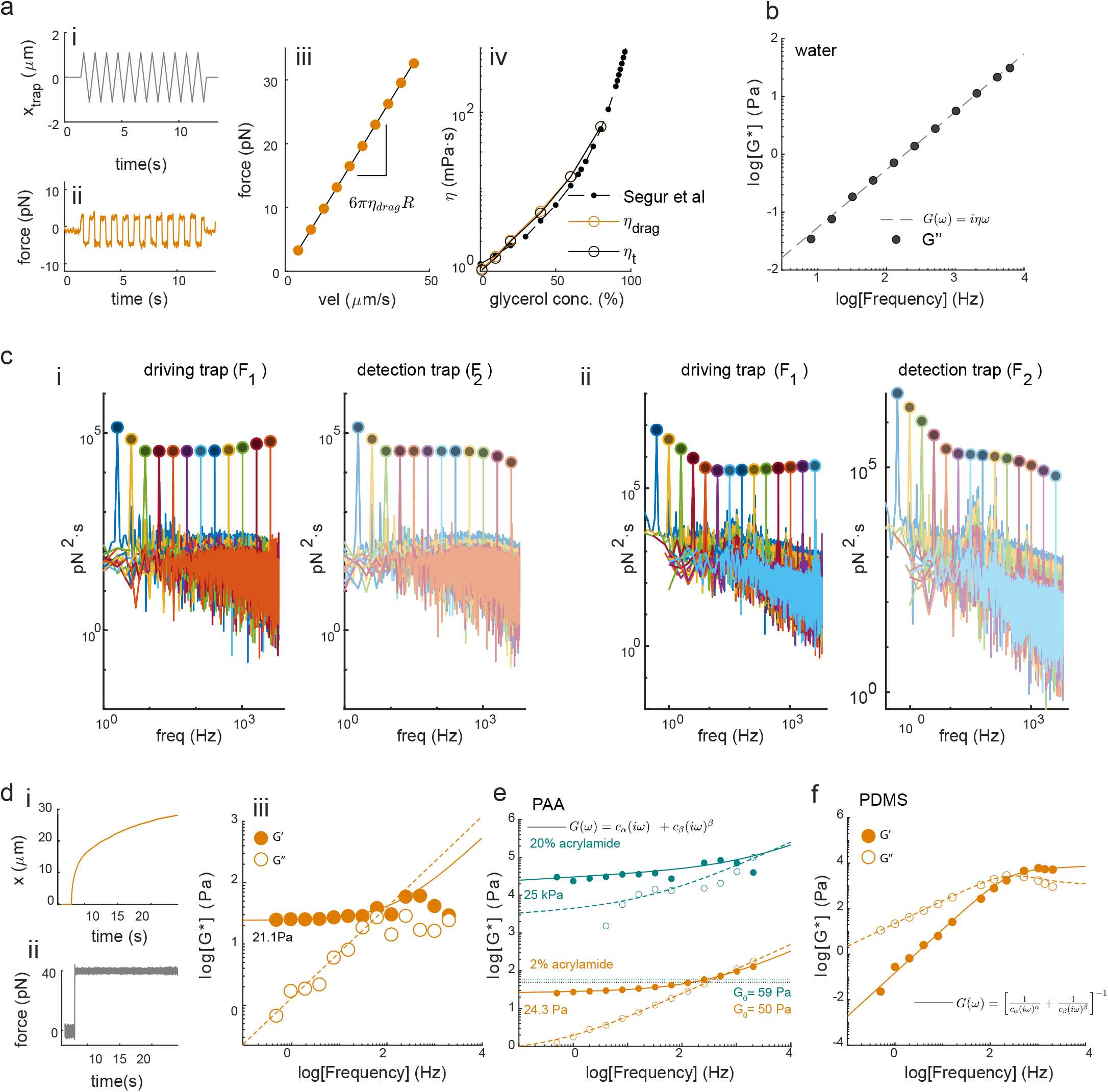
TimSOM accurately measures viscous and viscoelastic properties. **a**, Example of a triangular trajectory (i) used to generate constant drag forces (ii) to measure the viscosity of a purely viscous material. Viscosity was obtained from a linear fit (solid line, iii) to the averaged force values obtained for a series of triangles with different velocities (orange dots, iii). The drag force method, *η*_*drag*_, matched the values obtained with TimSOM, *η*^′^, with glycerol concentrations up to 80% (iv). **b**, Exemplary raw data of the G modulus obtained through Equation 1 which was compensated using Equation 46 in the Supplementary Text for water. **c**, Fourier and frequency domain analysis of TimSOM force signals. Power spectrum obtained from the force signals of the driving (*F*_1_) and static traps (*F*_2_) during an active microrheology measurement on a 1-μm microsphere in water (i, *f* = 2, …, 4096Hz) and in a 21.1 Pa PAA gel (ii *f* = 0.5, …, 4096Hz). The peaks are located at the imposed driving frequencies. The signal is clearly distinguishable and larger than the background noise. **d**, Force clamp experiment to retrieve the G modulus of a soft PAA gel. The creep motion of the trapped bead (*J*(*t*), i) upon a feedback-controlled constant force (*F*_0_ = 40 pN, ii) can be used to retrieve the frequency-dependent G modulus. Real (imaginary) part of the G modulus is plotted with filled (open) circles for the experimental retrieval of *G*(*ω*) and with solid (dashed) line for a fit using the Fractional Kelvin-Voigt model. Using the *J*(*t*) → *Ĝ*(*ω*) method: *C*_*α*_ = 21.1 ± 8.83 Pa (CI 95%). **e**, TimSOM measurements on two different gels with varying stiffness. The orange gel is the exact same one as in panel (d). Filled (open) circles correspond to the real (imaginary) part. The solid (dashed) lines are the real (imaginary) part of the fractional Kelvin-Voigt model as specified in the legends. For PAA gel, using the TimSOM method: *C*_*α*_ = 24.3 ± 1.28 Pa (CI 95%). The mint dots and lines correspond to a 20% acrylamide gel. The modulus is indicated in the figure, together with the trap stiffness *G*_0_ that was used to measure each gel. **f** Representative raw data of the G modulus for PDMS obtained through Equation 1,compensated using the FHA method (Equation 46, in Supplementary Text). Filled (open) circles correspond to the real (imaginary) part. The solid (dashed) lines are the real (imaginary) part of the model specified in the legends.

**Extended Data Figure 3.**
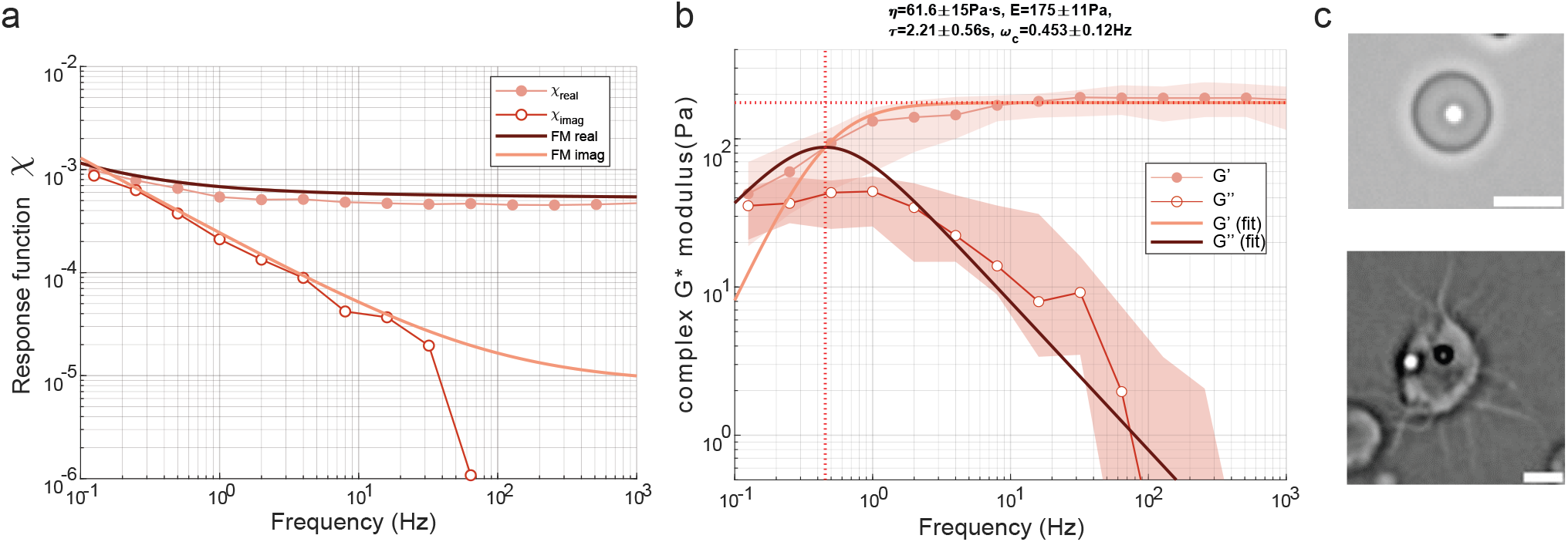
Rheology of CPEB4 droplets. **a** Response function acquired on naive CPEB4 condensates immediately after formation. Thick lines correspond to the fit of the data to a fractional Maxwell model **b** Rheological spectrum of CPEB4. Solid lines indicate the fit to a Maxwell model. G’, storage modulus; G”, loss modulus. Fit parameters are indicated on top of the graph: *η*, viscosity; E, plateau modulus; *τ*, relaxation time; *ω*_*c*_, crossover frequency. Mean ± standard deviation. **c** Representative micrograph of a fresh (top) and an ‘aged’ CPEB4 condensate (bottom) displaying emerging fibers indicating their solid transition. Scale bar = 5μm.

**Extended Data Figure 4.**
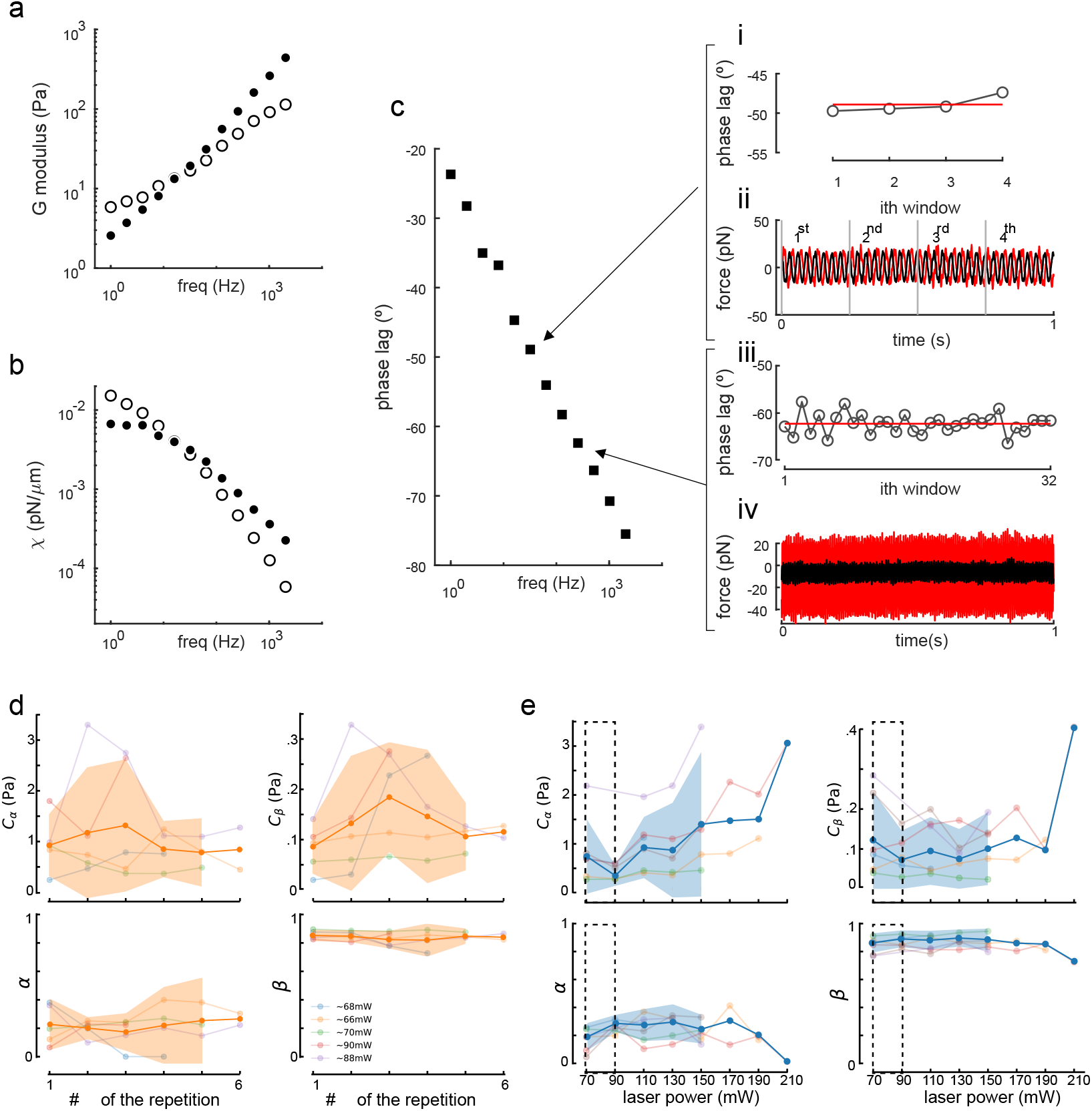
Performing the rheology routine does not induce phase lag difference in zebrafish cells. **a, b**, Representative measurement of the G modulus (a) and response function (b) on a microsphere in the cytoplasm of a zebrafish embryonic cell. **c**, Performing the rheology routine does not induce phase lag difference. Phase lag between the position and the force oscillatory signals. In inset i, the phase lag over consecutive time windows containing 8 cycles (inset ii) is shown for the oscillation at 32 Hz. In insets iii and iv, same as in i and ii, for the oscillation at 256 Hz. **d**, Performing rheological routines repeatedly in the same cell with the same power does not elicit a noticeable history effect. Each curve is acquired at the indicated laser power in the same cell. Solid curve = mean± 95% confidence interval. **e**, Performing rheological routines repeatedly with increasing laser power induces substantial stiffening in the cell, noticeable through an increase in *C*_*α*_. N=6 cells. Solid curve = mean± 95% confidence interval. The value of 90 mW in cell 1 was omitted as the fit to the fKV model did not converge. For clarity and convenience, we choose to start with 70 mW which provided enough trapping strength to resist cytoplasmic motion known as circus movement in these cells. The dotted box indicates the powers at which the measurements have been done throughout this manuscript.

**Extended Data Figure 5.**
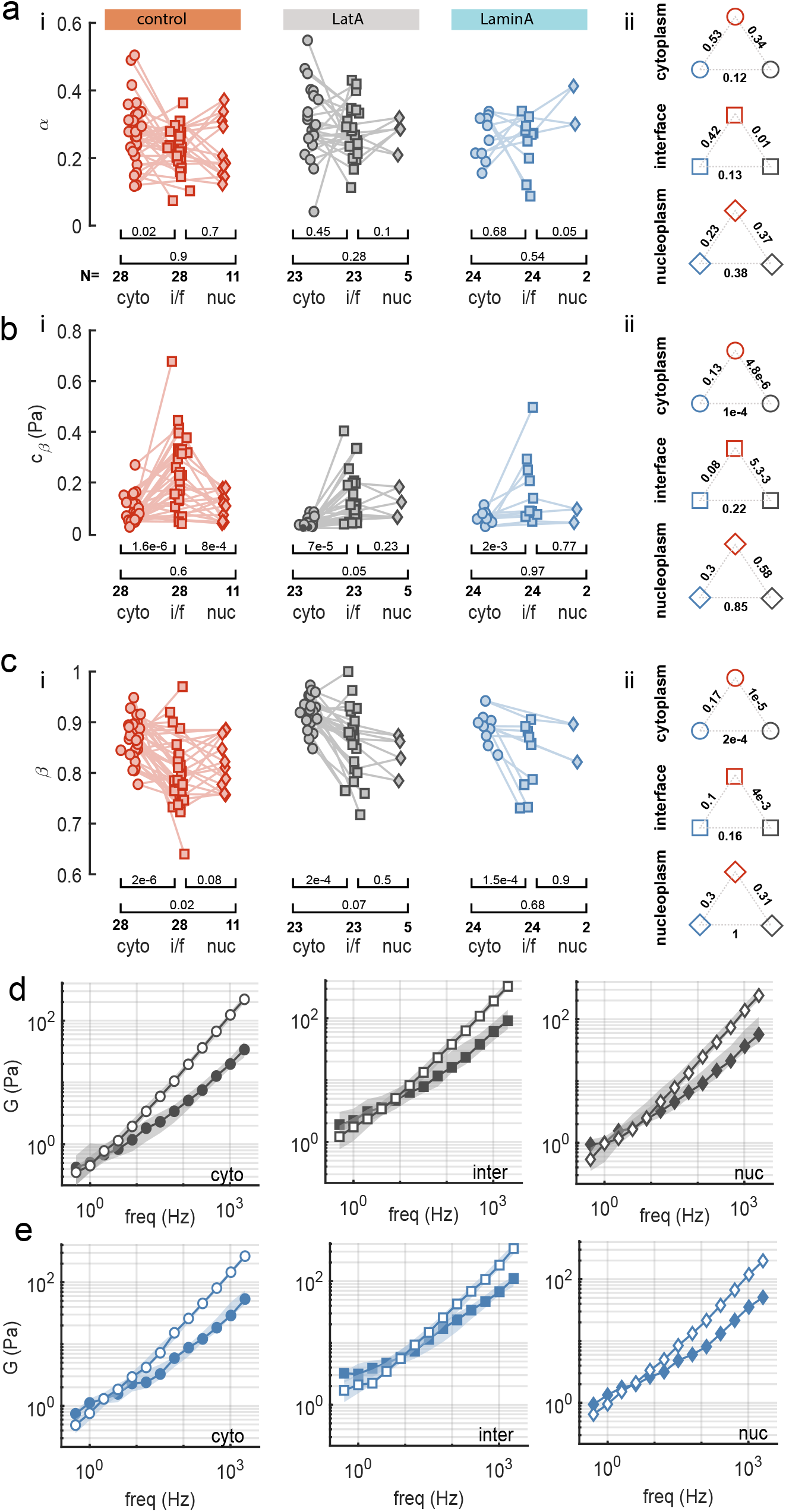
Active microrheology of zebrafish progenitor cells. **a**, (i) Exponent *α* of the low-frequency component derived from the fit of the fractional Kelvin-Voigt to the rheological spectrum in Fig 4. p-values below the graph derived from a paired t-test; N=number of cells measured. (ii) p-values derived from a pairwise, two-sided Mann-Whitney U-test of the indicated combinations. **b**, (i) Prefactor *C*_*β*_ of the high-frequency component derived from the fit of the fractional Kelvin-Voigt to the rheological spectrum in Fig 3. p-values below the graph derived from a paired t-test; N=number of cells measured. (ii) p-values derived from a pairwise, two-sided Mann-Whitney U-test of the indicated combinations. **c**, (i) Exponent *β* of the high-frequency component derived from the fit of the fractional Kelvin-Voigt to the rheological spectrum in Fig 3. p-values below the graph derived from a paired t-test; N=number of cells measured. (ii) p-values derived from a pairwise, two-sided Mann-Whitney U-test of the indicated combinations. **d, e**, Rheological spectrum derived from (d) latrunculin A treated cells and (e) Lamin A overexpressing cells

**Extended Data Figure 6.**
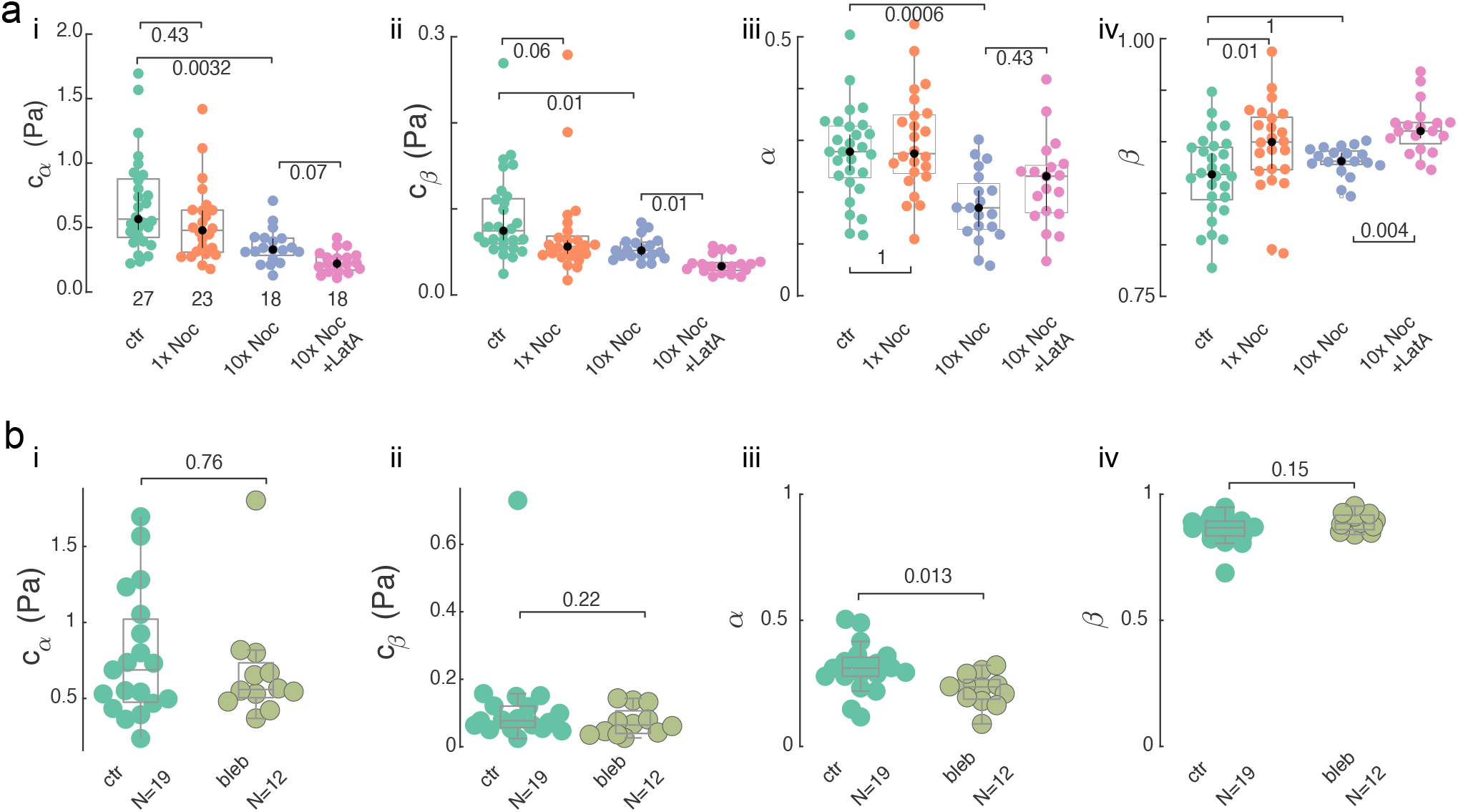
Cytoplasmic rheology is affected by tubulin organization but unaffected by myosin II activity. **a, b**, Dot plot of all four parameters extracted from a fit of the rheological spectrum to a fractional Kelvin-Voigt model comparing the cytoplasm of untreated control cells with cell treated with (a) varying concentrations of nocodazole (1μM and 10 μM) to depolymerize the microtubule cytoskeleton and combined LatA/Nocodazole to interfere with actin and microtubule network; and (b) 10 μM blebbistatin to inhibit myosin II contractility. (i) Prefactor *C*_*α*_ indicating magnitude of the low-frequency, elastic response; (ii) Prefactor *C*_*β*_ indicating magnitude of the high-frequency, viscous response; (iii) Exponent *α* indicating the low-frequency, solid behavior; (iv)) Exponent *β* indicating the high-frequency, fluid behavior. Values close to the horizontal bracket indicate the p-value of (a) Kruskal-Wallis test with a Dunn post-hoc test for pairwise comparisons between groups with Bonferroni adjustment, and (b) unpaired t-test.

**Extended Data Figure 7.**
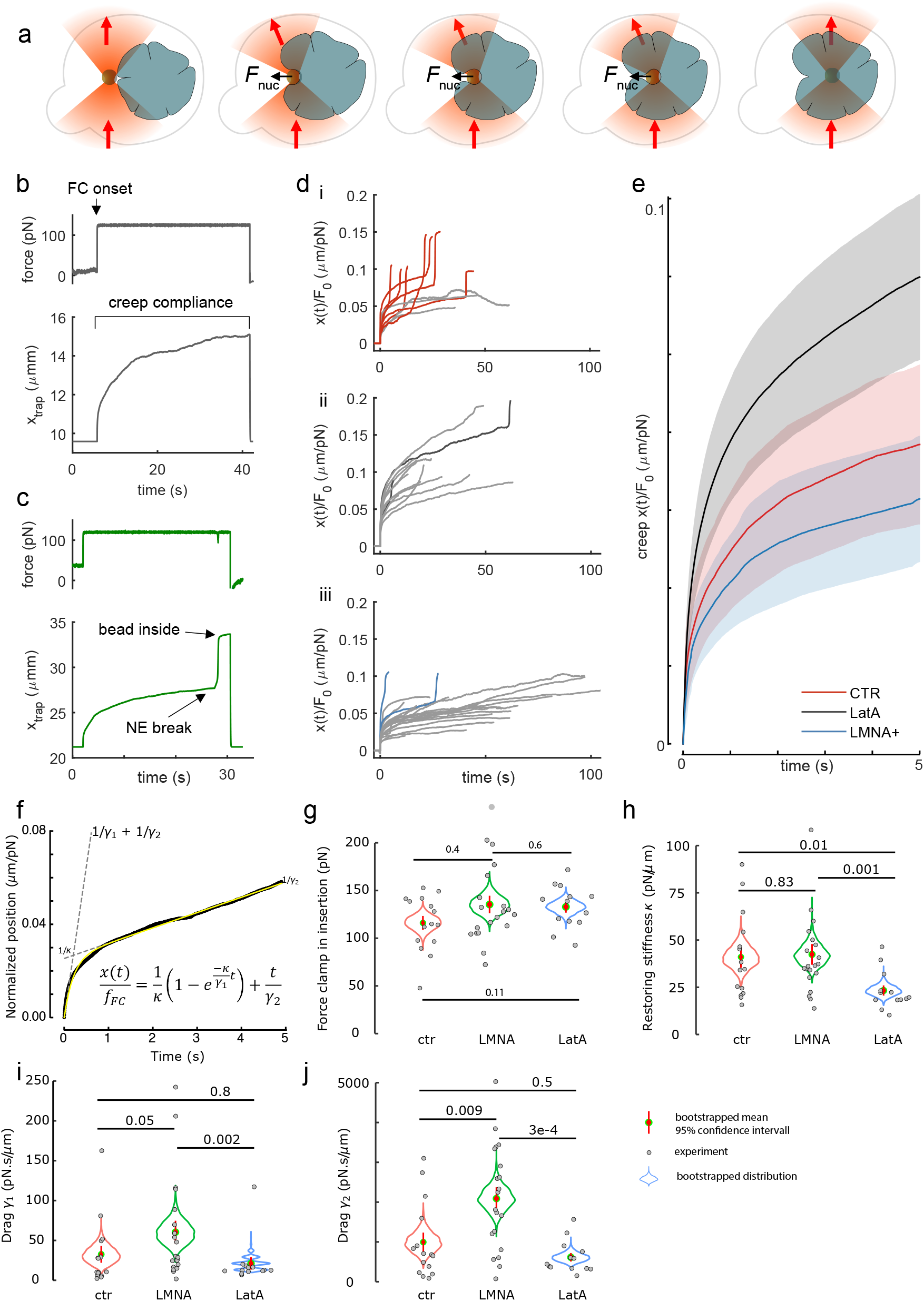
Force clamp protocol for microsphere insertion of into zebrafish embryonic cell nuclei. **a**, Schematics of the nuclear insertion process using the force clamp modality of the optical micromanipulation platform. A bead is brought into contact with the nucleus and a force clamp is set at *F*_0_ ∼ 100 − 150 pN. During some tens of seconds, the bead creeps against the nuclear envelope, eventually breaking it and entering the nucleus. **b**, Example of an unsuccessful bead insertion assay. The nuclear envelope is too stiff for the optical trap to enter the nucleus. At *t* ∼ 5 s, the force is set to 120 pN (top) and the trap pushes the bead against the nucleus. After *t* ∼ 40 s, the force clamp is turned off. **c**, Example of a successful insertion. Similar to (b), a force clamp of *F*_0_ = 120 pN is initiated at *t* ∼ 5 s. Around *t* ∼ 25 s, the nuclear envelope breaks and the trap accelerates into the nucleus. After this, the force clamp is turned off. **d**, Summary for control cells (i), cells incubated with latrunculin A (ii) and cells with lamin A overexpression (iii). Trajectories have been normalized with respect to the force setpoint as *x*(*t*)*/F*_0_. Colored traces (red: CTR; black: LatA; blue: LmnA+) indicate successful events in which the bead gets inserted into the nucleus. Gray lines are force compliance curves that didn’t succeed in inserting the bead into the nucleus. **e**, Normalized creep compliance curves over the first 5 seconds of creep test. Solid lines are medians and shaded areas correspond to the ±25% quantiles. Note, all curves show a fast initial compliance, followed by an inflection indicating an elastic plateau. **f**, Representative data showing the fitting procedure. Yellow line indicates the fit, dotted lines indicate the relevance of the extracted parameters. **g**, Quantification of the force applied to the nucleus during the constant force experiment. **h-j** Leading parameters extracted from the fit. (h) Restoring stiffness of the nuclear envelope; (i) resistance to deformation on short time scales and (j) resistance to deformation on long time scales. Numbers above the bracket indicate the p-value derived from two-tailed pairwise comparison using a Mann-Whitney U-test on the experimental data points (grey). Violin plots represent the distribution of the bootstrapped mean value (±95% confidence interval) calculated from 5000 virtual experiments, using the experimental data as a sample to describe the population.

**Extended Data Figure 8.**
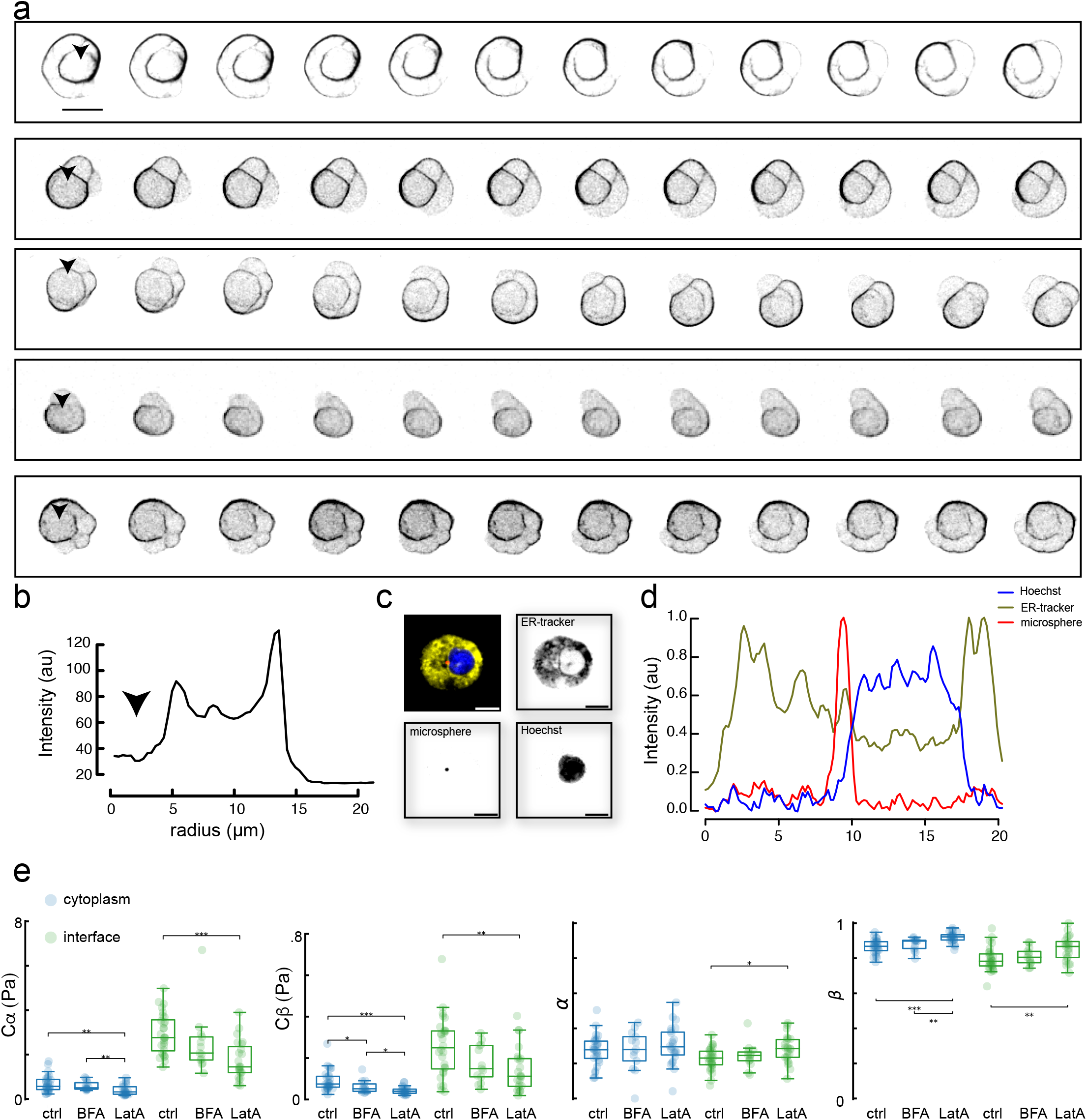
Actin distribution in isolated progenitor cells undergoing blebbing behavior. **a**, Montage of five different isolated mesendoderm progenitor cells, showing dynamics of filamentous actin stained with lifeact-GFP. Arrowhead indicates the location where the nucleus is expected. Note, how the ‘old’ cortex collapses onto the cell center and accumulates around the nucleus. Length of each sequence, t=120s. Scalebar=10 μm. Embryos were injected with 50pg lifeact-GFP to visualize filamentous actin and were isolated in sphere/dome stage according to the methods described in this manuscript. Data set from ref. [73]. **b**, Radial profile plot of the averaged intensity as a distance from the cell center. Arrowhead points to the nuclear location. Representative for N=10 cells. **c**, Representative image of a zebrafish cell stained with ER-tracker to label the endoplasmatic reticulum, Hoechst to label the nucleus to highlight the location of the bead with respect to these organelles. Scale bars=10*μ*m. **d**, Radial profile plot showing the intensity distribution of the three channels. Note, the microsphere is in contact with the nucleus, without large accumulation of ER in between, as judged from the absence of the green ER signal. **e**, Parameters extracted from the microrheology routine on zebrafish cells treated with 5 *μ*g/mL BrefeldinA (BFA) and LatA for comparison to the data in Fig. 3 and Extended Data Figure 5. Only significant p-values are indicated in the combinations, derived from a two-sided U-test.

**Extended Data Figure 9.**
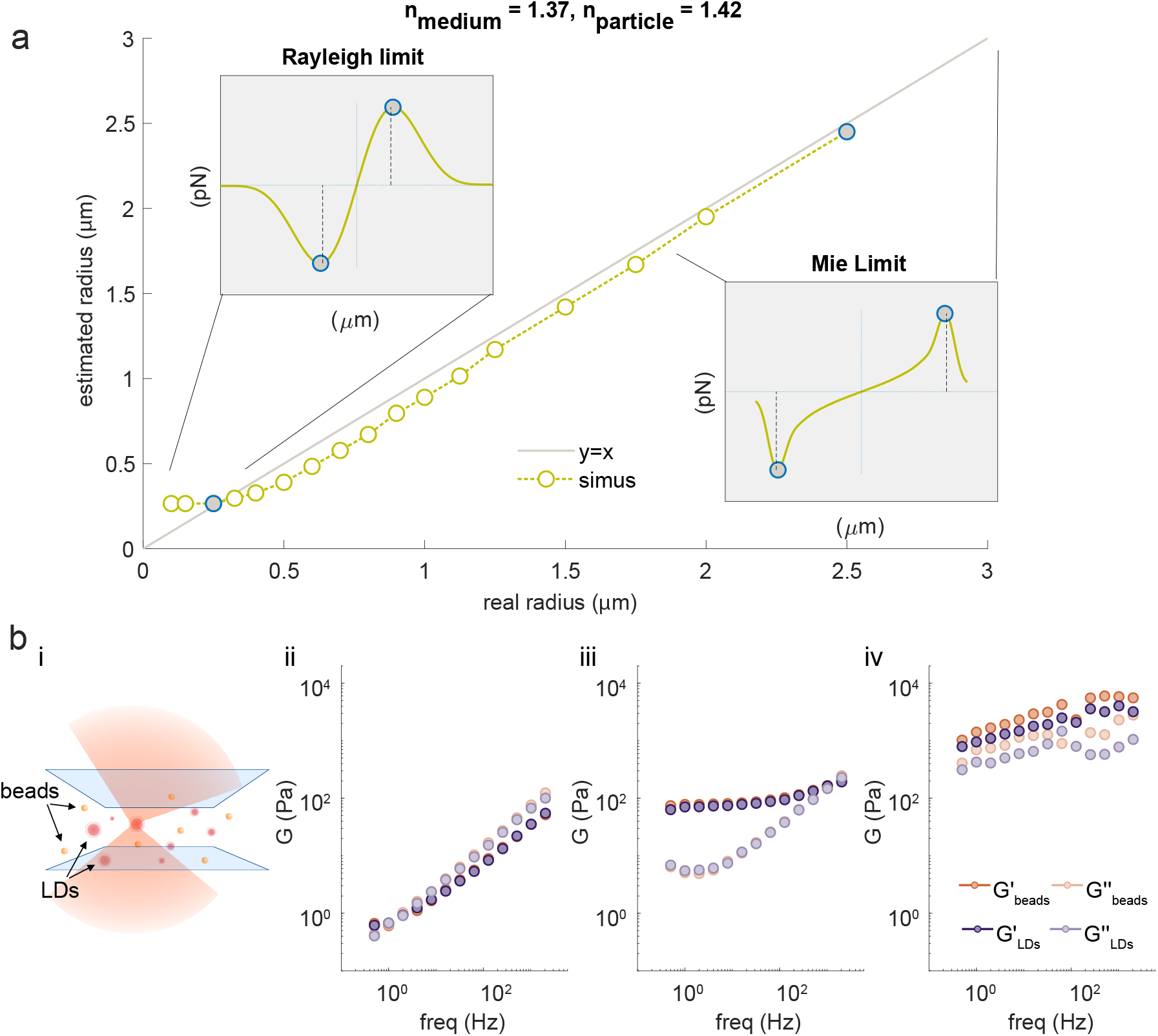
Lipid droplets correctly measure rheological properties up to several kPa. **a**, Estimation of the LD size using simulations with the Optical Tweezers Toolbox [84]. The refractive index of the lipid droplet was considered *n*_LD_ = 1.42 and that of the tissue *n*_worm_ = 1.37. The numerical aperture of the laser trapping beam was approximated to 1 as the entrance pupil of our NA=1.2 trapping objective is underfilled with a factor of 0.8. We considered radii from *R* = 0.1 μm to *R* = 2.5 μm. From the force profiles, we could measure the peak-to-peak radius, and finally obtained a look-up table from which the *real radius* is obtained from the *estimated radius* from the force scan across the LD. Insets i and ii show examples in the Rayleigh and Mie regimes, respectively. While the radius of probes in the strict Rayleigh regime degenerates and can’t be addressed, the estimated radius tends to the real radius in the Mie regime (geometrical optics). **b**, (i) Schematic of the experiment. Lipid droplets are isolated from *C. elegans* and embedded into a poly-acrylamide gel with varying crosslinker (bis-acrylamide) concentration. (ii-iv) Comparison of the rheological spectrum performed with the lipid droplets and the polystyrene microspheres, embedded into a gel with ii) 2-10 Pa, iii) 10-100 and iv) 100-1000 Pa elastic modulus.

**Extended Data Figure 10.**
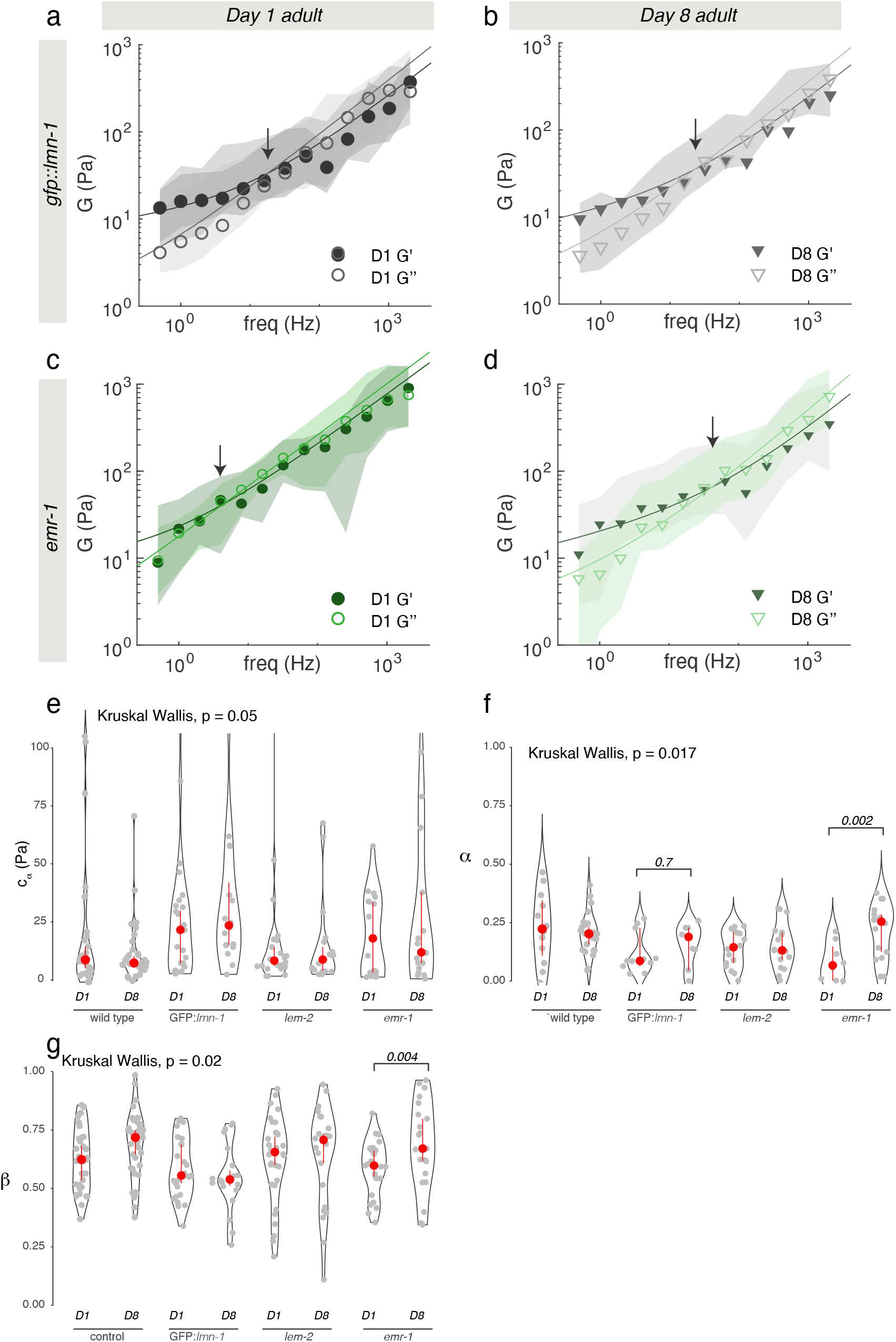
Mutations in nuclear envelope sensitize the cytoplasm of *C. elegans* to age-related changes in viscoelasticity. **a, b**, Rheological spectrum derived from dominant-negative *gfp:lmn-1* expressing animals at (a) day 1 and (b) day 8 adulthood. **c, d**, Rheological spectrum derived from *emr-1* mutant animals at (a) day 1 and (b) day 8 adulthood.The median and ±25%quantiles are represented by lines and shadows, respectively. **e**, Prefactor *C*_*α*_ of the low-frequency component derived from the fit of the fractional Kelvin-Voigt model to the rheological spectrum in Fig 5e,f and panels a-c above. **f**, Exponent *α* of the low-frequency component derived from the fit of the fractional Kelvin-Voigt model to the frequency spectrum. **g**, Exponent *β* of the high-frequency component derived from the fit of the fractional Kelvin-Voigt model to the frequency spectrum. Red circle indicates median±bootstrapped 95% confidence interval. For statistical comparison and p-values see Supplementary Table 2.

## 5 Materials and Methods

### 5.1 Theory and simulation of the trap displacement in a viscoelastic medium in a time varying trapping potential

The effect of having two alternating optical traps on the actual measurement of the response function, *χ*(*ω*), and thereby the G modulus, *G*(*ω*), is introduced in this section and detailed in the Supplementary Information. Two methods are introduced. First, we use a numerical method in the time domain, aiming at the trajectory of the trapped probe, in order to emulate the violation of the continuity condition between the force *F*_*T*_ (*t*) = *F*_1_(*t*) + *F*_2_(*t*) and position *x*_*p*_ = *F*_2_*/k* measurements (Figs.1). Second, by truncating the dynamic, oscillatory trapping potential at the first harmonic, we obtain an analytical expression in the frequency domain that we further use to compensate the time-sharing effect.

#### Study of the deviation via Fractional Differential Equations

Under an AOD-modulated time-sharing regime, the laser spot gradually vanishes from trap 1 and appears into trap 2 as the acoustic wave enters the laser beam cross section at the AOD crystal (see Supplementary Information). In our set-up, this occurs within a transition time of *τ* = 10 μs. After that, the laser spot remains at trap 2 for 30 μs and the transition is thereby reversed back into trap 1. Meanwhile, a voltage datum is sampled with a delay of 33 μs with respect to the rising edge of the acoustic wave. Because traps 1 and 2 fall within the linear regime of the trapped particle, forces acting onto it are equivalent to that from a single trap undergoing the following trapezoidal trajectory (see Supplementary Information):

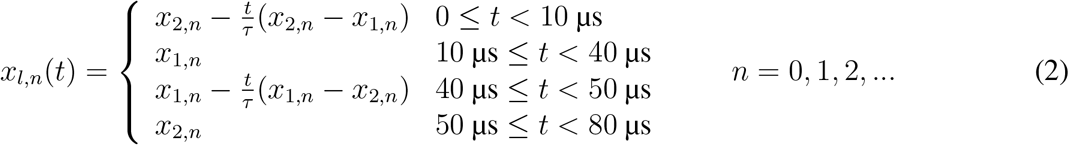

where *x*_2,*n*_ = 0, ∀*n* and *x*_1,*n*_ are discrete values of *x*_1_(*t*) = 2*x*_0_ sin(*ωt*) sampled at *f*_*t*_ = 12.5 Hz. Here, the amplitude of the oscillation in *x*_1_(*t*) is expressed as *A* = 2*x*_0_ for mathematical convenience. In order to capture the power law rheological behavior of most samples in biology, we use the fractional time-derivative operator, 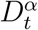, which describes the stress-strain relationship of a springpot, 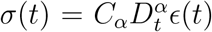 [47]. This element represents, continuously, all viscoelastic responses between the purely elastic unit —the *spring*, with *σ*(*t*) = *κϵ*(*t*)— and the purely viscous unit —the *dashpot*, with 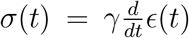. We numerically solved the equations detailed in Table 1 to obtain the trajectory of the probe bead in the lab frame, *x*(*t*), at different driving frequencies, *ω*_*j*_. The instantaneous force is thereby obtained as *F* (*t*) = *k*(*x*_*l*_(*t*) − *x*(*t*)), from which the interleaved BFP interferometry voltage signals, *V*_1_(*t*_1,*i*_) and *V*_2_(*t*_1,*i*_), are sampled at time points *t*_1,*i*_ and *t*_2,*i*_, respectively. Finally, the fast Fourier transform of the two signals is taken and the response function, 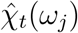, and G modulus, *Ĝ*_*t*_(*ω*_*j*_), obtained using Eqs. 1. We developed a package in Python to carry out the FDE simulations (see more details in the Suppl. Information). The comparison between the physical response function, 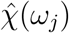, and the one accounting for the time-sharing deviation, 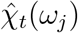, is shown in Fig. 1d for three different power-law materials. The corresponding G moduli (both the physical and the time-sharing deviated) are shown Extended Data Fig. 1.

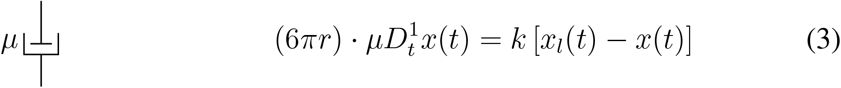

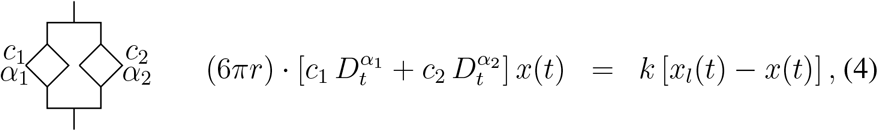

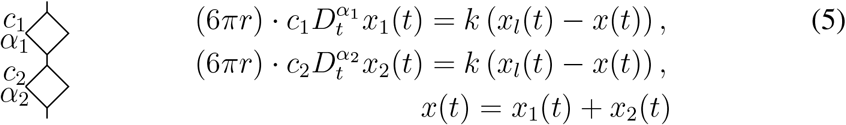

**Table 1.** Fractional-differential equations for a purely viscous dashpot (top), fractional Kelvin-Voigt (middle) and fractional Maxwell (bottom) models.

### 5.2 The First Harmonic Approximation (FHA)

The second approach to the study of deviation is based on analytically solving the equations of motion of the bead 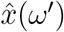 in the space of frequencies *ω*^′^. The equation to be solved is the algebraic equation

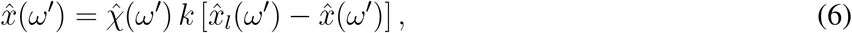

which becomes

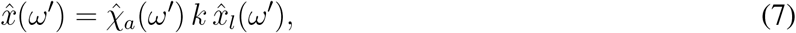

Here, 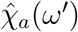 is the the response function of the active-passive system (see Supplementary Materials). 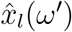 is the Fourier transform (FT) of the First Harmonic Approximation (FHA) of the trap trajectory, which in the time domain *t* reads:

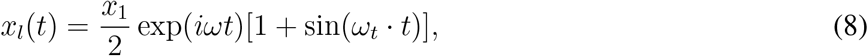

where the complex oscillation exp(*iωt*) is used, instead of a real one such as sin(*ωt*) or cos(*ωt*), with the sole purpose of simplifying the notation in the frequency space. After FT, the trap trajectory reads

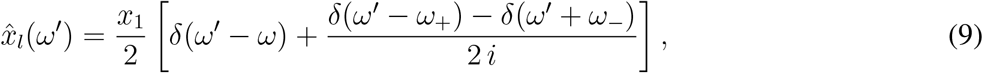

with *ω*_±_ = *ω*_*t*_ ± *ω* and where *δ*(*ω*^′^) is the Dirac distribution.

To get the linear response function *χ*_*t*_(*ω*) from the FHA approximation, the same steps identified for the FDE simulation must be followed, but this time using an analytical approach. The details of the calculations are given in the Supplementary Material. The obtained expression of 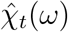 is a function of the response function of the material 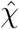, of the time sharing frequency *ω*_*t*_, of the trap stiffness *k* and reads

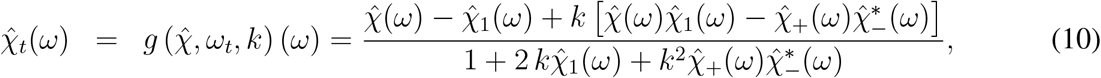

where

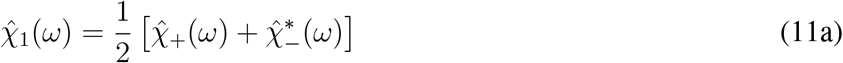

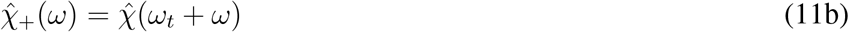

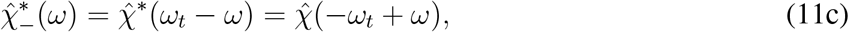

and the asterisk ^∗^ denotes the complex conjugate operation.

The results of the Eqs. 10 and 11 show that 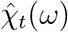 depends on the response function 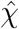 measured at different frequencies. In addition to the contribution at the excitation frequency *ω*, the contributions from the sidebands of the amplitude-modulated signal at the frequencies *ω*_−_ = *ω*_*t*_ − *ω* and *ω*_+_ = *ω*_*t*_ + *ω* also determine the behavior of 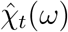.

Before using Eq. 10 to understand the causes of the deviation, the goodness of the FHA approximation should be checked by comparing the results predicted using Eq. 10 with those predicted using the FDE approach. To do so, we applied the FHA correction to the deviated data simulated through FDE 1d.

### 5.3 Optical micromanipulation and fluorescence microscopy

#### Optical tweezers

Our optical micromanipulation and microscopy platform is built around an inverted microscope (Nikon Ti-2) and has been used in previous publications and described extensively elsewhere [51]. A commercially available optical tweezers unit (SENSOCELL, Impetux Optics) is coupled to the rear epifluorescence port of an inverted Ti2 Nikon microscope. The trapping laser is modulated by a pair of acousto-optic deflectors (AODs, for *x* and *y* axis control) optically-conjugated to the entrance pupil of a water-immersion objective (Nikon Plan Apo, 60x NA=1.2), which in turn focuses the trapping beam onto the focal plane thus generating the optical traps. A direct force sensor substituting the bright-field illumination condenser captures the light from the optical traps, that has interacted with the sample, to detect transverse light momentum changes and thus measure the force directly[124]. Between the trapping objective and the optical force sensor, samples are mounted with two glasses as detailed below for each type of experiment.

#### Force and position detection with time-sharing optical tweezers

Optical force measurements were carried out through capture of trapping light momentum changes, which enables measurement of trapping forces on irregular samples, as well as in non-viscous media, with a sample-independent force measurements [124]. Briefly, light captured with a BFP interferometry system, optimized for light momentum detection (SENSOCELL, Impetux Optics), is conveyed to a position-sensitive detector (PSD), which volt-to-pN conversion factor, *α*, is calibrated by the manufacturer. The same system can be used to measure probe positions —relative to the trap—, after measurements of the trapping stiffness, *k* (pN/μm), which is performed by fast scanning the trap across the probe. Variations in the initial light momentum and trap power over the field of view are compensated through the driving software of the optical traps (LightAce, Impetux Optics). A detailed protocol for the start-up and use of this optical tweezers platform can be found elsewhere [51].

#### Active microrheology

Our active microrheology measurements consist of four steps:

- Measuring the initial light momentum variation around the trapped probe. This is only needed for solid samples that require trap/probe centering. Conveniently, for primarily liquid samples (water, protein droplets) the bead is pulled into the trapping potential without any centering routine.
- Measure the trap stiffness. To do this, we scan the laser across the microsphere, or lipid droplets as in experiments concerning the live animals, and fit a line to the linear regime, from -200 nm ≤ 0 ≥ 200 nm [24]. The slope of the line directly provides *k* through a link between force (as measured through momentum changes on the PSD[52]) and particle position such that 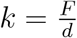.
- Apply a series of oscillations with specific parameters of frequency, amplitude and measurement duration. The frequencies and amplitudes applied in every measurement are specified in Supplementary Tables S3-5
- Choice of *G*_0_
- Retrieval of the 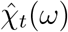 and *G*(*ω*) values through the First Harmonic Approximation method.

The experimental scheme detailed in Section 5.1 is implemented using the manufacturer’s AOD-addressing software (LightAce, Impetux Optics) and analysed through an executable interface (RheoAnalysis, Impetux Optics), which includes the TimSOM compensation pipeline, 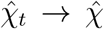 thoroughly described in the Supplementary Text 1.1.12. The AODs are addressed at a frequency of 25 kHz, for which, in the dual-trap time-sharing configuration, each trap is addressed at 12.5 kHz, hence the maximum oscillatory frequency for active microrheology results 6.25 kHz (Nyquist frequency).

#### Spinning-disk confocal microscopy

A multimodal spinning-disk confocal unit (Andor Dragonfly 505, Oxford Instruments) is coupled to the left port of the microscope. It is composed of seven excitation lasers stabilized through a wavefront filter (Borealis, Oxford Instruments) and the corresponding dichroic optics for multiple beam combinations. For GFP and mCherry parallel imaging, the 488 nm and 561 nm laser wavelengths were chosen, were directed to the sample through a quad-band dichroic mirror (405 nm, 488 nm, 561 nm, 640 nm) and a spinning disk with an array of 40 μm diameter pinholes rotating at 6000 rpm. The fluorescence emission signal was collected by the same objective, sent back to the confocal unit and directed to the cameras through the dichroic mirror. A long-pass dichroic mirror (cutoff wavelength: 565 nm) split the red and green signals, which were respectively imaged by two sCMOS cameras (Andor Sona, Oxford Instruments). For Hoechst 33342 and GFP parallel imaging, the 405 nm and 488 nm laser wavelengths and the same quad-band dichroic mirror were chosen. Fluorescence emission was further split with a 500 nm cutoff wavelength long-pass dichroic mirror.

#### Microrheology in calibrated materials

##### Glycerol solutions

Different concentrations of glycerol mixed with fluorescent bead solution (1 μm, Carboxylate-Modified microspheres, Thermofisher) were prepared to calibrate the TimSOM technique in a viscous material. The concentrations ranged from 0% (MilliQ water) to 95% glycerol and were prepared gently avoiding the formation of bubbles from a 99% glycerol stock solution. Since the material is purely viscous, the trapped microsphere will relax into the trap focus, ensuring alignment with no need of a probe/trap centering routine (see below). We obtained the viscosity, *η*_*T imSOM*_, for the glycerol mixtures from the slope of the loss modulus *G*^′′^(*ω*) = *η*_*T imSOM*_ *ω* (Extended Data 2b).

To benchmark the results, we performed Stokes-drag force measurements, on the same bead, after the time-sharing microrheology routine. We applied triangular waveforms with increasing velocities, *v*_trap_, until the microsphere escaped the optical trap (Extended Data Fig. 2a). The plateau force during the bead movement, *F*_drag_, increased with velocity, indicative for a purely viscous liquid. We then estimated the viscosity, *η*_*drag*_, by fitting *F*_drag_(*v*_trap_) = (6*πη*_drag_*Rv*_trap_) to the force-velocity plots, where *R* is the radius of the trapped microsphere (Extended Data Figure 2a ii). To avoid changes in viscosity due to laser absorption, trapping power was left below *P* =65 mW [125]. To avoid hydrodynamic interaction with the chamber surfaces, the trapping plane was kept at a relative height of *z* = 20 μm [126].

##### Polyacrylamide gels

Soft PAA gels were polymerized from a modification to the recipe used in [127]. A gel premix (GPM) made of 30% acrylamide (Sigma A8887) and 0.8% bisacrylamide solution (Sigma 66670) is mixed with 5μL of fluorescent microsphere solution at stock concentration (2% solids, F8816, Thermofisher) and diluted in water or a solution of isolated lipid droplets for lipid droplet characterization (Extended Data Fig. 9). Exact quantities for obtaining specific gel stiffness are tabulated in 2. GPM containing microspheres was thoroughly resuspended and later mixed with 5 μL of amonium persulfate (APS, Sigma A3678, 10% diluted in PBS) and 1.5 μL TEMED (Sigma T9281) for the polymerization process. Prior to gelling, 40 μL of the solution were transferred into the optical trapping cavity. The cavity was then closed with a #1.5 cover glass as described above[51]. Measurements were performed after 30 minutes of the sample preparation in order to have the gel homogeneously polymerized.

Because the gels are a viscoelastic solid material, in general, the embedded probe will not relax into the center of the optical trap once it is placed on it. A probe/trap centering routine, followed by the trap stiffness measurements, was therefore carried out before the microrheology measurement over the frequency series. Similar to previous reports, we modeled the rheological spectrum as a fractional Kelvin Voigt material to extract the relevant parameters [55].

To estimate the limits of the detection in the TimSOM configuration and therefore the maximum stiffness one can measure, we prepared a 30kPa PAA gel. We performed rheology routine at maximum laser power (300mW) and Fourier transformed the force and the displacement signal derived from the active and the passive trap.

##### Creep Compliance measurements

An independent measurement of the G modulus of polyacrylamide gel was obtained through measurement of the creep compliance upon the application of a step in stress, using LightAce’s force clamp utility at a given setpoint force. We used a trap stiffness of 890pN /*μ*m and 220*μ*N/*μ*m for stiff and soft gels respectively. To do so, we set the proportional gain to Kp=0.001, 0.004*μ*m/ pN, the integral gain to Ki=1.1, 4.5 *μ*m/ pN. s and the derivative gain to Kd=0 for a stiff and a soft gel respectively. With these parameters we set a constant force of *F*_0_ = 40 pN, which leads to a typical viscoelastic compliance curve, *x*(*t*). We thereby used the mathematical model derived by Evans et al. [54], implemented in custom Matlab code, to calculate the frequency-dependent shear modulus from measurements of *J*(*t*) in the time domain. For two gels with different stiffness, results are plotted in Supp. Data Fig. 2c, d. The position of the trapped probe was calculated as *x*_bead_ = *x*_trap_ − *F/k*_OT_. The low-frequency plateau modulus was calculated from the frequency-dependent storage G-modulus after fitting to a fractional Kelvin Voigt model and indicated in the Extended Data Fig. 2c, d.

##### PDMS

Poly-dimethyl-siloxane prepolymer was mixed with a curing agent at 100:1 and mixed thoroughly with *d*=1 μm microspheres (F8816, Thermofisher). The solution was degassed in a vacuum pump to eliminate bubbles introduced during the mixing. The mixture was used immediately for rheology to avoid long-term curation of the silicone. Due to the high viscosity of the solution, the bead did not relax into the trap in a time short enough for experiment viability, hence the probe/trap centering routine was used.

#### Zebrafish experiments

##### Husbandry

Wild-type embryos were obtained from the AB strain background and actin was visualized with a stable transgenic line (Tg(actb2:Lifeact-GFP)) [128]. Zebrafish embryos were kept in E3 medium at 25° to 31 °C before experiments and their stages were identified according to hours post-fertilization (hpf) and morphological criteria [129]. All protocols used have been approved by the Institutional Animal Care and Use Ethic Committee (PRBB–IACUEC) and implemented according to national and European regulations. All experiments were carried out in accordance with the principles of the 3Rs (replacement, reduction, and refinement).

##### Microsphere injection and cell preparation

Zebrafish embryos were injected 1 nL of microspheres (F8816, Thermofisher, diameter *d*=1μm) at 1:5 of the stock solution at the one-cell zygote stage in order to ensure proper distribution of microspheres, as described elsewhere [51]. When the embryos reached sphere stage (approximately at 4 hpf), they were dechorionated with a pair of forceps and their cells were manually dissociated and left for at least 30 min to recover[130]. To visualize the nucleus, dissociated cells were incubated for six minutes in DNA-Hoechst at 1 μg/mL final concentration following the protocol in [51] A layer of double scotch tape (approximately 20 × 20 mm wide)
 was used as a spacer between the lower and the upper surfaces. A 1×1 cm hole was made and the layer was adhered onto the bottom dish (GWST-5040, Willco). After incubation with concanavalin A to promote cell adhesion (*t* = 30 min, 100 μL, 0.5 mg/mL, C5275 Sigma,[131]), cells were plated onto the bottom dish surface and the cavity was covered at the top with a 22×22 cm cover glass (Ted Pella).

The following are the mRNAs were synthesized *in vitro* using the mMessage mMachine Kit SP6 Kit(Ambion AM1340M). They were injected at the one-cell stage together with the microspheres :

- Lap2B-GFP[12] (inner nuclear membrane visualization) 80 pg per embryo
- GPI-GFP (plasma membrane visualization, we thank Michael Brand for the pCS2 + GPI-GFP plasmid) 50 pg per embryo
- RFP-Lamin A (Lamin A overexpression, this study) 200 pg per embryo
- NLS-GFP (soluble nuclear marker), 100 pg per embryo

Pharmacological inhibitors were used at the following concentrations: 100nM latrunculin A [42], 10μM blebbistatin(−) (Tocris Bioscience), 1 and 10μM nocodazole (Sigma), 5μg/mL brefeldin A (Sigma-Aldrich) [132].

Alexa Fluor 488 Annexin V staining (ThermoFisher) was used for the detection of apoptotic cells. DNA-Hoechst (ThermoFisher) was used to stain the cell nucleus and was incubated for 7min at 1μg/mL, ER-Tracker Green (BODIPY FL Glibenclamide) was used to stain the endoplasmatic reticulum (Extended Data Fig. 8c, d) at 1μM with an incubation time of 20min, as reported in the corresponding protocols. After incubations, cells were washed by centrifuging them at 200g for 3min and resuspended in DMEM [12, 51].

##### Active microrheology measurements in zebrafish progenitor stem cells

Details of the trapping experiments and microsphere selection are described in reference [51]. To per-form active microrheology measurements inside a cell, first, a cell with one or two microspheres in the cytoplasm is selected. Then, a working optical plane is identified looking at nucleus fluorescence, where the nucleus has its biggest cross-section. Subsequently, the microsphere is trapped and placed in the selected plane between the plasma and nuclear membrane, avoiding adhesions to either one. After placement, the active microrheology is initialized, with parameters shown in Supplementary Table 5.

After the measurement of the cytoplasm is completed, to get the measurement in the cytoplasmnucleus interface, the bead is placed to be in contact with the nuclear envelope. This is done by slowly moving the microsphere toward the nucleus and following the force signal. When the microsphere touches the NE there is a peak of force followed by an exponential relaxation after the bead stops moving. In this position, the microrheology routine is once again started, this time with the oscillations being perpendicular to the NE. Finally, after the bead is inserted in the nucleus with the protocol described in the following section, the same routine is performed with the microsphere trapped inside the nucleus. The oscillations of this measurement are done perpendicular to the insertion direction. All active microrheology measurements are performed using a trap power of 60mW to 100 mW at the sample plane and were found not to influence sample behavior (Supplementary Text 5.1).

##### Microsphere insertion into the nucleus

The process of inserting the microsphere inside the cell nucleus consists of applying a routine in which the force applied by the optical trap is clamped using the optical force feedback system explained above (Force set-point *F* = 100 − 200 pN). In this way, the force applied is constant and what is changing is the trap position and thus the microsphere position. First, the bead is placed in contact with the nucleus at the optical plane where its fluorescent cross section is the biggest. Then the clamp is set from 100 pN to 150pN and it is activated until the bead is inserted, or until the microsphere does not indent more or gets lost. The first 5s of the bead trajectory, and before insertion, were used to calculate the creep compliance of the nucleus.

##### Analysis of the creep-compliance data

Microsphere displacement while clamping the force was analyzed as a creep-compliance routine fitted with a Jeffrey’s model using a custom Python script. All experimental routines were analyzed, including those where insertion was unsuccessful. For successful insertion trials, only the initial phase—during which the bead indents but does not penetrate the nucleus—was considered. For these routines, the normalized position was fitted using[117]:

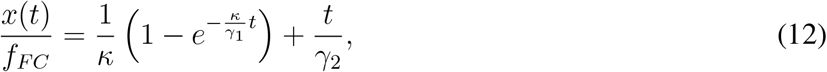

where *x*(*t*) is the displacement of the microsphere which is divided by the clamping force *f*_*F C*_. This normalization compensates for variations in the forces used for bead insertion in different cells. The fittings allow the extraction of the restoring stiffness *κ* and the viscoelastic drags *γ*_1_ and *γ*_2_.

#### Experiments in *C. elegans*

##### *C. elegans* maintenance

Strains were maintained and manipulated under standard conditions [133]. Nematode strains were grown at 20°C on nematode growth medium (NGM) plates with OP50 bacteria and, synchronized using the standard alkaline hypochlorite treatment method [134]. Only age-matched, synchronized adult hermaphrodites were used in this study. N2 wildtype strain was used as a control unless otherwise stated.

##### *C. elegans* transgenesis

Nematode strains used in this study are described in SuppTable 6. MSB1136 was generated in two steps. First, *sfGFP11* ssODN sequence (Table 2) was endogenously inserted at the N-terminus of lmn-1 locus by HDR (homology-directed repair) using CRISPR-Cas9 technology and following a co-CRISPR strategy using *dpy-10* as a marker to enrich for genome-editing events [135, 136, 137, 138]. Thus, obtaining MSB1126. Guide RNA (SuppTable 7) was designed using both Benchling (www.benchling.com) and CCTop [139] online tools. Second, MSB1126 was crossed to CF4588 [140] (TABLE 1) provided by CGC, finally obtaining MSB1136 (TABLE 1). MSB1136 shows high expression LMN-1::sfGFP signal in somatic cells and does not present any phenotype in terms of development and lifespan compared to the reference wildtype strain N2, thus considering MSB1136 as a wildtype-like strain.

**Table 2.** Reagents for PAA Gels of different stiffness. All quantities are in microliters

| estimated stiffness | Acryl-Bisacryl | Water | TMED | APS | Beads |
| --- | --- | --- | --- | --- | --- |
| 1-10 Pa | 41 | 540 | 1.5 | 5 | 5 |
| 10-100 Pa | 46 | 535 | 1.5 | 5 | 5 |
| 100-1000 Pa | 55-65 | 526-516 | 1.5 | 5 | 5 |
| 30000 Pa | 400 | 178 | 1.5 | 5 | 5 |

##### Lipid droplet isolation

*C. elegans* lipid droplets isolation was performed following the protocol previously described [81] with minor modifications. Briefly, for sample preparation, MSB1136 animals were grown on peptone-enriched plates seeded with NA2 bacteria and, synchronized using the standard alkaline hypochlorite treatment method [134]. 2 × 10^4^ synchronized larva 1-stage nematodes were plated and grew at 20°C until either day 1 or day 8 adulthood for sample collection. For lipid droplets day 8 collection, animals were washed off with M9 and separated from laid eggs by gravity, to avoid generation mixing, before transfer to new plates. On the day of the experiment, plates were washed off, and animals were collected using PBS. After washing the sample three times with buffer A [81], the pellet was resuspended in the same buffer supplemented with protease inhibitor (Sigma Aldrich, P8340). From now on, the sample was manipulated either on ice or at 4°C. Nematodes homogenization and cell disruption were performed by using an ultrasonic bath (VWR, USC300TH) four times, 1 min per time, with 30-s intervals. Lipid droplets were collected in buffer B [81] from the post-nuclear fraction by ultracentrifugation followed by three washing steps as described in Ding et al. 2013. Isolated lipid droplets were used within the same day for active microrheology routine in polyacrylamide gels (see section above).

##### Refractive index matching assay

Refractive index of freshly isolated lipid droplets were measured following the standard procedure [103] using the commercially available iodixanol solution (OptiPrep, Sigma Aldrich, D1556) [82]. Lipid droplets solution were gently mixed, accordingly, at different iodixanol concentrations until reaching the matching refractive concentration (48%) and, 0.5% membrane dye BioTracker NIR750 (Sigma Aldrich, SCT113). For lipid droplets’ optical trap scanning and imaging, the sample was mounted in optical trapping cavity and then sealed with a 1.5-cover glass, as previously described.

##### Lipid droplets characterization by TimSOM *in vitro*

To prepare PAA gels of different stiffness containing both lipid droplets and microspheres, freshly isolated lipid droplets solution was gently mixed with the reagents noted in Table 2. The water was substituted by the same volume of lipid droplet solution for each class of PAA gel. The mix was transferred into the optical trapping cavity and then sealed with a #1.5-cover glass, as previously described. To extract both lipid droplet size and stiffness values measurements were performed alternatively using lipid droplets and microspheres following the procedure detailed above.

##### *in vivo* TimSOM on *C. elegans* intestinal epithelial cells during ageing

All strains were seeded on NGM plates on the same day and let them grow until day 1 (three days post-seeding) and day 8 (ten days post-seeding) adulthood. For animals evaluated on day 8, they were transferred to new plates every day during the egg-laying period to avoid mixing generations. Because both LW697 and BN20 presented one-day developmental delay compared to wild type and BN19, active microrheology measurements at day 1 and day 8 adulthood for those strains were performed four days and eleven days post-seeding, respectively. The day of the experiment, nematodes were mounted on 2% agar pads, immobilized with 10 μm levamisole hydrochloride solution (Sigma Aldrich, 31742), and then covered with a 25 mm-by-25 mm cover glass (Ted Pella, #1.5) sealed with fingernail polish. Active microrheology was performed, as described above for zebrafish progenitor stem cells, on *C. elegans* intestinal cells using endogenous lipid droplets within 1h after immobilization to avoid damage to the animals.

##### Cuticle curvature evaluation

Cuticles of different age-matched animals were stained with slight modifications as previously described[141]. In short, *C. elegans* strains D1 and D8 were collected and washed using a solution of 0.5 % Triton X-100 and M9. Then, animals were incubated in 0.33 mg/mL DiI solution (Sigma Aldrich, 42364) for three hours, in agitation and in the dark. Three washes with M9 were made prior to animal mounting, under the same conditions as for TimSOM experiment. For curvature evaluation, Z-scan images were made from different parts of the worm body. From orthogonal sections, the width and height of each portion were measured and width/height ratios were calculated for each condition.

### 5.4 Protein expression and sample preparation

MEC-2 C-terminal and UNC-89 SH3 domain proteins, as well as CPEB4, were expressed in *E. coli* and purified as described[11, 64]. The MEC-2::UNC-89 sample was prepared in a 10:1 mixture (200:20 μM) at room temperature. The CPEB4 sample was prepared at a 25 μM final concentration with 300 μM Salt Buffer, at room temperature.

For dual trap active-passive microrheology, the surface of the lower coverglass was passivated using this protocol [142]. We used PEG beads (*d* = 1 μm, Micromod, 01-54-103), which were outside the protein condensate and were partially attached to it after short contact [11].

For TimSOM, the coverglass was treated with poly-L-lysine coupled to a polyethyleneglycol group (PLL-PEG, 30 min, 0.5 mg/ml, PLL(20)-g[3.5]-PEG(2), SuSoS). Droplets partially attached to the coverglass, such that the gradient force did not move the floating droplets during the rheology routine. We used carboxy-terminated microspheres that exclusively were found to enter inside the condensates (*d* = 1 μm, Thermofisher, F8816). Measurements of *G*(*ω*), from 0.125 Hz to 512 Hz, were repeated, for all the droplets tested, N=5 times in order to increase the signal-to-noise ratio. Laser power was kept below 50 mW to avoid heating of the protein condensate.

### 5.5 Data analysis

The analysis of the data measured with the active microrheology is performed in RheoAnalysis (IM-PETUX). In this program, the complex shear modulus’ is retrieved performing the required compensations (Supplementary Text 1.1.12). Furthermore, fittings from first two third-order models can be applied to the data set by selecting the respective initial conditions. Analysis of other calibration experiments such as the Strokes-drag force measurements, creep compliance measurements, and simulations was done on personalized Matlab and Python scripts.

## Notes

### Competing Interest Statement

PAF is holder of the patent #WO/2022/171898 protecting the time-shared rheology technique. XS is cofounder of Nuage Therapeutics. All other authors do not declare any conflicts of interests.

### Summary of Updates

Presentation of the results and methods.

