## Supplementary material for "Measuring age-dependent viscoelastic properties of organelles, cells and organisms via Time-Shared Optical Tweezer Microrheology": si material

3.2 Rheology of the zebrafish cytoplasm is unaffected by bead insertion into the nucleus 49

**4 Supplementary Text 4: Aberrations of the optical microscopy and optical trapping performance inside *C. elegans* 50**

**5 Supplementary Text 5: General consideration for intracellular rheology with optical tweezers 53**

**6 Supplementary Data Tables 57**

**7 Supplementary Videos 59**

### 1 Supplementary Text 1: Time-sharing - Concepts and Implementation

#### 1.1 Introduction

The use in biomechanical measurements of active microrheology performed with optical tweezers is increasing [1, 2, 3, 4, 5, 6]. The most widely used methods are based on the active-passive methodology introduced by [7] where data acquired from the passive power spectrum are combined with the active power spectrum to obtain the trap stiffness $k$. The latter quantity is essential to fix the $k/(6\pi a)$ scale of the complex G-modulus that can be deduced from the active power spectrum data $\hat{R}_{L-S}(\omega)$, where $a$ is the probe radius.

The active spectrum $\hat{R}_{L-S}(\omega)$ depends on the ratio of two positions: the position of the laser $\hat{x}_L(\omega)$ or stage $\hat{x}_S(\omega)$, which are known positions, and the position of the bead $\hat{x}(\omega)$, which change as response of the movement of the laser or stage and must therefore be measured. Measuring the position of the bead can be done in several ways: using a camera to track the particle [8, 9] or by using Back-Focal-Plane Interferometry (BFPI) [10] to get its displacement. However, most cameras have a rather low spatial resolution and are limited by their frame rate, but the position of the micro-sphere can be calibrated a priory and in principle multiple microspheres can be tracked at the same time. In contrast, the use of BFPI requires in general a *in situ* calibration of $\beta$ [11, 12], which is the conversion factor of the V signal to displacement $\Delta x$. This is not the case if the measurement is carried out by using the optimized BFPI, based on direct measurement of light momentum changes [13, 14], which led to the direct measure of the force exerted by the trap on the particle without the need of any *in situ* calibration.

Based on this passive-active methodology, the different groups developed their own methods [15, 16, 17] to reduce the time required to perform the necessary calibrations and data acquisition time, particularly because of the high time cost of low-frequency measurements, which moreover can call into question the validity of the measurement [18].

Of all the possible configurations to perform microrheology, the one that uses an additional detection laser[1, 3, 19, 20, 21] to measure bead displacement is the best performing in terms of its ability to measure probes with high G, being able to measure uncured PDMS samples with G between $10^4$ and $10^5$ Pascal [12]. However, the use of an additional laser increases the complexity of the equipment and its operation due to alignment and constraints in the experimental field of view.

Here, we have developed a new method for micro-rheology measurements in rigid samples with a single, time shared laser trap to drive the trapped microspheres and measure their displacements (Main Text Fig. 1a).

#### 1.2 The sensitivity calibration $\beta$ erased by symmetry

The method uses two traps generated by the same laser source that act simultaneously on the bead. Before starting the active micro-rheology routine, trap 1 and trap 2 are both placed in the center of the particle. Once centered, trap 1 starts oscillating at a driving frequency $\omega$, i.e $x_1(t) = x_0 \sin(\omega t)$, while trap 2 remains fixed at $x_2(t)$=0, with $x_0$ as the amplitude of the oscillation and $x_1(t)$ and $x_2(t)$ as the particle trajectory of the driving and the static trap on the lab frame. In other words: Trap 1 will act as a *driving trap*, while trap 2 will behave as a static, *detection trap.* The method uses BFPI to obtain two voltages $V_1(t)$ and $V_2(t)$ representative of the forces or, equivalently, of the displacement of the traps from the center of the bead, since for small displacements the following linear relationship holds $F_1(t) = k_1(x_1(t) - x(t)) = k_1\beta_1 V_1(t)$ and $F_2(t) = k_2(x_2(t) - x(t)) = k_2\beta_2 V_2(t)$, where $x(t)$ is the trajectory of the centre of the bead and at the start of the micro-rheology routine $x(t = 0) = 0$. Without loss of generality it is assumed that the two traps behave identically, i.e., $\beta_1 = \beta_2 = \beta$ and $k_1 = k_2 = k/2$, where $k$ is the sum of the stiffness of the two traps. The active power spectrum [7] defined by

$$\hat{R}_L(\omega) \;=\; -\frac{k}{i\omega}\hat{\chi}_a(\omega) \tag{1}$$

and introduced using the response function of the active-passive system $\hat{\chi}_a(\omega)$

$$\hat{x}(\omega) \;=\; \hat{\chi}_a(\omega)\hat{F}_a(\omega), \tag{2}$$

which describes the reaction of the position of the bead $\hat{x}(\omega)$ in the transition from the passive stage $\hat{F}_a(\omega) = 0$ to the active stage $\hat{F}_a(\omega) \neq 0$, can be written using the Fourier transform $\hat{V}_1(\omega)$ and $\hat{V}_2(\omega)$ of the acquired signals $V_1(t)$ and $V_2(t)$.

By definition, $\hat{F}_a(\omega)$ carry only the force contributions coming from the movement of the traps. Because only $x_1(t)$ is moving it follows that $\hat{F}_a(\omega) = k\hat{x}_1(\omega)/2 = \hat{F}_1(\omega) - \hat{F}_2(\omega) = k\beta(\hat{V}_1(\omega) - \hat{V}_2(\omega))/2$. On the other side $\hat{x}(\omega)$ is directly connected to the signal of traps 2, in fact $\hat{x}(\omega) = -2\hat{F}_2(\omega)/k = -\beta\hat{V}_2(\omega)$. Thus the response function of the active-passive system reads

$$\hat{\chi}_a(\omega) = \frac{\hat{x}(\omega)}{\hat{F}_a(\omega)} = -\frac{2\hat{V}_2(\omega)}{k\left[\hat{V}_1(\omega) - \hat{V}_2(\omega)\right]}, \tag{3}$$

and, the active power spectrum follows straightforward

$$\hat{R}_L(\omega) = \frac{2\hat{V}_2(\omega)}{i\omega\left[\hat{V}_1(\omega) - \hat{V}_2(\omega)\right]}. \tag{4}$$

Above is a new notation for the response function $\hat{\chi}_a(\omega)$ of the active-passive system, new to that used by Fischer [7], to distinguish it from the response function $\hat{\chi}(\omega)$ of the bead embedded in the viscoelastic material, which for its part reads

$$\hat{x}(\omega) = \hat{\chi}(\omega)\hat{F}_{tot}(\omega), \tag{5}$$

where now $\hat{F}_{tot}(\omega)$ is the total force exerted by the traps and acting on the bead. Whereas for $\hat{F}_a(\omega)$ the contribution came only from the motion of trap 1, for $\hat{F}_{tot}(\omega)$ the static contributions are also taken into account, which arise from the fact that even when the traps are stationary but there is a difference in position between the traps and the center of the bead an external force from the laser acts on the bead. $\hat{V}_1(\omega)$ and $\hat{V}_2(\omega)$ are proportional to this position difference. Explicitly $\hat{F}_{tot}(\omega) = \hat{F}_1(\omega) + \hat{F}_2(\omega) = k\beta(\hat{V}_1(\omega) + \hat{V}_2(\omega))/2$, and

$$\hat{\chi}(\omega) = \frac{\hat{x}(\omega)}{\hat{F}_{tot}(\omega)} = -\frac{2\hat{V}_2(\omega)}{k\left[\hat{V}_1(\omega) + \hat{V}_2(\omega)\right]}, \tag{6}$$

In all response functions mentioned up to now, forces and particle displacements are supposed to be co-linear with a vector passing through the center of the bead. The susceptibility of the active-passive system differs from that of the bead by having in parallel the stiffness of the trap

$k$, i.e.

$$\frac{1}{\hat{\chi}_a(\omega)} = k + \frac{1}{\hat{\chi}(\omega)}. \tag{7}$$

Its connection with the G-modulus is given by

$$\hat{G}(\omega) = \frac{1}{6\pi a}\left[\frac{1}{\hat{\chi}(\omega)} + \omega^2 m\right], \tag{8}$$

with $a$ as the radius of the probe. But since the inertial term $\omega^2 m$ appearing in other works is irrelevant at the frequencies of our interest, $\omega^2 m$ becomes significant for frequencies higher than 10 MHz, the following expression holds from here on

$$\hat{G}(\omega)\hat{\chi}(\omega) = \frac{1}{6\pi a} \tag{9}$$

It follows the expression of the G-modulus for the proposed method

$$\hat{G}(\omega) = \frac{1}{6\pi a}\frac{1}{\hat{\chi}(\omega)} = -\frac{k}{12\pi a}\frac{\hat{V}_1(\omega) + \hat{V}_2(\omega)}{\hat{V}_2(\omega)}. \tag{10}$$

Surprisingly the method does not require any explicit knowledge of the conversion factor $\beta$ to get the active power spectrum $\hat{R}_L(\omega)$ or to determine $\hat{G}(\omega)$. This is because the signals $V_1(t)$ and $V_2(x)$ have the same conversion factor.

That means the accuracy of the proposed method is based on the ability to generate two identical traps from a single laser source and to obtain two measurements of $V_1(t)$ and $V_2(t)$ that have identical sensitivities.

##### 1.3 The stiffness of the traps $k$ fixes the scale and the precision of the measurement

The precision of the method from its part depends strongly on the selection of the measurement scale

$$G_0 = \frac{k}{12\pi a} \tag{11}$$

which is adjustable by varying the power of the laser beam which control the strength of the total traps stiffness $k$. If $|\hat{G}(\omega)|$ is much larger than $G_0$ then $|\hat{V}_1(\omega) + \hat{V}_2(\omega)| >> |\hat{V}_2(\omega)|$,

$$\hat{G}(\omega) \approx -\frac{k}{12\pi a}\frac{\hat{V}_1(\omega)}{\hat{V}_2(\omega)}, \tag{12}$$

and the relative error over the magnitude of the G-modulus is of the order of the relative error of the signal measurement.

$$\frac{\Delta|\hat{G}(\omega)|}{|\hat{G}(\omega)|} \approx 2\frac{\Delta|\hat{V}(\omega)|}{|\hat{V}(\omega)|}. \tag{13}$$

This corresponds to the case where the material is much stiffer (or viscous, i.e $(6\pi a)\mu\omega > k$) than the trap. In that case the force exerted by the driving trap, trap 1, is much higher than the force exerted by the static trap, trap 2. This is the right regime we were working in to obtain precise measurements.

The regime to avoid, takes place when the scale $G_0$ is much larger than $|\hat{G}(\omega)|$ in this case $|\hat{V}_1(\omega) + \hat{V}_2(\omega)| << |\hat{V}_2(\omega)|$ and the precision on the magnitude of the G-modulus can becomes very poor because

$$\frac{\Delta|\hat{G}(\omega)|}{|\hat{G}(\omega)|} \approx \frac{|\hat{V}_2(\omega)|}{|\hat{V}_1(\omega) + \hat{V}_2(\omega)|}\frac{\Delta|\hat{V}(\omega)|}{|\hat{V}(\omega)|}. \tag{14}$$

In that case the forces exerted by the two traps, trap 1 and trap 2, are practically identical in magnitude but they have opposite orientation. The relative error over the magnitude of the G-modulus becomes much larger than those of the single signals.

#### 1.4 The time-sharing implementation of the method and the criterion of simultaneous traps

To put the methods into practice by generating two identical traps two strategies are easily identifiable. The first is to separate the laser beam into two beams with orthogonal polarization. The second is to create the two traps by time-sharing, that is, by high-frequency multiplexing the laser position between the two trap positions. Both strategies have their merits and drawbacks.

Separation by polarization satisfies the criterion that the two traps must act simultaneously on the bead for the Eqs. (6) and (10) to be valid. It does, however, suffer from the drawback of having to align two different optical paths and having to introduce two pairs of deflectors as in the case of using a separate detection laser. It is also necessary to work with two BFPI systems, one for each polarisation, to get the two signal $V_1(t)$ and $V_2(t)$.

The solution based on time-sharing violates the criterion of simultaneous traps but is much more simple from an optical point of view. The optical path to be aligned is only one, and with a single detector it is possible possible to extracts $V_1(t)$ and $V_2(t)$ by synchronising the detector sampling with the steering of the two traps. The two traps can be moved using a single pair of deflectors.

In this paper, the time-sharing strategy[22] is used to implement the method described so far. A pair of acoustic optical deflectors (AODs) are used to deflect at high frequency, the laser beam between the positions of the two traps. Since the solution does not satisfy the criterion of simultaneity of the traps, but they appear intermittently and complementary to each other, an important part of this manuscript is devoted to the resulting consequences. In particular, predictions of the deviation of the raw measurements obtained by time-sharing from the expected result in the case of using two simultaneous traps are presented.

It will be shown that the deviation for most practical cases involving biological samples, such as the cytoplasm of most common cells, is negligible. For viscoelastic liquids, e.g. Maxwell materials, the deviations strongly affect the real part of the response function, such that the solid behaviour at frequencies larger than the crossover frequency is not accessed for inadequately stiff traps. In this particular case the time-sharing measurement degenerates and the elastic component of the material becomes inaccessible to the measurement. Our simulations also show that the real part $\chi_1'(\omega)$ but not the imaginary part of the response function is affected (the real and imaginary parts are convolved and cannot be distinguished), and that such behavior is proportional to the rigidity of the trap $k$. However, as we'll discuss further, the first-order term in the expansion (Eq. 36) actually contains information about the elastic part of the Maxwell model, which helps to resolve this issue. Thus, measuring the response function with different trap stiffness is able to break the degeneracy of the time-sharing technique and allows access to $E$ in the Maxwell regime. For all other types of samples (including the fractional Maxwell systems) a measurement compensation method has been developed to correct the deviation induced by the intermittent nature of the traps.

#### 1.5 Two approaches to study the deviation of the method due to its implementation by time-sharing technique

The prediction of the deviation is done by studying the measured G-modulus $G_m(\omega)$ for viscoelastic materials behaving according to the fractional Kelvin-Voigt and fractional Maxwell models. These two models are seen as generalized viscoelastic models that extend the applicability of the classical Kelvin-Voigt, Maxwell, and structural damping model [23] to describe with few parameters the power-law behavior of most biological samples and gels [24]. The next subsection introduces their mathematical description.

Based on the fractional models, two complementary approaches were used to study the deviation. The first is based on numerical resolution of Fractional Differential Equations (FDEs) to obtain the dynamics of the bead under the effect of the two intermittent traps. Since fractional derivatives are non-local linear operators, the simulation takes more time $O(n^2)$ than solving ordinary differential equations $O(n)$, where $n$ is the number of simulated time points. The second approach is analytical and is based on an approximation we have called: First Harmonic Approximation (FHA). The response of the bead is predicted in the frequency domain. The solution is an analytical, explicit equation expressing the relationship between the measured response function $\hat{\chi}_m(\omega)$ and that expected $\hat{\chi}(\omega)$ if two simultaneous traps are used. The analytical solution is immediate, simple and easy to interpret. A data compensation procedure was developed from the FHA approximation. The FDE approach, on the other hand, is more accurate and is an excellent tool for checking the accuracy and validity of FHA predictions.

Our practical implementation of the time-sharing technique passed through the use of acousto-optical AOD deflectors to deflect the laser beam between the positions of the two traps with a steering frequency of $25kHz$. The deflection is due to Bragg diffraction of the laser beam on refractive index fringes created by the passage of an acoustic wave through a birefringent crystal.

The deflection is holographic in nature. The laser beam gradually disappears from the position of trap 1 to appear progressively at the position of trap 2, without having to pass through the points connecting the two positions, contrary to what is obtained by using steering mirrors. The sum of the intensities of the two traps always remains constant even during the transition from one position to the other. Assuming that the transition between the two traps is linear in time and that their positions lie in the linear region of the bead where the stiffness k is constant, then the two forces are given by $F_1 = k(1 - t/\tau)(x_1 - x(t))$, $F_2 = -k\, x(t)\, t/\tau$ (where $t \in [0, \tau]$, $[0, \tau]$ is the transition interval which in our case measures $10us$) and since both act on the same rigid bead, the resultant force is given by the total force $F_{tot}(t) = F_1(t) + F_2(t) = k(1 - t/\tau)x_1 = kx_l(t)$ with $x_l(t) = (1 - t/\tau)x_1$. This means that the two time-sharing traps can be described as a single trap continuous trajectory $x_l(t)$ connecting positions of trap 1 and trap 2. Finally, the effect on the bead is identical to that obtained in the case of using steering mirrors, although the way of acting on the bead is different.

Supplementary Fig.1 a) shows the trajectory of the $x_l(t)$ trap used in the case of the FDE approach. The trajectory is characterized by a high-frequency trapezoidal waveform with a plateau of $30\mu s$ and rise and fall times of $10\mu s$; at each time-sharing cycle the amplitude of the trapezoid is modulated with the position of trap 1 $x_1(t_{1,i})$. The red and blue dots show the time points, $t_{1,i}$ and $t_{2,i}$, at the sampling of trap 1 and trap 2, respectively. Sampling is performed with a known delay of $33\mu s$ from the beginning of the trapezoid rising edge. The sampling of the two traps have a period of $80\mu s$ and are shifted by $40\mu s$. This delay between the traps sampling will have consequences.

Supplementary Fig.1 b) shows the approximate trajectory used for $x_l(t)$ in the first harmonic approximation (FHA), where

$$x_l(t) = x_1(t)\frac{[1 + \sin(\omega_t \cdot t)]}{2}. \tag{15}$$

The name given to this approximation comes from the fact that now $x_l(t)$ is approximated up to the first harmonic in the carrier $\omega_t = 2\pi f_t$ frequency. Where $f_t$ it corresponds to the time-sharing refreshing frequency that is half of the steering frequency $f_t = f_s = 25kHz/2 = 12.5kHz$.

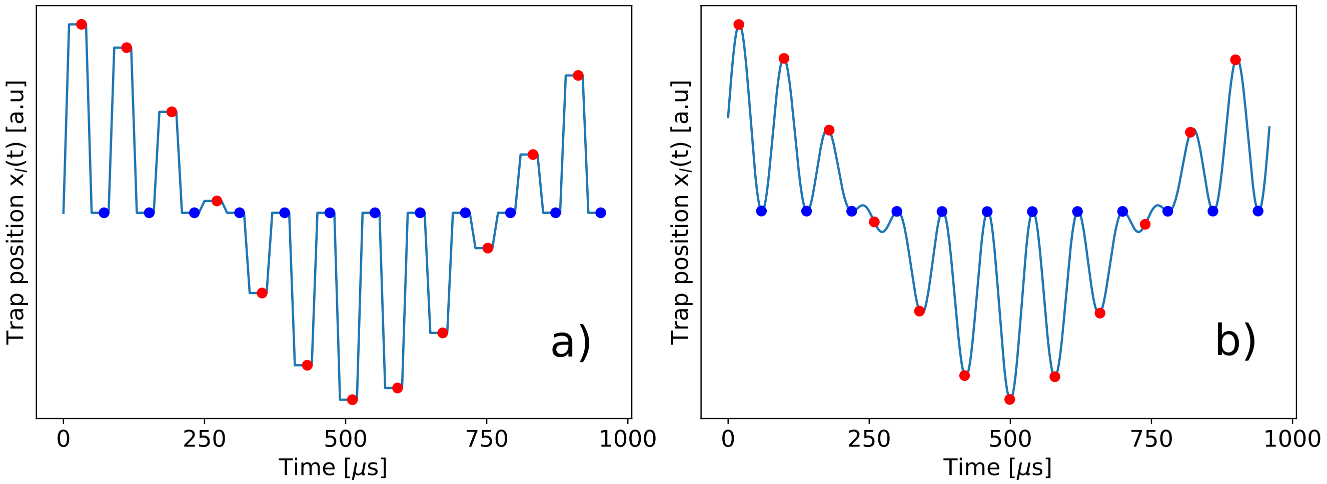

**Supplementary Fig. 1**: Trajectory of traps $x_l(t)$ generated by the time-sharing technique to emulate the two traps needed for the micro-rheological method under consideration. Red and blue dots are acquired at the time points, $t_{1,i}$ and $t_{2,i}$, at the sampling of trap 1 and trap 2, respectively. At these instants the position of $x_l(t_{1,i}) = x_1(t_{1,i})$ and $x_l(t_{2,i}) = x_2(t) = 0$. Left figure shows the trajectory used in the FDE simulation. The trajectory is characterized by a trapezoidal waveform with $12.5kHz$ frequency, a plateau of $30\mu s$ and rise and fall times of $10\mu s$; at each time-sharing cycle, the amplitude of the trapezoid is modulated with the position of trap 1 $x_1(t_{1,i})$. Right figure shows the trajectory used to obtain the FHA analytical solution. The trajectory is approximated up to the first harmonic in the carrier $\omega_t = 2\pi f_t$ frequency, see Eq. 15.

#### 1.6 The mathematical description of the fractional Kelvin-Voigt and fractional Maxwell models

The mathematical description of materials exhibiting viscoelastic power-law behavior goes through the use of the fractional spring-pot element, see Supplementary Fig. 2 , as the constitutive unit for building the viscoelastic model. This fractional element depends on two parameters: $c$ and $\alpha$ [24].

From its representation in the space of frequencies, see Eq. 16, it is easy to see that $c$ defines the magnitude of the G modulus in [Pa] at $\omega = \omega_0 = 1[rad/s]$, and, that $\alpha$ is its power-law exponent and can span $\alpha \in [0, 1]$. When $\alpha = 0$ the spring-pot element behaves like a spring of stiffness $c = E\,[Pa]$, while when $\alpha = 1$ its behavior is that of a liquid of viscosity $\mu = c/\omega_0[Pa.s]$. Interestingly, the exponent $\alpha$ allows for a continuous description of the transition between a Newtonian liquid $\alpha = 1$ and a Hookean solid $\alpha = 0$. The constant $\omega_0$ is necessary to define the meaning of $c$ and to give consistency to the physical units of the model, but from here on it will be omitted from the equations and will be implicitly included in $\omega/\omega_0 \to \omega$.

The description of the spring-pot element in the time domain, Eq. 17, is more complex than its frequency representation. The stress $\sigma(t)$ of the material depends on the fractional derivative $\alpha$ of its strain $\epsilon(t)$. The fractional derivative $D_t^\alpha$ is a nonlocal linear integral operator used to generalize the concept of the derivative of order $n$, $d^n/dt^n$, restricted to $n$ belonging to the natural number, to a positive arbitrary real number $\alpha$. There are many different definitions for the fractional derivative. Eq. 17 uses Caputo's definition [25]. However, in the case of linear rheological problems one can also use the Riemann-Liouville definition, which yields equivalent models [26]

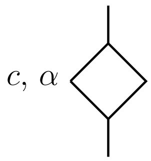

$$\hat{G}(\omega) = \frac{\hat{\sigma}(\omega)}{\hat{\epsilon}(\omega)} = c\left(i\frac{\omega}{\omega_0}\right)^\alpha \qquad (16)$$

$$\sigma(t) = cD_t^\alpha \epsilon(t) = \frac{c}{\Gamma(1-\alpha)}\int_0^t (t-t')^{-\alpha}\frac{d\epsilon(t')}{dt'}dt' \qquad (17)$$

**Supplementary Fig. 2**: The spring-pot element. Its symbol and equations, used to generalize the behavior of viscoelastic materials, ranging from the Hookean solid $\alpha = 0$ to the Newtonian liquid $\alpha = 1$, via the gel point $\alpha = 0.5$ [27]. Eq. 16 shows its representation in the frequency domain $\omega$. $\hat{G}(\omega)$ is given by a simple algebraic equation where $c$ is the magnitude of the modulus G at frequency $\omega = \omega_0 = 1$. Eq. 17 shows its description in the time domain. Here the springpot element connect the stress $\sigma(t)$ on the material to the fractional derivative $\alpha$ of its strain $\epsilon(t)$. The fractional derivative $D_t^\alpha$ follows here the Caputo's definition [25].

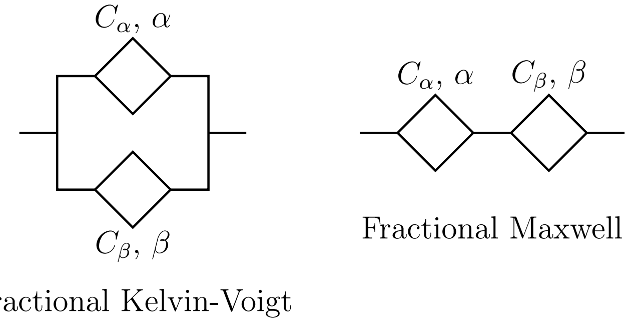

**Supplementary Fig. 3**: The two generalized viscoelastic models used to describe the samples. If $\alpha = 0$ and $\beta = 1$, or vice versa, the two fractional models correspond to the classical Kelvin-Voigt viscoelastic model and the Maxwell viscoelastic model, respectively. In the case where $\alpha \neq 0$ and $\beta = 1$ Kelvin-Voigt's fractional model corresponds to the structural damping model [23]

The fractional Kelvin-Voigt and fractional Maxwell models are obtained by connecting two springpot elements in parallel and series, respectively, as shown in Supplementary Fig 3. The G modulus and response function of the material described by these models read,

$$\hat{G}(\omega) = C_\alpha(i\omega)^\alpha + C_\beta(i\omega)^\beta \ , \ \hat{\chi}(\omega) = \frac{1}{6\pi a}\frac{1}{C_\alpha(i\omega)^\alpha + C_\beta(i\omega)^\beta}, \tag{18}$$

in case of fractional Kelvin-Voigt model, and,

$$\hat{G}(\omega) = \frac{1}{\frac{1}{C_\alpha(i\omega)^\alpha} + \frac{1}{C_\beta(i\omega)^\beta}} \ , \ \hat{\chi}(\omega) = \frac{1}{6\pi a}\left[\frac{1}{C_\alpha(i\omega)^\alpha} + \frac{1}{C_\beta(i\omega)^\beta}\right], \tag{19}$$

in case of fractional Maxwell model.

Supplementary Fig. 4 shows the real components $G'(\omega)$ and imaginary components $G''(\omega)$ of the G-modulus $\hat{G}(\omega) = G'(\omega) + iG''(\omega)$ for the two models given the same set of parameters: $C_\alpha$=2. 91, $\alpha$=0.143, $C_\beta$=0.127, $\beta = 0.7146$. The first springpot, $\alpha = 0.143 < 0.5$ is a viscoelastic solid, while the second springpot, $\beta = 0.7146 > 0.5$ is a viscoelastic liquid [27]. The difference between fractional Kelvin-Voigt fractional and fractional Maxwell is manifested in the asymptotic behaviors of the two models. For the low-frequency limit $\omega \to 0$, the Kelvin-Voigt behaves as the viscoelastic solid springpot, while for the high-frequency limit $\omega \to \infty$ it is the viscoelastic liquid that dominates. In the case of the fractional Maxwell it is exactly the opposite. In the case $\alpha < 0.5 < \beta$ for both models the real $G'(\omega)$ and the imaginary $G''(\omega)$ cross at the crossover frequency $\omega_c$ [24]. The crossover frequency $\omega_c$ separates the frequency ranges in which the models behave as viscous solids or as viscoelastic liquids. In the other cases, i.e. $\alpha < \beta < 0.5$ or $\alpha > \beta > 0.5$ there will be no crossover frequency.

#### 1.7 Study of the deviation via Fractional Differential Equations (FDEs)

The first approach to study the deviation is based on numerical solving of Fractional Differential Equations to get the displacement $x(t)$ of the bead, embedded in a viscoelastic material described by one of the two previously introduced fractional models, under the influence of a trap that follows the trajectory $x_l(t)$ shown in Supplementary Fig. 1a.

The FDE describing the movement equation of the bead for the the case of fractional Kelvin-

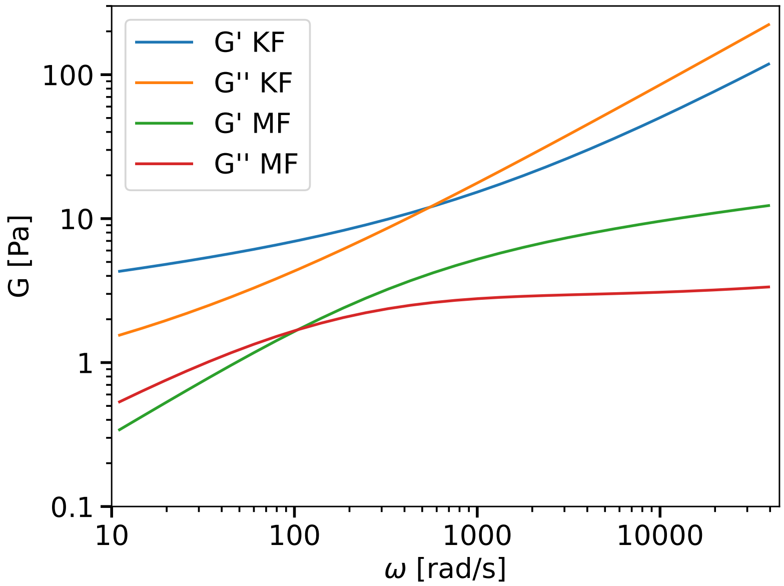

**Supplementary Fig. 4**: . The real $G'(\omega)$ and the imaginary $G''(\omega)$ components of the G-modulus $\hat{G}(\omega) = G'(\omega) + iG''(\omega)$ for the two fractional Kelvin-Voigt (FKV) and the fractional Maxwell (FM) models. The set of parameters is the same for the two models: $C_\alpha$=2.91, $\alpha$=0.143, $C_\beta$=0.127, $\beta = 0.7146$.

Voigt material reads

$$(6\pi r) \cdot \left[C_\alpha \, D_t^\alpha + C_\beta \, D_t^\beta\right] x(t) \quad = \quad k \left[x_l(t) - x(t)\right], \tag{20}$$

while in case of Maxwell fractional materials the movement is described by a set of FDEs which read

$$(6\pi r) \cdot C_\alpha D_t^\alpha x_1(t) \quad = \quad k \left(x_l(t) - x(t)\right), \tag{21}$$

$$(6\pi r) \cdot C_\beta D_t^\beta x_2(t) \quad = \quad k \left(x_l(t) - x(t)\right), \tag{22}$$

$$x(t) \quad = \quad x_1(t) + x_2(t) \tag{23}$$

The inertial term $mx''(t)$ has been omitted from the equations, because, as already mentioned, its contribution is negligible at the studied frequencies. Also the random force exerced on the bead by the surrounding media, which is considered in active-passive studies, is not considered in our simulations due to the fact the deviation affect only the active part of the microrheology.

To solve numerically the FDEs the discretized Gruenwald-Letnikov fractional derivative [28] has been used to estimate numericaly the Riemann-Liouville one. This last is equivalent to the Caputo definition in case of $x(0) = 0$ [26]. The "Short-Memory" principle [28] has been applied to bound the numerical effort to $O(n)$ iterations, where $n$ is the number of time point discretization.

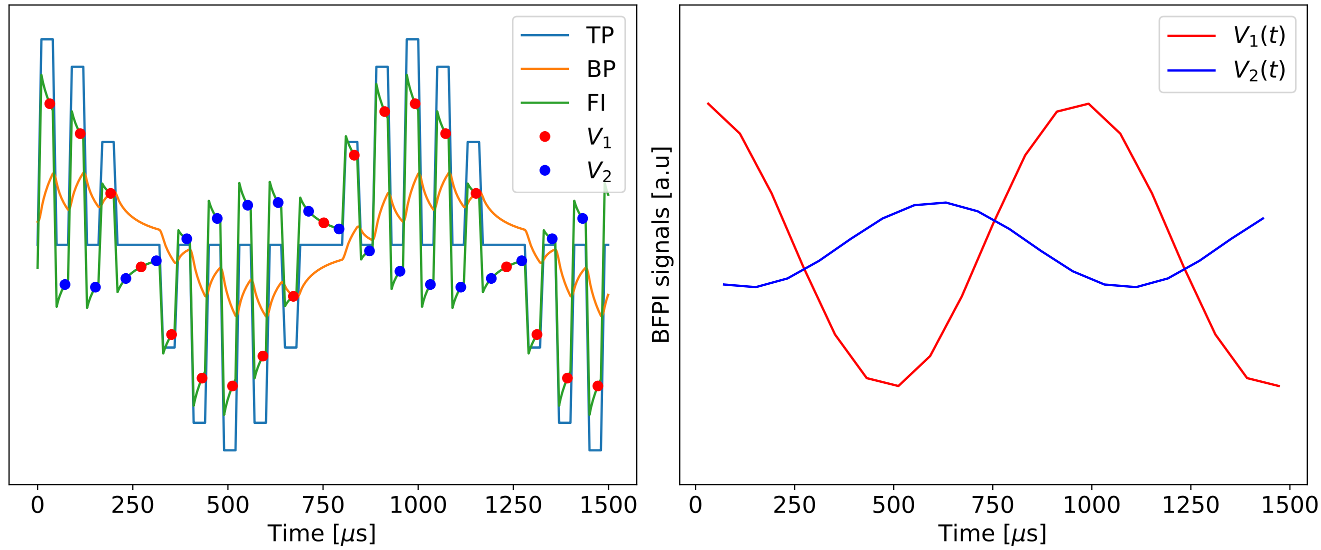

**Supplementary Fig. 5**: . The left Figure show the different data generated using the FDE approach for numerical calculation of $G_t(\omega_j)$ where: $\omega_j = 2\pi f_j$ and $f_j$=1000Hz; TP is the displacement of the trap $x_l(t)$; BP is the trajectory of the bead $x(t)$ obtained by solving the FDEs under the influence of the trap displacement; FI is the instantaneous force exerted by the trap on the bead $F(t) = k(x_l(t) - x(t))$; $V_1(t_{1,i})$ and $V_2(t_{2,i})$ are the signals measured by BFPI, proportional to $F(t_{1,i})$ and $F(t_{2,i})$, respectively, which allow us to determine $G_t(\omega_j)$ using Eqs. 24 and 25. The right figure shows the two trap signals $V_1(t)$ and $V_2(t)$ as shown to the user.

To obtain the value of the G-modulus measured by time-sharing $G_t(\omega_j)$ , for each frequency point $\omega_j$ it is necessary to generate the different data shown in Supplementary Fig. 5 in the following order.

First, the trap displacement $x_l(t)$ is generated for the given $\omega_j = 2\pi f_j$ (in the figure $f_j$=1000Hz). Second, the FDE is solved to find the displacement of the bead $x(t)$. Third, the instantaneous force exerted by the trap on the bead $F(t) = k(x_l(t) - x(t))$ is calculated. Fourth, the signals measured by the BFPI, $V_1(t_{1,i})$ and $V_2(t_{2,i})$, which are proportional to $F(t)$, are extracted by sampling the signal at well-defined time points $t_{1,i}$, $t_{2,i}$. Fifth, the fast Fourier transform (FFT) of the trap signals $V_1(i)$ and $V_2(i)$ is calculated and one of the two phases is corrected to account for the time delay $\tau_d = t_{2,i} - t_{1,i} = 40\mu s$ existing between the sampling of the two signals.

$$
\begin{aligned}
\hat{V}_1(\omega_j) &= FFT\left(V_1(i)\right) \\
\hat{V}_2(\omega_j) &= e^{-i\omega_j\tau_d} FFT\left(V_2(i)\right)
\end{aligned} \tag{24}
$$

Finally, the linear time-sharing response function $\hat{\chi}_t(\omega)$ and the G modulus $\hat{G}_t(\omega)$ can be derived using the formulas

$$
\hat{G}_t(\omega_j) = -\frac{k}{12\pi a}\frac{\hat{V}_1(\omega_j)+\hat{V}_2(\omega_j)}{\hat{V}_2(\omega_j)} \text{ and } \hat{\chi}_t(\omega_j)\cdot\hat{G}_t(\omega_j) = 6\pi a \tag{25}
$$

This simulation workflow was implemented to predict the deviation in case of fractional Kelvin-Voigt and fractional Maxwell samples.

Supplementary Fig. 6 shows the results for the fractional Kelvin-Voigt model with the springpot parameters $C_\alpha$=2.91, $\alpha$=0.143, $C_\beta$=0.127, $\beta = 0.7146$ (same as Supplementary Fig. 4). The simulation was run for two different values of the trap stiffness k (KF1: $k = 45[Pa]$, KF2: at $k = 450[Pa]$). The deviation is small for frequencies up to $1000[Hz] \approx 6000[rad/s]$. It remains lower than 15% for $k = 45[Pa]$ and lower than 30% for $k = 450[Pa]$. By increasing the trap stiffness, the magnitude of the deviation increases. In any case, for fractional Kelvin-Voigt samples, the time-sharing measurement correctly reflects the behavior of the sample.

In case of fractional Maxwell materials the situation is more complex. Supplementary Fig. 7 shows the results for this model with springpot parameters equal to those used for the fractional Kelvin-Voigt. Again, the simulation was run for two different values of the trap stiffness k (KF1: $k = 9[Pa]$, KF2: at $k = 90[Pa]$) In the Maxwell fractional case, the slope of $\hat{G}_m$ does not decrease once the cross over frequency $\omega_c$ is passed. The fractional solid behavior that should emerge at $\omega > \omega_c$ does not show up. Looking at $\chi_m$ it is clear that the deviation affects the real part, $\chi'_m$, much more than the imaginary part, $\chi''_m$. For measurements made with a sufficiently low value of $k$, in our example $k = 9[pN/um]$, the predicted deviation of $\chi''_m$ turns out to be less than 10%.

To understand the causes of the predicted deviation, we will examine the problem using the first harmonic approximation (FHA) in the next section. As we will see, it will still be possible to

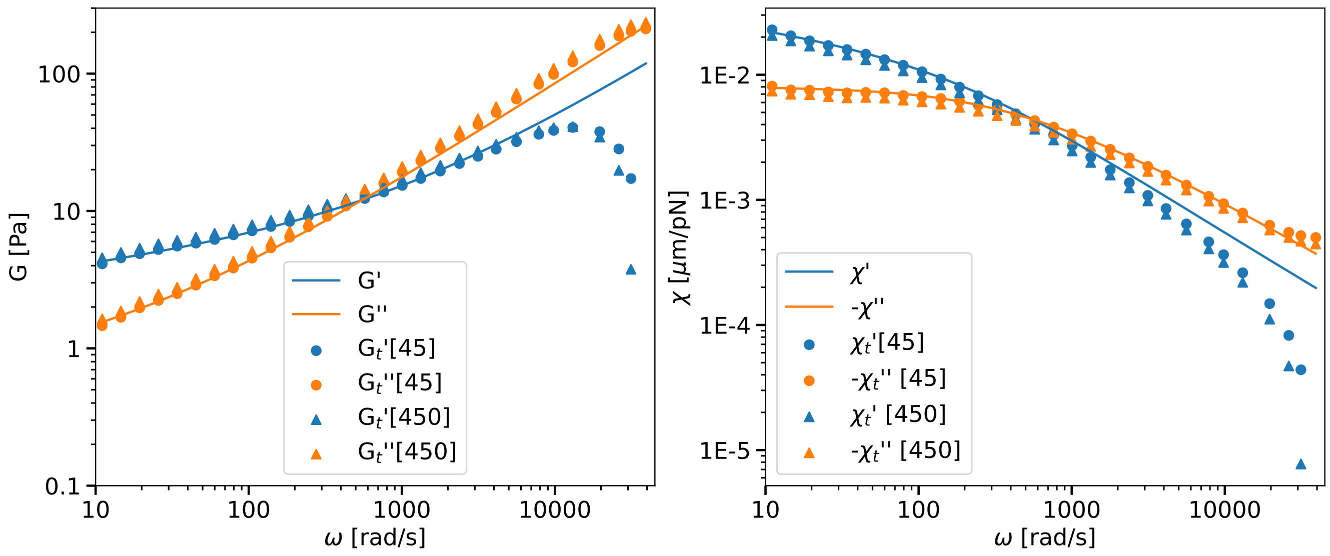

**Supplementary Fig. 6**: FDE prediction of the deviation of the rheological measure in the case of a fractional Kelvin-Voigt material. The set of model parameters is the same as in Supplementary Fig. 4. The measurement obtained by time-sharing: G-modulus $\hat{G}_t(\omega) = G'_t(\omega)+iG''_t(\omega)$ and linear response function $\hat{\chi}_t(\omega) = \chi'_t(\omega)+i\chi''_t(\omega)$. Model behavior: The G-modulus $\hat{G}(\omega) = G'(\omega)+iG''(\omega)$ and the linear response function $\hat{\chi}(\omega) = \chi'(\omega) + i\chi''_t(\omega)$ are used as references. Time-sharing predictions were computed for two different values of the trap stiffness k. [45] $\rightarrow$ k=45[Pa], [450] $\rightarrow$ k=450[Pa]. For KF1 the deviation is less than 15% up to frequencies of $1000[Hz] \approx 6000[rad/s]$.

compensate for the deviation and recover the true sample behavior for most fractional Maxwell samples, except for samples that exhibit classical Maxwell behavior, that is, where $\alpha = 0$ and $\beta = 1$.

#### 1.8 The First Harmonic Aproximation (FHA)

The second approach to the study of deviation is based on analytically solving the equations of motion of the bead $x(\omega')$ in the space of frequencies $\omega'$. The equation to be solved is the algebraic equation

$$\hat{x}(\omega') = \hat{\chi}(\omega')\, k\, [\hat{x}_l(\omega') - \hat{x}(\omega')]\,, \tag{26}$$

which becomes

$$\hat{x}(\omega') = \hat{\chi}_a(\omega')\, k\, \hat{x}_l(\omega'), \tag{27}$$

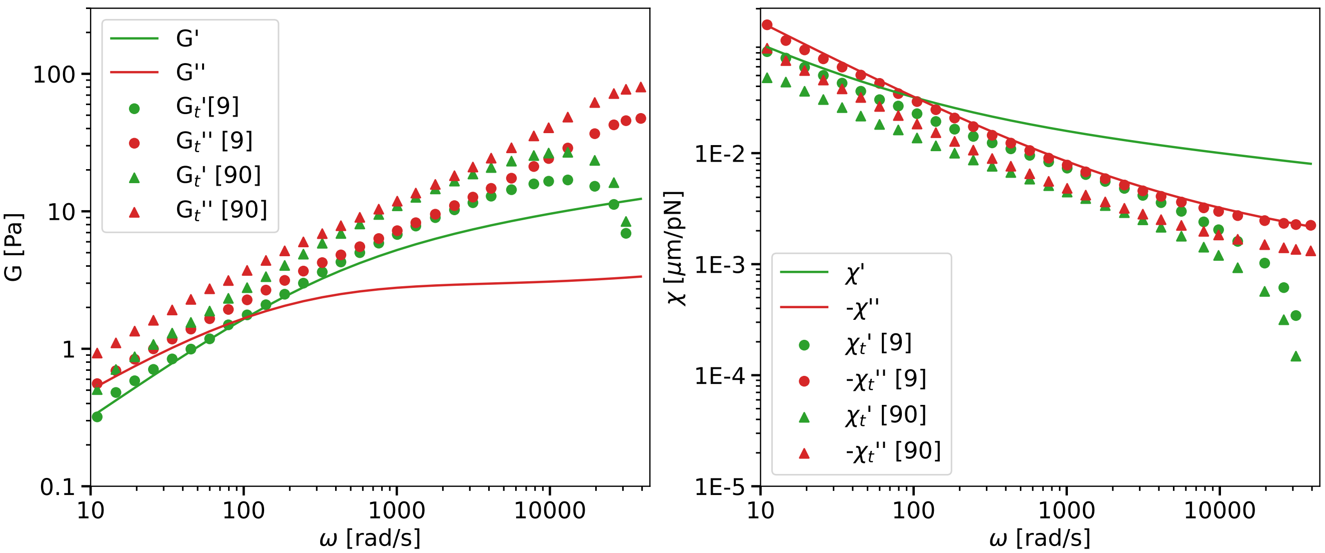

**Supplementary Fig. 7**: FDE prediction of the deviation of the rheological measure in the case of a Maxwell fractional material. The set of model parameters is the same as in Supplementary Fig. 4. The measure obtained with time-sharing: G-modulus $\hat{G}_t(\omega) = G'_t(\omega) + iG''_t(\omega)$ and linear response function $\hat{\chi}_t(\omega) = \chi'_t(\omega)+i\chi''_t(\omega)$. Model behavior: The G-modulus $\hat{G}(\omega) = G'(\omega)+iG''(\omega)$ and the linear response function $\hat{\chi}(\omega) = \chi'(\omega) + i\chi''(\omega)$ are used as references. Time-sharing predictions were calculated for two different values of trap stiffness k: [9] $\rightarrow$ k=9[pN/um]; [90] $\rightarrow$ k=90[pN/um]. The deviations become huge for frequencies above the crossover frequency $\omega_c \approx 100[rad/s]$. Only the values of $\chi''_t(\omega)$ obtained for $k = 9[pN/um]$ remain true to $\chi''(\omega)$.

using the definition given in Eq. 7 for $\hat{\chi}_a(\omega')$. $\hat{x}_l(\omega')$ is the Fourier transform (FT) of the First Harmonic Approximation (FHA) of the trap trajectory which in the time domain $t$ reads

$$x_l(t) = \frac{x_1}{2}\exp(i\omega t)[1+\sin(\omega_t \cdot t)], \tag{28}$$

where the complex oscillation $\exp(i\omega t)$ is used instead of a real one such as $\sin(\omega t)$ or $\cos(\omega t)$ with the sole purpuse of simplifying the notation in the frequency space, which after FT reads

$$\hat{x}_l(\omega') = \frac{x_1}{2}\left[\delta(\omega'-\omega) + \frac{\delta(\omega'-\omega_+) - \delta(\omega'+\omega_-)}{2\,i}\right], \tag{29}$$

with $\omega_\pm = \omega_t \pm \omega$ and where $\delta(\omega')$ is the Dirac distribution.

Supplementary Fig. 8 shows the trap's trajectory in the time domain $x_l(t)$, graph on the left of the figure, and its normalized power spectrum $|2x_l(\omega')/x_1|^2$, on its right, for an excitation frequency of $f = 1000[Hz] \equiv \omega \approx 6283[rad/s]$. From the power spectra it is evident that, in addition to the desired excitation (peak at $6283[rad/s]$), there are two additional peaks located at $\omega_- \approx 72256[rad/s]$, $\omega_+ \approx 84823[rad/s]$, which telecommunications specialists could identify as the power spectrum of a dual-band amplitude-modulated waveform with carrier suppression. These additional peaks are responsible for the deviation of $\hat{\chi}_t$ from $\hat{\chi}$ already observed with the FDE approach.

To get the linear response function $\chi_t(\omega)$ from the FHA approximation the same steps identified for the FDE simulation must be followed but this time using an analytical approach. The details of the calculations are given as Appendix 2. The obtained expression of $\hat{\chi}_t(\omega)$ is a function of the response function of the material $\hat{\chi}$, of the time sharing frequency $\omega_t$, of the trap stiffness $k$ and reads

$$\hat{\chi}_t(\omega) \;=\; g\left(\hat{\chi},\omega_t,k\right)(\omega) = \frac{\hat{\chi}(\omega) - \hat{\chi}_1(\omega) + k\left[\hat{\chi}(\omega)\hat{\chi}_1(\omega) - \hat{\chi}_+(\omega)\hat{\chi}^*_-(\omega)\right]}{1 + 2\,k\hat{\chi}_1(\omega) + k^2\hat{\chi}_+(\omega)\hat{\chi}^*_-(\omega)}, \tag{30}$$

where

$$\hat{\chi}_1(\omega) \;=\; \frac{1}{2}\left[\hat{\chi}_+(\omega) + \hat{\chi}^*_-(\omega)\right], \tag{31}$$

$$\hat{\chi}_+(\omega) \;=\; \hat{\chi}(\omega_t + \omega), \tag{32}$$

$$\hat{\chi}^*_-(\omega) \;=\; \hat{\chi}^*(\omega_t - \omega) = \hat{\chi}(-\omega_t + \omega), \tag{33}$$

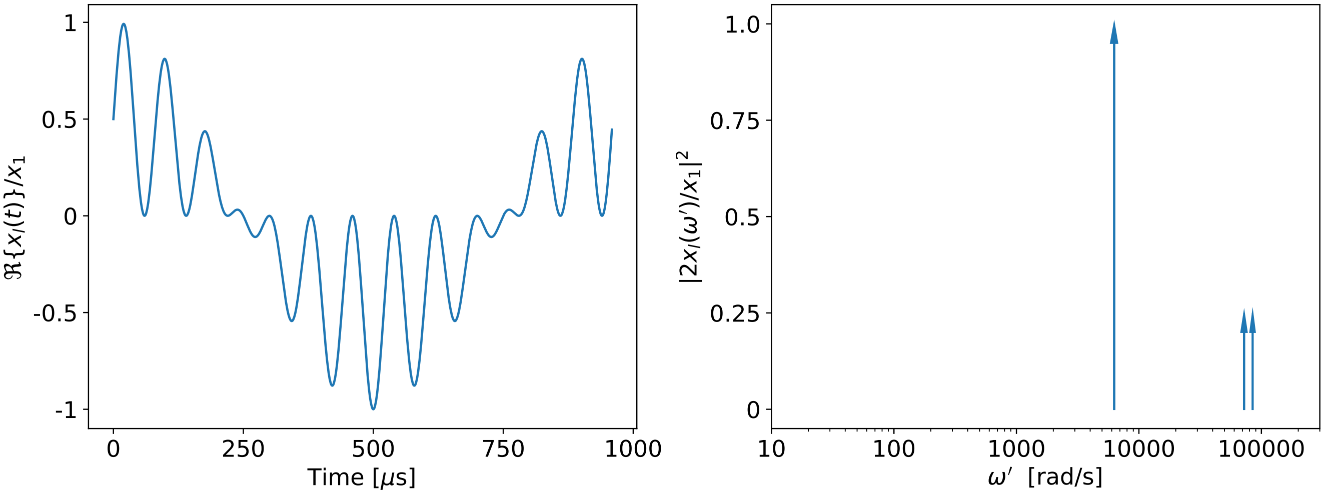

**Supplementary Fig. 8**: FHA of the trajectory of the time-shared trap. The left figure shows the trap trajectory in the time domain $x_l(t)$ for an excitation frequency of $f = 1000[Hz] \equiv \omega \approx 6283[rad/s]$ and a time-sharing frequency $f_t = 12500[Hz] \equiv \omega_t \approx 78539[rad/s]$. The right figure shows the normalized power spectra of the trap trajectory where the following are visible: the wanted excitation peak at $\omega' = \omega$ and the two unwanted additional high-frequency peaks at $\omega' = \omega_+$ and $\omega' = \omega_-$ responsible for the deviation of $\hat{\chi}_t$ with respect to $\hat{\chi}$.

and the asterisk $^*$ denotes the complex conjugate operation.

The results of the Eqs. 30, 31, 32 and 33 show that $\hat{\chi}_t(\omega)$ depends on the response function $\hat{\chi}$ measured at different frequencies. In addition to the contribution at the excitation frequency $\omega$, the contributions from the sidebands of the amplitude-modulated signal at the frequencies $\omega_- = \omega_t - \omega$ and $\omega_+ = \omega_t + \omega$ also determine the behavior of $\hat{\chi}_t(\omega)$ (Supplementary Fig. 8).

Before using Eq. 30 to understand the causes of the deviation, the goodness of the FHA approximation should be checked by comparing the results predicted using Eq. 30 with those predicted using the FDE approach for various model materials (Supplementary Fig. 9 and 9). We can also test the ability of the analytical FHA formalism to predict the deviations introduced by the time-sharing rheology and compare them to the ones obtained from the numerical FDE simulations. As shown in Main Text Figure 1e, the FDE accurately models the experimental time-sharing. Now we can see in Supplementary Fig. 9 that the time-sharing deviation is indeed well described by the FHA. As can be appreciated, the difference between the time sharing response function (thin lines) and the ideal response function of a fractional Kelvin-Voigt material (thick lines) is less than 20% up to 12566 rad/s. Likewise, the FHA matches the FDE with a difference of less then 6%. This means that the FHA can be used to find the ideal response function that

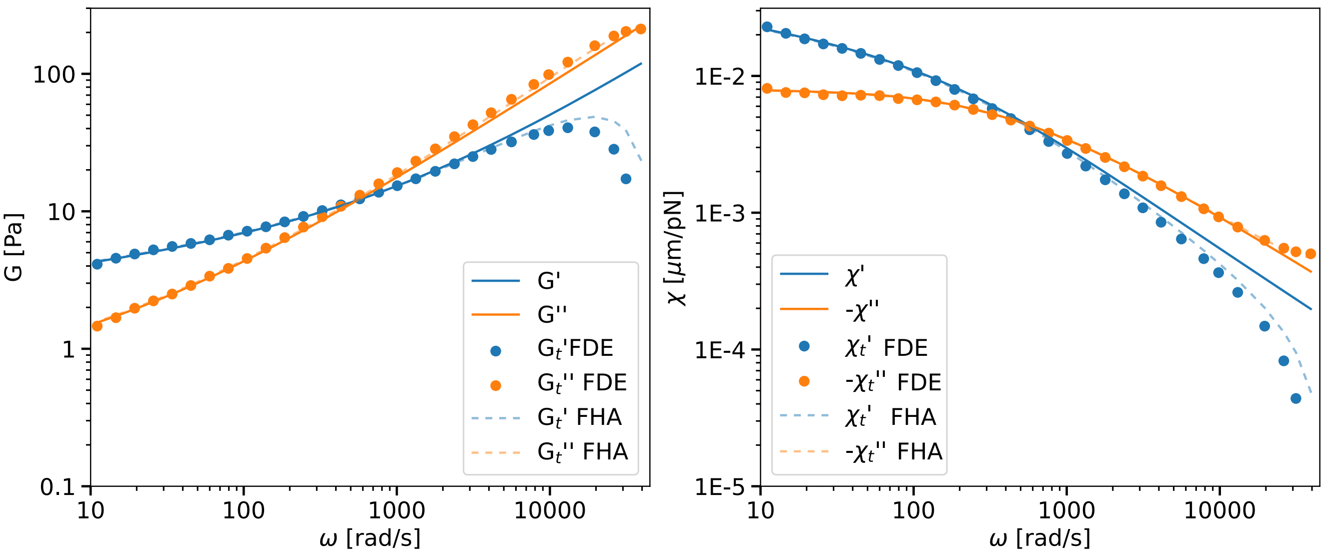

**Supplementary Fig. 9**: Comparison of FDE (circles) and FHA (dashed lines) predictions of the deviation of the rheological measure in the case of a fractional Kelvin-Voigt material (solid lines). The set of model parameters is the same as in Supplementary Fig. 4. The time-shared FDE and FHA predictions were calculated for a trap stiffness k=45[pN/um]. The largest difference between the two predictions is shown in $\chi_t'(\omega)$. However, the difference between $\chi_t'(\omega)$ differs less than 20% for frequencies up to $f = 2000Hz \equiv \omega \approx 12566 rad/s$. Moreover, the difference between the two predictions (FDE vs FHA) of $\chi_t''(\omega)$ differs less than 6% for the entire range of frequencies up to the Nyquist frequency $f_n = f_s/2 = 6250Hz \equiv \omega \approx 39270 rad/s$.

describes the material if we had performed a time-continuous measurement of stress and strain.

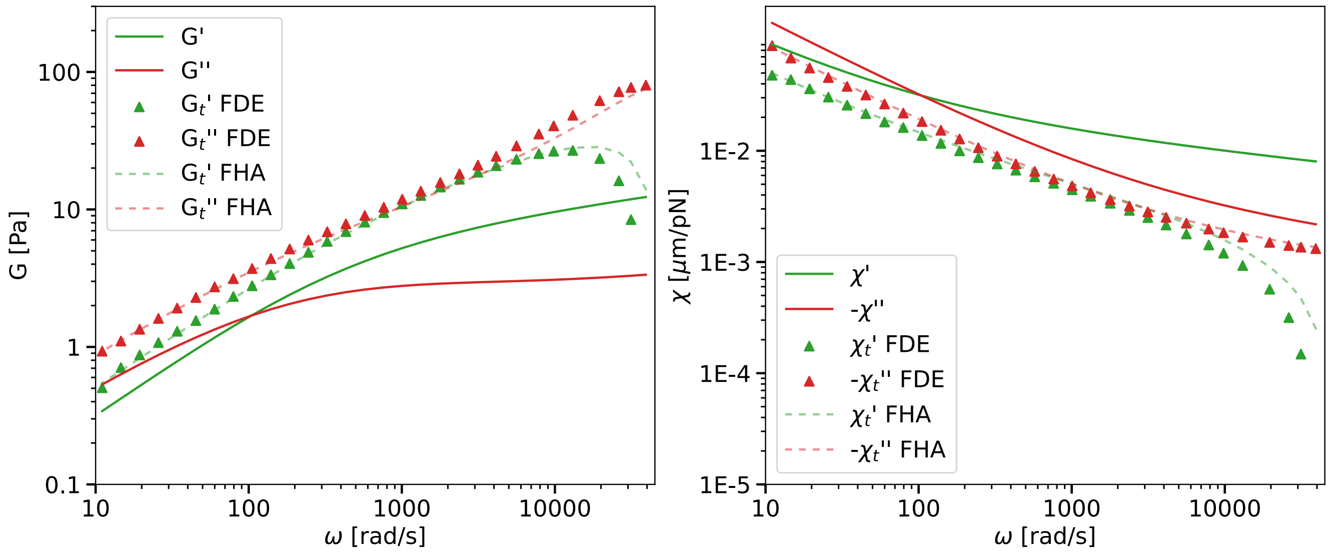

**Supplementary Fig. 10**: Comparison of FDE (triangles) and FHA (dashed lines) predictions of the deviation of the rheological measure in the case of a fractional Maxwell material (solid lines). The set of model parameters is the same as in Supplementary Fig. 4. The time-shared FDE and FHA predictions were calculated for a trap stiffness k=90[pN/um]. The largest difference between the two predictions is shown in $\chi_t'(\omega)$. However, the difference between $\chi_t'(\omega)$ differs less than 30% for frequencies up to $f = 2000[Hz] \equiv \omega \approx 12566[rad/s]$. In contrast, the difference between the two predictions of $\chi_t''(\omega)$ is the smallest and differs less than 7% for the entire range of frequencies up to the Nyquist frequency $f_n = f_s/2 = 6250[Hz] \equiv \omega \approx 39270[rad/s]$.

Supplementary Fig. 10 shows the representative case of fractional Maxwell material, measured with a trap stiffness of $k = 90[pN/um]$, which was the one that showed the largest deviation among the cases already treated using the FDE approach. For this extreme case, the predictions of $\hat{\chi}_t(\omega)$, obtained with the two approaches, FDE and FHA, show that the $\chi_t'(\omega)$ differs less than 30% for frequencies up to $f = 2000[Hz] \equiv \omega \approx 12566[rad/s]$, and, $\chi_t''(\omega)$ differs less than 7% for the whole range of frequencies up to the Nyquist frequency $f_n = f_s/2 = 6250[Hz] \equiv \omega \approx 39270[rad/s]$. This kind of precision on logarithmic behavior such as rheological one is more than acceptable. Moreover, the physics behind the deviation is completely captured by FHA. The difference in prediction between the two approaches could be reduced by taking into account more harmonics of the $\omega_t$ carrier in the $x_l(t)$ representation.

#### 1.9 Understanding the deviation induced by the time-sharing technique

FHA is used to understand the causes of $\hat{\chi}_t(\omega)$ deviation by analyzing the behavior of Eq. 30. For this purpose, $\hat{\chi}_t(\omega)$ is expanded as a first-order Taylor series in k. The result reads

$$\hat{\chi}_t(\omega) \approx g_0(\hat{\chi}, \omega_t)(\omega) + g_1(\hat{\chi}, \omega_t)(\omega) \cdot k + O(k^2), \tag{34}$$

where,

$$g_0(\hat{\chi}, \omega_t)(\omega) = \hat{\chi}(\omega) - \hat{\chi}_1(\omega), \tag{35}$$

$$g_1(\hat{\chi}, \omega_t)(\omega) = 2\hat{\chi}_1(\omega)^2 - \hat{\chi}(\omega)\hat{\chi}_1(\omega) - \hat{\chi}_+(\omega)\hat{\chi}_-^*(\omega). \tag{36}$$

The zero-order term and the first-order term contain different kinds of information, which will be discussed separately.

The zero-order term is the most important because it cannot be avoided. It corresponds to the residual deviation when the trap stiffness $k$ becomes negligible with respect to $1/|\hat{\chi}(\omega_t/2)|$. This is the limit where the material is much stiffer (or viscous, i.e., $(6\pi a)\mu\omega > k$ ) than the trap. In such a case, the force exerted by the driving trap, trap 1, is much higher than that exerted by the static trap, trap 2. As mentioned in subsection 1.3 this is the right regime in which to work to obtain accurate measurements. We will see here that this is the case in most cases, with one exception.

The analysis of $g_0 = g_0' + ig_0''$ begins by using Eq. 31 to express $\hat{\chi}_1$ and examining its real part $g_0'$ and imaginary part $g_0''$ separately, which read

$$g_0'(\omega) = \chi'(\omega) - \frac{1}{2}\left[\chi'(\omega_t + \omega) + \chi'(\omega_t - \omega)\right] = \chi'(\omega) - \chi'(\omega_t) + O(\omega^2) \tag{37}$$

and,

$$g_0''(\omega) = \chi''(\omega) - \frac{1}{2}\left[\chi''(\omega_t + \omega) - \chi''(\omega_t - \omega)\right] = \chi''(\omega) - \omega \left.\frac{d\chi''(\omega')}{d\omega'}\right|_{\omega'=\omega_t} + O(\omega^3). \tag{38}$$

The behavior of the linear response $\hat{\chi}$ at the time-sharing frequency $\omega_t$ determines the zero-order deviation. The real part depends on its value $\chi'(\omega_t)$, and the imaginary part on its derivative $d\chi'(\omega)/d\omega|_{\omega=\omega_t}$.

Consider our two models, the fractional Kelvin-Voigt and the fractional Maxwell, characterized by a crossover frequency $\omega_c$ that is supposed to be located at a much lower frequency than $\omega_t$. Under such an assumption $\hat{\chi}(\omega)$ at a sufficiently high frequency $\omega > \omega_c$, but lower than Nyquist's frequency, i.e., $\omega < \omega_n = \omega_t/2$ behaves like the fractional springpot that reads

$$\hat{\chi}(\omega) \;\approx\; \frac{1}{6\pi a} \cdot \frac{1}{c\,(i\omega)^{\alpha}} \tag{39}$$

where $\alpha = \beta > 0.5$ in the fractional Kelvin-Voigt case, and $\alpha = \alpha < 0.5$ in the fractional Maxwell case. In this context it is convenient to rewrite Eqs. 37 and 38 as

$$\frac{g_0'(\omega)}{\chi'(\omega)} \;\approx\; 1 - \left(\frac{\omega}{\omega_t}\right)^{\alpha} \tag{40}$$

and

$$\frac{g_0''(\omega)}{\chi''(\omega)} \;\approx\; 1 - \alpha\left(\frac{\omega}{\omega_t}\right)^{\alpha+1} \tag{41}$$

where Eq. 39 has been used for $\hat{\chi}(\omega)$ and $\omega$ is considered lower than the Nyquist frequency $\omega_n$. The residual deviation of the imaginary part, Eq.38, it is practically negligible. The worst case is given by $\alpha = 1$ where $\chi_t''(\omega)/\chi''(\omega) = 1 - (\omega/\omega_t)^2$.

On the other hand, the zero-order deviation of the real part, Eq.37, can be large and depends strongly on $\alpha$. It increases when $\alpha$ decreases. For fractional Kelvin-Voigt material, where $\alpha > 0.5$, $\chi'(\omega)$ decreases more rapidly with frequency than for fractional Maxwell material, where $\alpha < 0.5$, and the deviation of the former is smaller than that of the latter. However, even for the Maxwell fractional model, as long as $\alpha > 0$ the deviation introduced by the time-sharing technique does not completely cancel out the information about $\chi'(\omega)$ contained in $\chi_t'(\omega)$.

The failure of the technique occurs when $\alpha = 0$, corresponding to classical Maxwell model. In this case $\chi_t'(\omega)/\chi'(\omega) = 0$. At first glance, any information about the elastic part of the classical Maxwell model is not accessible by the time-sharing technique. The zero-order term of $\hat{\chi}_t(\omega)$ turns out to be effectively degenerate in this case.

However, as we will see below, the first-order term of the expansion Eq. 36 contains information about the elastic part of the Maxwell model and breaks the degeneracy. The linear response function for the classical Maxwell model reads

$$\hat{\chi}(\omega) = \frac{1}{6\pi a}\left[\frac{1}{E} - \frac{i}{\mu\omega}\right], \tag{42}$$

where $E$ is the elastic modulus in [Pa] and $\mu$ is the dynamic viscosity in [Pa·s]. By using Eq. 42 to express the imaginary part $g_1''(\omega)\cdot k$ of the first-order deviation contribution, Eq. 36, it follows that

$$\begin{aligned} g_1''(\omega)\cdot k &= \left[4\chi_1'\chi_1'' - (\chi'\chi_1'' + \chi_1'\chi'') - \left(\chi_+'\chi_-^{*\prime\prime} + \chi_+''\chi_-^{*\prime}\right)\right](\omega)\cdot k \\ &\approx \frac{G_0}{E}\frac{1}{\mu\omega} \end{aligned} \tag{43}$$

where $G_0$ is the scale of the measurement in [Pa], see Eq. 11, and it reads $G_0 = k/12\pi a$. Therefore, as the trap stiffness $k$ changes, the $G_0$ scale of the measurement changes. The apparent viscosity of $\chi_t''(\omega)$ increases proportionally to $k/(12\pi a E)$. This means that by varying $k$ it is possible to access the value of the elastic part $E$ of the classical Maxwell model. The degeneracy of $\hat{\chi}_t(\omega)$ with respect to $E$ is broken if we look at the variation of $\chi_t''(\omega)$ as a function of $k$.

The imaginary part $g_1''(\omega)$ of the first-order term expresses the contribution resulting from the mixing between the real and imaginary parts of the response function that the time-sharing technique generates. Such mixing is proportional to the rigidity of the trap $k$. Usually $k$ must be chosen such that $G_0$ turns out to be smaller than the measured $|G(\omega)|$ in order to obtain an accurate measurement. But in the classical Maxwell case, the measurement at different $G_0$ taken for values of the same order of $E$ allows to break the degeneracy of the time-sharing technique for this kind of samples, enabling the measurement of $E$.

#### 1.10 Introduction to the compensation procedure of the deviation

The micro-rheology method we introduced uses two symmetrical traps that act simultaneously on the bead. The first trap acts as a driving trap, while the second one acts as a detection laser. The fact that the two traps have the same stiffness $k/2$ causes the sensitivity calibration $\beta$ to be cancelled by symmetry and the only dimensional factor of the rheological method is $G_0$ which depends on $k$ and the radius of the bead $a$. The two traps should act simultaneously on the bead for the method to work properly. The time-sharing technique makes it possible to generate two identical traps, but it cannot satisfy the criterion of simultaneity. The latter fact causes the results obtained with this technique to be affected by a deviation that has been studied in the previous sections. In this section we would like to see under what conditions it is possible, from the data measured with the time-sharing technique, to answer the counterfactual question: What would we have measured with our rheological method without having violated the simultaneity criterion?

To answer this counterfactual question, several conditions must be true. First, we must have a sufficiently good understanding of the origin of the deviation induced by the time-sharing technique. Second, that this understanding passes through a mathematical description of the deviation that can accurately quantify it. Third, that the relationship existing between the set $\mathbb{B}$ of the response functions $\hat{\chi}(\omega)$ and the set $\mathbb{D}$ of the measured by the time-sharing technique $\hat{\chi}_t(\omega)$ is a bijective function $g(\hat{\chi}) : \mathbb{B} \to \mathbb{D}$. Only under these conditions is it possible to invert the function and obtain $\hat{\chi} = g^{-1}(\hat{\chi}_t)$

Having said that, one might think that all the conditions necessary to compensate for the deviation have been mentioned. But they have not. There is another condition that is the most important from a practical point of view. The fourth condition is that all possible confounding factors that might generate a false association between results and sample properties be eliminated. There are many undesirable phenomena that can act as confounding factors. Asymmetry in the duration of the trap cycle, sampling delays, cross-talking between measurement signals, inaccuracy in trap placement-all these phenomena must be reduced as much as possible in order to obtain reliable measurements with respect to a larger number of characterizable samples.

Let us begin by answering to the list of conditions we mentioned above. The first and second conditions were discussed in the previous section 1.9. The FHA results are used to discuss the third condition. To prove that the function $g(\hat{\chi})$ is bijective, we need to show that it is injective, i.e., that $\forall\ \hat{\chi}_a,\ \hat{\chi}_b \in \mathbb{B}$ so that $g(\hat{\chi}_a) = g(\hat{\chi}_b) \implies \hat{\chi}_a = \hat{\chi}_b$. To discuss this, we begin by considering the zero-order expansion of $g(\hat{\chi}, k)$ in k, that is, $g_0(\hat{\chi})$ already introduced with the Eq. 35. $\forall\ \hat{\chi}_a$, $\hat{\chi}_b \in \mathbb{B}$, it is possible to write $\hat{\chi}_b = \hat{\chi}_a + \hat{f}$ where $\hat{f} \in \mathbb{B}$ it is also a response function $\hat{f}(\hat{\omega})$ that is bounded, infinitely differentiable and smooth. Therefore, using the Eq. 37 up to the second order in $\omega$

$$
\begin{aligned}
g_0'(\hat{\chi}_b)(\omega) &= \chi_b'(\omega) - \chi_b'(\omega_t) - \frac{\omega^2}{2}\left.\frac{d^2\chi_b'}{d\omega^2}\right|_{\omega=\omega_t} + O(\omega^4) \\
&= \chi_a'(\omega) - \chi_a'(\omega_t) - \frac{\omega^2}{2}\left.\frac{d^2\chi_a'}{d\omega^2}\right|_{\omega=\omega_t} \\
&+ f'(\omega) - f'(\omega_t) - \frac{\omega^2}{2}\left.\frac{d^2 f'}{d\omega^2}\right|_{\omega=\omega_t} + O(\omega^4),
\end{aligned}
\tag{44}
$$

and using Eq. 38 for the imaginary part,

$$
\begin{aligned}
g_0''(\hat{\chi}_b)(\omega) &= \chi_b''(\omega) - \omega\left.\frac{d\chi_b''}{d\omega}\right|_{\omega=\omega_t} + O(\omega^3) \\
&= \chi_a''(\omega) - \omega\left.\frac{d\chi_a''}{d\omega}\right|_{\omega=\omega_t} + f''(\omega) - \omega\left.\frac{df''}{d\omega}\right|_{\omega=\omega_t} + O(\omega^3),
\end{aligned}
$$

it follows that the equality $g(\hat{\chi}_a) = g(\hat{\chi}_b)$ implies $f'(\omega) - f'(\omega_t) - \frac{\omega^2}{2}\left.\frac{d^2 f}{d\omega^2}\right|_{\omega=\omega_t} + O(\omega^4) = 0$ and $f''(\omega) - \omega\left.\frac{df''}{d\omega}\right|_{\omega=\omega_t} + O(\omega^3) = 0$. The only nontrivial solution of these equations is $f(\omega) = A + iB\omega$ with A and B $\in \mathbb{R}$. However, the imaginary part $i\omega B$ is unbounded and must be discarded because does not belong to the set of response function $\mathbb{B}$. Then the only valid solution is $f(\omega) = A$ with A $\in \mathbb{R}$. The physical meaning of $A$ is that of an elastic component $A = 1/(6\pi a E)$ where E is the elastic modulus in [Pa]. Then from our calculus we found that the two response functions $\hat{\chi}_a(\omega)$ and $\hat{\chi}_b(\omega) = \hat{\chi}_a(\omega) + 1/(6\pi a E)$ gives the same result $g_0(\hat{\chi}_b) = g_0(\hat{\chi}_a)$. So we can affirm that the zero-order function $g_0(\hat{\chi})$ is not injective. This is the generalisation of the degeneracy problem of the classical Maxwell model treated in the section 1.9, where we found that we cannot access the value of the elastic modulus E.

The result obtained shows that all types of response functions that have a classical elastic element in series that can be described as $\hat{\chi}_b(\omega) = \hat{\chi}_a(\omega) + 1/(6\pi a E)$ are indistinguishable from $\hat{\chi}_a(\omega)$ using the time-sharing technique up to the zero order in k. However, if we define the subset of the response function set $\mathbb{B}$, $\mathbb{B}^- = \{\forall\hat{\chi} \in \mathbb{B} | \lim_{\omega\to\infty} |\hat{\chi}(\omega)| = 0\}$ as domain and $\mathbb{D}^0 = \{\hat{\chi}_t = g_0(\hat{\chi}) | \forall\hat{\chi} \in \mathbb{B}^-\}$ as codomain of the function $g_0(\hat{\chi}) : \mathbb{B}^- \to \mathbb{D}^0$, $g_0(\hat{\chi})$ becomes bijective and then invertible.

This is an important result because in the practice of microrheology, samples belonging to $\mathbb{B} - \mathbb{B}^-$ are rare and the most interesting samples are found in $\mathbb{B}^-$, especially in the subset of fractional response functions. The deviation for these samples is compensated by the inversion of $g_0$. That is, by finding a way to calculate $\hat{\chi} = g_0^{-1}(\hat{\chi}_t)$.

It is important to emphasize again the fact that for all the samples it is convenient to make the measurements with a trap that is much less rigid than the material behaviour at the Nyquist frequency. That is, one must choose a $k << 1/|\hat{\chi}(\omega_t/2)|$ or equivalently $G_0 << |\hat{G}(\omega_t/2)|$. So in the worst case it will be necessary to make several iterations to find the optimal stiffness of the trap $k$ with which to make the measurements.

Having correctly chosen $k$ the only relevant term of $g$ is the zero-order component $g_0$ and thus what has been discussed so far applies. Otherwise, the behavior of at least the first term $g_1$ must be taken into account and, in the case of considering measurements taken at a single value of $k > 1/|\hat{\chi}(\omega_t/2)|$, one might obtain that $g$ is no longer a bijective function even for the subset $\mathbb{B}^-$. This has not been proven, but it could be the case because of the presence of quadratic terms in $\hat{\chi}$ and is therefore best avoided.

However, as discussed in the previous chapter for the case of the classical Maxwell model, see Eq. 43, the first-order term $k \cdot g_1(\chi)$ contains information about the elastic component $E$ that could be connected in series with the response function $\hat{\chi}(\omega) \in \mathbb{B}^-$ determined using the inversion of the zero-order term $g_0(\hat{\chi})$ as just described. A simple method to determine the presence of $E$ is to postulate an extension of the measured response function $\hat{\chi}(\omega, E) = \hat{\chi}(\omega) + 1/(6\pi a E) \in \mathbb{B}$. Then, by introducing $\hat{\chi}(\omega, E)$ into the FHA Eq. 30 and comparing the results with those obtained from a series of measurements of $\hat{\chi}_t(\omega)$ for different $k$, it is possible to find $E$ by variational optimization.

Before describing the compensation procedure of the results in detail, the next chapter is dedicated to discuss the fourth condition mentioned above: the identification and elimination of possible confounding factors.

#### 1.11 Testing the time-sharing technique on known samples

As mentioned above, the practical implementation of the time-sharing technique can be affected by a multitude of confounding factors that can compromise its performance. To verify the proper implementation of the technique, we tested it with known samples so that the measurements obtained by time-sharing could be compared with the predictions generated by the FDE and FHA approaches before applying the compensation procedure (see also Extended Data Fig. 2). We chose pure water, glycerol/ewater mixtures, different poly-acryl amide gels and uncured PDMS as models.

The Supplementary Fig. 11 shows several measurements of $\hat{G}_t(\omega)$ for pure water, made using the time-sharing technique implemented in the SENSOCELL optical tweezers platform. The left plot shows three different measurements made at $k$=114[pN/um] and $r = 0.5[\mu m]$, while the right plot shows three more measurements obtained for $k$=12[pN/um] and $r = 0.5[\mu m]$. The measurement scale of the left graph is set to $G_0 = k/(12\,\pi\,r) = 6[Pa]$ and that of the right graph to $G_0 = 0.31[Pa]$.

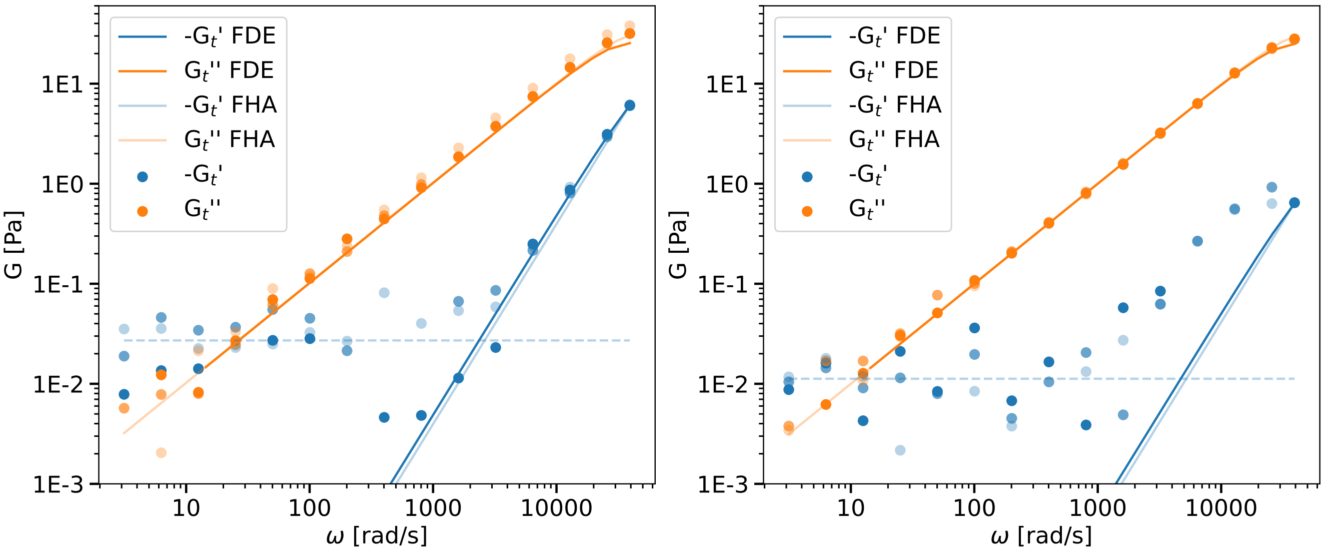

**Supplementary Fig. 11**: $\hat{G}_t(\omega)$ for pure water obtained using two different measurement scales $G_0 = k/(12\,\pi\,r)$. The left-hand graph shows the case where $G_0 = 6[Pa]$ and the right-hand graph the case where $G_0 = 0.31[Pa]$. Each graph shows three different measurements, indicated with dots of different shades. The solid lines show the predictions provided by the FDE simulation and the FHA approximation. The discontinuous blue lines show the level of the mean value of the noise bottom affecting the real component $G'$.

The different measurements are indicated with dots of different shades, while the solid lines show the predictions provided by the FDE simulation and the FHA approximation for a Newtonian liquid of viscosity $\mu = 0.001[Pa.s]$.

The measured imaginary component $G''_t(\mu)$, orange dots, of the right-hand graph fit the theoretical predictions, orange lines, better than the left-hand graph. This is because the measurement scale of the left-hand graph, $G_0 = 6[Pa]$, is larger than the magnitude of $G''_t(\omega)$ for almost the entire range of measurable frequencies (up to $\omega = 6000[rad/s]$) and, as already explained in the chapter 1.3, this compromises the precision of the measurement. In the case of the graph on the right, the measurement scale, $G_0 = 0.31[Pa]$, is smaller than the magnitude of $G''_t(\omega)$ for frequencies greater than $\omega = 310[rad/s]$ and the precision of the measurement is preserved for a wide range of frequencies.

The explanation with respect to the behaviour of the real components of $G'_t(\omega)$ is more complex. Several factors influence the results. The high-frequency behaviour of the mesurations, blue dots, of the graph on the left are the only ones that fit the predictions from about $\omega = 1000[rad/s]$. In other cases, the predictions, the blue lines, are at lower values than the experimental data. Two distinct phenomena can be identified.

On the one hand, the experimental results show the appearance of a noise bottom whose average value depends on the value of the measurement scale $G_0$ and is shown on the graphs by discontinuous blue lines. Its magnitude is $0.03[Pa]$, for the left-hand graph, and $0.01[Pa]$, for the right-hand graph. This bottom on the real part is due to the resolution of the instrument. As already explained in the chapter 1.3, when the scale $G_0$ is much larger than $|\hat{G}(\omega)|$ the forces exerted by the two traps, trap 1 and trap 2, are practically identical in magnitude, but have opposite orientations. The result is that $||V_2| - |V_1||$ should continuously decrease to 0. But a noise bottom of the order of the resolution of the analogue-to-digital converter $V_{res}$ limits its minimum value.

On the other hand, the ratio between the real and imaginary components, $G'_t(\omega)/G''_t(\omega)$, is limited by the relative phase precision between $V_1$ and $V_2$, which is of the order of a few percent. Values of $G'_t(\omega)$ for which this ratio would be less than the phase error are not achievable.

The values of $\hat{G}_t(\omega)$ obtainable with water in the frequency range of the instrument are limited to a few Pascals. However, the rheological method studied here was developed to perform microrheology measurements in samples with $\hat{G}_t(\omega)$ of the order of $1 \cdot 10^3[Pa]$ without having to use an additional detection laser. In this regime the ratio of $V_1/V_2$ is greater than 100 and a cross-talk $c_t$ of the order of 1% can drastically change the measurement result. The simplest model to describe the cross-talk between $V_1$ and $V_2$ reads

$$\begin{aligned} V_{1,c}(t_{1,i}) &= V_1(t_{1,i}) + c_t \cdot V_2(t_{2,i-1}) \\ V_{2,c}(t_{1,i}) &= c_t \cdot V_1(t_{1,i}) + V_2(t_{2,i}) \end{aligned} \tag{45}$$

where $V_{1,c}$, $V_{2,c}$ are the resulting modified signals. The physical sources of the cross-talk can be electrical, e.g. due to a slow sensor response, or result from poor trap placement caused by acoustic coupling in AODs.

To test the ability of the method and instrument to measure $\hat{G}_t(\omega)$ values larger than $1 \cdot 10^3 [Pa]$, a mixture of 20% water by volume and 80% glycerol was used. The dynamic viscosity of the mixture is approximately 60 times that of water, with a dynamic viscosity of $\mu = 0.06[Pa.s]$, and a $\hat{G}''(\omega) \approx 1 \cdot 10^3 [Pa]$ should be reachable for $\omega \approx 17000[rad/s]$.

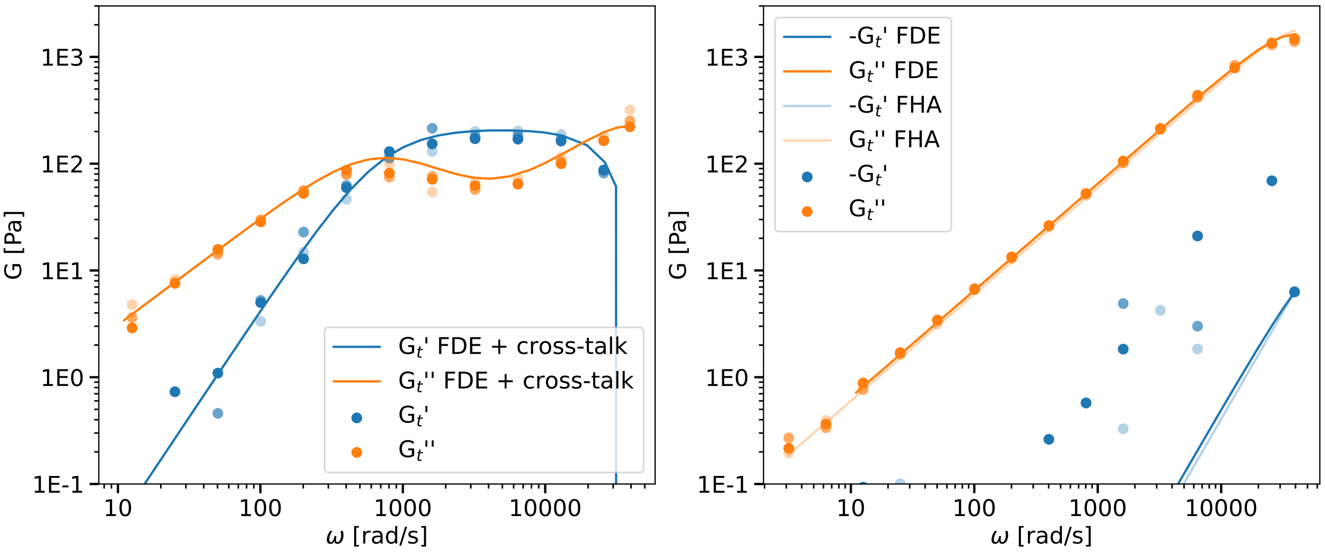

**Supplementary Fig. 12**: $\hat{G}_t(\omega)$ for a mixture of 20% water by volume and 80% glycerol obtained with a measurement scale of $G_0 = 6[Pa]$. Each graph shows three different measurements, indicated with dots of different shades. The left-hand graph shows the case where the instrument is affected by cross-talk estimated at $c_t = -3\%$. The right-hand graph, on the other hand, shows the result obtained from an optimized instrument which as been purged of the main cross-talk sources. The solid lines show the predictions provided by the FDE simulation or FHA approximation. The FDE simulation used to fit the left-hand data include the cross-talk by using Eq. 45.

Supplementary Fig. 12 shows different measurements of $\hat{G}_t(\omega)$ for such a mixture. Both graphs are obtained using a measurement scale $G_0 = 6[Pa]$. The left graph shows the measurement result for an instrument affected by a cross-talk $c_t = -3\%$. The right-hand graph, on the other hand, shows the result obtained from an instrument purged of the main crosstalk sources, where in addition the residual crosstalk of $c_t = 0.17\%$ has been mathematically compensated.

The measured imaginary component $G''_t(\mu)$, orange dots, of the right-hand graph fits perfectly with the theoretical predictions, orange lines, reaching the value $1 \cdot 10^3 [Pa]$ close to the predicted frequency.

The left-hand graph, on the other hand, shows that $G''_t(\mu)$ stays below $1 \cdot 10^3 [Pa]$ in the case where cross-talk is present. Furthermore, the real component $G'_t(\mu)$, assumes a behaviour that might suggest that a sample of polymer solution is being measured and that at high frequency the rubbery plateau is being observed. This is clearly false and shows the importance of minimising all possible sources of cross-talk that could lead to false interpretations of experimental results. Particularly for those from samples with high $\hat{G}_t(\omega)$ values.

We also performed additional experiments in uncured PDMS. We found the expected Maxwell-like behavior and a stiffness that matches previous accounts[12]. For a detailed discussion of the limits of the techniques, please see Section 5.2 below.

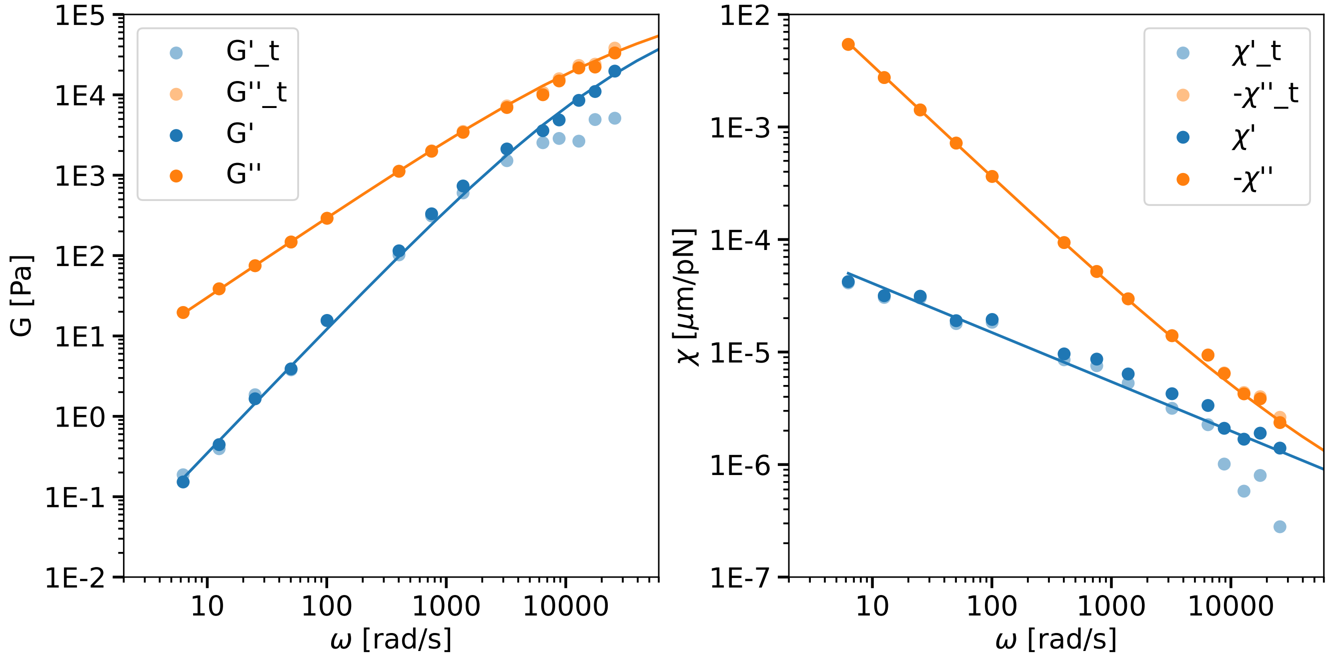

**Supplementary Fig. 13**: PDMS 1% curing agent trap stiffness k=309[pN/$\mu$m]

#### 1.12 The compensation procedure

In the chapter 1.10, we showed that if the response function $\hat{\chi}(\omega)$ of the studied sample belongs to the set $\mathbb{B}^- = \{\forall\hat{\chi} \in \mathbb{B} | \lim_{\omega\to\infty} |\hat{\chi}(\omega)| = 0\}$ and the measurement scale $G_0$ is smaller than $\hat{G}(\omega_t/2)$ then the mapping between response function $\hat{\chi}(\omega)$ and the time-sharing response function $\hat{\chi}_t(\omega) = g\left(\hat{\chi}, \omega_t, k\right)(\omega)$ is invertible. That guarantees the existence of only one element $\hat{\chi}(\omega) \in \mathbb{B}^-$ associated with a given time-sharing response function $\hat{\chi}_t(\omega)$.

We have also seen that, once $\hat{\chi}(\omega)$ has been determined, it is possible to postulate an extension of the response function $\hat{\chi}(\omega, E) = \hat{\chi}(\omega) + 1/(6\pi E)$ to enlarge the set of materials that can be investigated by the method to $\mathbb{B}$. In such a case, a series of measurements of $\hat{\chi}_t(\omega)$ for different $k$ will be required to determine $E$.

The purpose of the compensation procedure is to answer the counterfactual question: What would we have measured with our rheological method if we had not violated the simultaneity criterion? To answer this question for every single measured point of $\omega_i$, Eq. 30 is partially inverted. The result reads

$$\hat{\chi}(\omega_i) = \frac{\hat{\chi}_t(\omega_i)\left[1 + 2\,k\hat{\chi}_1(\omega_i) + k^2\hat{\chi}_+(\omega_i)\hat{\chi}_-^*(\omega_i)\right] + \hat{\chi}_1(\omega_i) + k\hat{\chi}_+(\omega_i)\hat{\chi}_-^*(\omega_i)}{1 + k\hat{\chi}_1(\omega_i)}, \tag{46}$$

from which we obtain the compensated measure $\hat{\chi}(\omega_i)$ as a function of the time-sharing measure $\hat{\chi}_t(\omega_i)$ and the contributions of the first harmonic $\hat{\chi}_+(\omega_i)$ and $\hat{\chi}_-(\omega_i)$. These in turn depend on the behaviour of $\hat{\chi}(\omega)$ near the time-sharing frequency $\omega_t$ and must be evaluated. It is not possible to derive their value directly from the compensated measurements because the highest value of $\omega_i$ is equal to the Nyquist frequency $\omega_t/2$. Hence, $\hat{\chi}_1(\omega)$, $\hat{\chi}_+(\omega_i)$ and $\hat{\chi}_-(\omega_i)$ must be deduced by extrapolating the behaviour of $\hat{\chi}(\omega)$ to frequencies beyond the measurement range. Many procedure are possible. The one we chosed is based on the utilisation of models to parametrise the $\mathbb{B}^-$ set of response function. Once chosed the right model $\hat{\chi}_m$ between the set of avaible model, $\hat{\chi}_1(\omega)$, $\hat{\chi}_+(\omega_i)$ and $\hat{\chi}_-(\omega_i)$ will be evaluated by using it. Explicitely

$$\hat{\chi}_1(\omega) = \frac{1}{2}\left[\hat{\chi}_+(\omega) + \hat{\chi}_-^*(\omega)\right], \quad (47)$$

$$\hat{\chi}_+(\omega) = \hat{\chi}_m(\omega_t + \omega), \quad (48)$$

$$\hat{\chi}_-(\omega) = \hat{\chi}_m(\omega_t - \omega), \quad (49)$$

Here we give a list of the most used models up to 2nd order. For models combining more than two spring-pots, the equivalent expressions for $\hat{\chi}_m$ can be used:

- The Newtonian fluid:

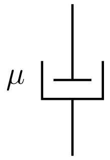

$$\hat{\chi}_m(\omega) = \frac{1}{6\pi a}\frac{1}{i\mu\omega} \text{ with } \mu \in \mathbb{R}^+ \quad (50)$$

- Single power law:

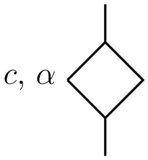

$$\hat{\chi}_m(\omega) = \frac{1}{6\pi a}\frac{1}{c(i\omega)^\alpha} \text{ with } c \in \mathbb{R}^+,\ \alpha \in ]0,1] \quad (51)$$

The single power law model was defined to exclude the pure elastic model $\alpha = 0$, which does not belong to $\mathbb{B}^-$, and to include the Newtonian fluid model, which does belong to $\mathbb{B}^-$. The simplest fractional model that includes the two models introduced so far involves a single power law element $c, \alpha$. In this sense we will call these models first-order models. Following the same logic, we define the second-order models belonging to $\mathbb{B}^-$.

- Kelvin-Voigt:
- Structural damping:
- fractional Kelvin-Voigt:

  with $C_\alpha$ , $C_\beta \in \mathbb{R}^+$, $\alpha$, $\beta \in [0,1]$ and $\alpha \neq \beta$.

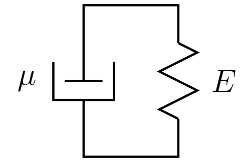

$$\hat{\chi}_m(\omega) = \frac{1}{6\pi a}\frac{1}{E + i\mu\omega} \text{ with } E,\, \mu \in \mathbb{R}^+ \tag{52}$$

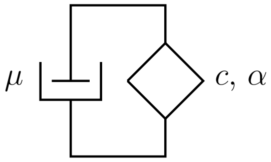

$$\hat{\chi}_m(\omega) = \frac{1}{6\pi a}\frac{1}{c(i\omega)^{\alpha} + i\mu\omega} \text{ with } c \in \mathbb{R}^+,\, \alpha \in [0, 1] \tag{53}$$

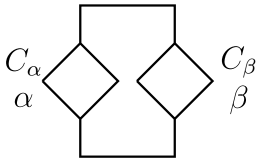

$$\hat{\chi}_m(\omega) = \frac{1}{6\pi a}\frac{1}{C_\alpha(i\omega)^{\alpha} + C_\beta(i\omega)^{\beta}} \tag{54}$$

- Fractional Maxwell:

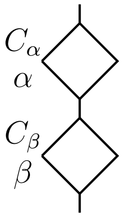

$$\hat{\chi}_m(\omega) = \frac{1}{6\pi a}\left[\frac{1}{C_\alpha(i\omega)^{\alpha}} + \frac{1}{C_\beta(i\omega)^{\beta}}\right] \tag{55}$$

with $C_\alpha$ , $C_\beta \in \mathbb{R}^+$ and $\alpha,\, \beta \in (0, 1]$

The fractional Kelvin-Voigt model includes the Structural dumping and Kelvin-Voigt models. On the other hand, the fractional Maxwell model has been defined here with $\alpha$ and $\beta$ different from zero, so as to exclude the Maxwell model that belongs to $\mathbb{B} - \mathbb{B}^-$. The third-order models are all included in four fractional models which we will call Kelvin-Maxwell-fractional (KMF), Maxwell-Kelvin-fractional (MKF), Kelvin-Kelvin-fractional (KKF) and Maxwell-Maxwell-fractional (MMF). Their definitions follow

- Kelvin-Maxwell-fractionals (KMF):

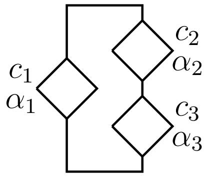

$$\hat{\chi}_m(\omega) = \frac{1}{6\pi a}\frac{1}{c_1(i\omega)^{\alpha_1} + \frac{1}{\frac{1}{c_2(i\omega)^{\alpha_2}} + \frac{1}{c_3(i\omega)^{\alpha_3}}}} \tag{56}$$

with $c_1$ , $c_2$, $c_3 \in \mathbb{R}^+$ , $\alpha_1$, $\alpha_2$, $\alpha_3 \in [0, 1]$ and $\alpha_2 \neq \alpha_3$.

- Maxwell-Kelvin-fractional (MKF):

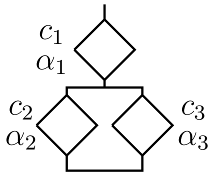

$$\hat{\chi}_m(\omega) = \frac{1}{6\pi a}\left[\frac{1}{c_1(i\omega)^{\alpha_1}} + \frac{1}{c_2(i\omega)^{\alpha_2} + c_3(i\omega)^{\alpha_3}}\right] \tag{57}$$

with $c_1$ , $c_2$, $c_3 \in \mathbb{R}^+$ , $\alpha_1 \in ]0, 1[$, $\alpha_2$, $\alpha_3 \in [0, 1]$ and $\alpha_2 \neq \alpha_3$.

- Kelvin-Kelvin-fractional (KKF):

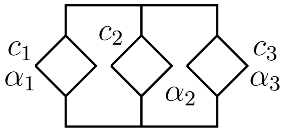

$$\hat{\chi}_m(\omega) = \frac{1}{6\pi a}\frac{1}{c_1(i\omega)^{\alpha_1} + c_2(i\omega)^{\alpha_2} + c_3(i\omega)^{\alpha_3}} \tag{58}$$

with $c_1$ , $c_2$, $c_3 \in \mathbb{R}^+$ , $\alpha_1$, $\alpha_2$, $\alpha_3 \in [0, 1]$ and $\alpha_1 \neq \alpha_2 \neq \alpha_3$.

- Maxwell-Maxwell-fractional (MMF):

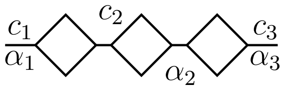

$$\hat{\chi}_m(\omega) = \frac{1}{6\pi a}\left[\frac{1}{c_1(i\omega)^{\alpha_1}} + \frac{1}{c_2(i\omega)^{\alpha_2}} + \frac{1}{c_3(i\omega)^{\alpha_3}}\right] \tag{59}$$

with $c_1$ , $c_2$, $c_3 \in \mathbb{R}^+$ , $\alpha_1$, $\alpha_2$, $\alpha_3 \in ]0, 1]$ and $\alpha_1 \neq \alpha_2 \neq \alpha_3$.

The KMF model includes the following models: the Zener KM or also known as the Maxwell representation of the standard linear solid model ($\alpha_1 = 0$ ,$\alpha_2 = 0$, $\alpha_3 = 1$), the anti-Zener KM or also known as the Maxwell representation of the standard linear liquid model ($\alpha_1 = 1$ , $\alpha_2 = 0$, $\alpha_3 = 1$) which have not been explicitly stated to save space. On the other hand, the set of parameters of the MKF model has been restricted to $\alpha_1 \neq 0$ to exclude the Kelvin representation of the standard solid model ($\alpha_1 = 0$ ,$\alpha_2 = 0$, $\alpha_3 = 1$) which belongs to $\mathbb{B} - \mathbb{B}^-$. The MK anti-Zener model, also known as Jeffreys fluid, is instead included in the MKF model ($\alpha_1 = 1$ ,$\alpha_2 = 0$, $\alpha_3 = 1$).

The KKF and MMF models are extensions to third-order models of the Maxwell-fractional and Kelvin-fractional second-order. The latter two models are rarely used due to the reduced frequency range that can be explored with the time-sharing method. In fact, normally to limit the measurement to a few minutes, the measurements made concern the range from 1 [rad/sec] to 10000 [rad/sec]. For this frequency range, the Kelvin-fractional and Maxwell-fractional second-order models are more than sufficient. Given this list of models to be used to parameterize the set $\mathbb{B}^-$ of response functions explorable by our method the following question arises: How to choose the right model to performe the compensation given a set of experimental measurement $\hat{\chi}_t(\omega_i)$? First of all, the measures must fulfill two prerequisites. The first is that they must be precise enough to show a clear trend as a function of frequency. The second is that they must extend to high frequencies, preferably up to $f = 2000[Hz] \equiv \omega \approx 12566[rad/s]$, for the extrapolation of the behavior of the response function close to the time-sharing refreshing frequency $f_s = 12500[Hz] \equiv \omega \approx 78539[rad/s]$ to be accurate.

That said, it is possible to identify the model to choose by observing the behavior of the imaginary part of the measured time-sharing response function $\hat{\chi}''_t(\omega_i)$ as a function of frequency. As seen in section 1.7 the deviation affecting $\hat{\chi}''_t(\omega_i)$ is small and considering its behavior not beyond the frequency $f = 2000[Hz] \equiv \omega \approx 12566[rad/s]$, it can be assumed, for this specific purpose, that $\hat{\chi}''_t(\omega_i) \approx \hat{\chi}''(\omega_i)$. The behavior of $\hat{\chi}''_t(\omega_i)$ holds the signature of the model to be chosen $\hat{\chi}_m(\omega)$. Supplementary Fig. 24 shows the three most important patterns for $-\hat{\chi}''_t(\omega_i)$ in log-log representation. If $-\hat{\chi}''_t$, always in the log-log representation, follows a negative slope line, then the model to choose is the single power law. This is the case with the green data marked PL. If the slope turns out to be equal to -1 then it is a Newtonian fluid. If, on the other hand, $-\hat{\chi}''_t$ turns out to be a concave function, i.e., that its slope is increasingly negative as frequency increases, then it is a Kelvin fractional or one of its children: Structural dumping or Kelvin-Voigt. This is the case with the orange data marked KF. Finally if $-\hat{\chi}''_t$ is convex, that is, its slope is less negative as frequency increases then we are dealing with a Maxwell fractional sample. This case is shown in red and is labeled MF.

Once the model type for $\hat{\chi}_m(\omega)$ has been selected, its parameters $p = \{c_j, \alpha_j\}$ have to be fitted to the data by minimizing the sum of the relative error between the measured data $\hat{\chi}_t(\omega_i)$ and its FHA predictions $g(\chi_m, \omega_i, p)$, see Eq. 30. More explicitly, the parameter set $p$ is chosen to minimize the following functional

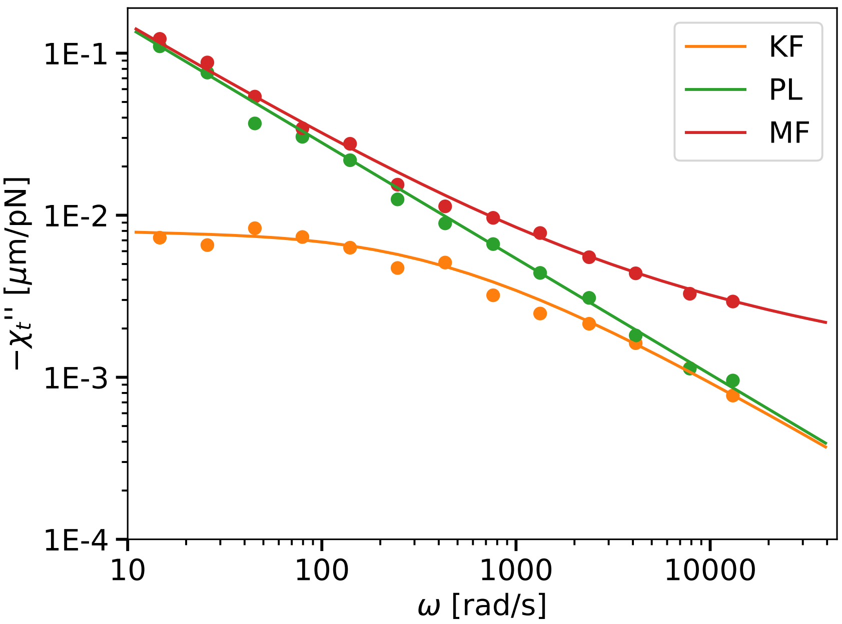

**Supplementary Fig. 24**: The three most important patterns for $-\hat{\chi}_t''(\omega_i)$ in a log-log graph. The PL data set shows the signature of a first-order, power law or Newtonian fluid model: a straight line. The KF data set shows the signature of Kelvin fractional model: a concave function. The MF data set shows the signature of Maxwell fractional model: a convex function.

$$E(p) = \sum_i \log(|\chi_t'(\omega_i)/g'(\hat{\chi}_m, \omega_i, p)|)^2 + \log(|\chi_t''(\omega_i)/g''(\hat{\chi}_m, \omega_i, p)|)^2. \tag{60}$$

Having found the parameter set $p$, the model can be used to evaluate $\hat{\chi}_1(\omega)$, $\hat{\chi}_+(\omega_i)$ and $\hat{\chi}_-(\omega_i)$ by using the equations 47, 48 and 49 and the answer to our counterfactual question, 'What would we have measured with our rheological method if we had not violated the simultaneity criterion?' is obtained straightforwardly by evaluating Eq. 46.

#### 1.13 Concluding remarks.

In the preceding sections we have learned how time-sharing and its implementation lead to controlled deviations from the ideal, time continuous expectations. In summary, we have two overlapping aspects:

1. The measurement errors that may arise due to the method based on two laser beams with

same power (driving and static). This errors are higher at low frequencies where $|V1|$ is about $|V2|$ as pointed out by the referee and required that $G >> G_0$.

- This aspect may make the measurement inaccurate at low frequencies, particularly if you are going to measure a very soft material where $|V1|/(|V1|-|V2|) >> 1$ (e.g. 1000 times larger), but it does not affect the accuracy of the measurement at high frequencies
- As we have shown in Supplementary Fig 6, varying the trap stiffness 10 times, does not lead to variations in the expected moduli.

2. The measurement deviation due to the intermittency of the laser beams (the time-sharing).

   - The deviation only affects the highest frequencies when the relaxation time of the material is faster than the time sharing. In other words, during the switching from position 1 to position 2, the material may have relaxed back to the original position, which in the worst case degenerates the measurements. Thus, we have to compensate for this deviation.

The two aspects, although overlapping, have completely different origins and act on different regions of the spectra. The stiffness of the trap now is the single most important parameter which sets the limits of the experimentally accessible material properties. It can be shown that time-shared optical tweezer microrheology works best when the absolute value of the complex shear modulus $G(\omega)$ is much higher than the strength of the trap ($G_0 = \frac{k}{12\pi a}$). In that case the voltage change of the active trap dramatically surpasses the passive voltage change ($|V_1(t) + V_2(t)| >> |V_2(t)|$). However, in cases where the strength of the trap surpasses the complex G modulus of the sample, the bead jumps instantaneously between the two traps without deflection such that $V_1 \approx V_2$. Together, the trap stiffness should be chosen to obtain accurate measurements at low frequencies with stiffness not larger than 1000 times that of the assessed material. Finally, we have provided an analytical solution to understand the problem in the light of a dual-band amplitude-modulated waveform with carrier suppression. We also eventually provide a solution to correct for the observed deviations and recover the response of the particle as if we had measured the viscoelastic material in a standard, time-continuous way.

### 2 Supplementary Text 2: Derivation of the First Harmonic Approximation (FHA) for $\hat{\chi}_t(\omega)$

To derive the equation 30 the following steps are followed.

1. Compute $\hat{x}(\omega')$ by solving Eq. 27 where $\hat{x}_l(\omega')$ is given by Eq. 29.
2. Apply the inverse Fourier transform to obtain $x(t)$ and compute the signal read by the sensor $V(t) = \beta^{-1}(x_l(t) - x(t))$.
3. Extract the two signals sequences $V_1(n)$ and $V_2(n)$ by sampling $V(t)$ at the time points $t \in \{t_{1,n}\} = \{t_{1,n} = \tau_t \cdot (n + 1/4) | \, n \in \mathbb{N}\}$, and, $t \in \{t_{2,n}\} = \{t_{2,n} = \tau_t \cdot (n - 1/4) | \, n \in \mathbb{N}\}$ respectively.
4. Apply the discrete-time Fourier transform (DTFT) to obtain the signal spectrum $\hat{v}_1(\tilde{\omega})$ and $\hat{v}_2(\tilde{\omega})$.
5. Extract the Fourier coefficients $\hat{V}_1(\omega)$, $\hat{V}_2(\omega)$ and compute $\chi_t(\omega)$ by using Eq. 25.

#### 2.1 The movement of the bead $\hat{x}(\omega')$

The equation of motion was introduced in chapter 1.8,

$$\hat{x}(\omega') = \hat{\chi}_a(\omega') \, k \, \hat{x}_l(\omega'), \tag{27}$$

where $\hat{\chi}_a(\omega') = 1/(k + 1/\hat{\chi}(\omega'))$ was defined in chapter 1.2, and, $\hat{x}_l(\omega')$ is the Fourier transform (FT) of the First Harmonic Approximation (FHA) of the trap trajectory, discussed in chapter 1.8,

$$x_l(t) = \frac{x_1}{2} \exp(i\omega t)[1 + \sin(\omega_t \cdot t)]. \tag{28}$$

After FT the trap trajectory reads

$$\hat{x}_l(\omega') = \frac{x_1}{2}\left[\delta(\omega' - \omega) + \frac{\delta(\omega' - \omega_+) - \delta(\omega' + \omega_-)}{2\,i}\right], \tag{29}$$

with $\omega_\pm = \omega_t \pm \omega$ and where the following convention was used for the FT

$$\hat{F}(\omega') \quad = \textstyle\int_{-\infty}^{\infty} F(t)\exp(-i\,\omega'\,t)dt. \tag{61}$$

It should be noted that the laser trajectory $x_l(t)$ of Eq. 28 is not a real function as it should be. The Eq. 28 was chosen only to easily obtain the expression (FHA) of $\hat{\chi}_t(\omega)$. The intermediate results that follow in the next lines, such as the trajectory of the bead $x(t)$ and the sensor signal $V(t)$ will not be easy to interpret. If the reader wishes to use this approach to obtain these values, he or she must also repeat the calculation for the conjugate trajectory of the laser $x_l^*(t)$ and then combine the results to obtain the answer for the real trajectory of the laser $\mathrm{Re}\{x_l(t)\} = (x_l(t) + x_l^*(t))/2$. To go faster, the reader can make use of the property that the Fourier transform of the complex conjugate $F^*(x)$ is $\hat{F}^*(-\omega')$. In the remainder of the annex, only the solutions for the complex laser trajectory $x_l(t)$ will be treated. Therefore, the motion of the bead $\hat{x}(\omega')$ is simply given using Eq. 29 to express the trajectory of the laser $x_l(t)$ and replacing it in Eq. 27, it follows that

$$\hat{x}(\omega') = \frac{k\,\hat{\chi}(\omega')}{1 + k\,\hat{\chi}(\omega')}\,\frac{x_1}{2}\left[\delta(\omega' - \omega) + \frac{\delta(\omega' - \omega_+) - \delta(\omega' + \omega_-)}{2\,i}\right] \tag{62}$$

#### 2.2 The signal read by the sensor $V(t)$

Applying the inverse FT

$$F(t) \quad = \textstyle\int_{-\infty}^{\infty} \hat{F}(\omega')\exp(i\,\omega'\,t)d\omega', \tag{63}$$

to Eq. 62 it follows

$$\begin{aligned} x(t) \quad &= \quad \frac{k\,\hat{\chi}(\omega)}{1 + k\,\hat{\chi}(\omega)}\,\frac{x_1}{2}\exp(i\omega t) + \frac{k\,\hat{\chi}(\omega_+)}{1 + k\,\hat{\chi}(\omega_+)}\,\frac{x_1}{4i}\exp(i\omega_+ t) \\ &- \quad \frac{k\,\hat{\chi}(-\omega_-)}{1 + k\,\hat{\chi}(-\omega_-)}\,\frac{x_1}{4i}\exp(-i\omega_- t). \end{aligned} \tag{64}$$

On the other hand Eq. 28 can be written as

$$x_l(t) = \frac{x_1}{2}\exp(i\omega t) + \frac{x_1}{4i}\exp(i\omega_+ t) - \frac{x_1}{4i}\exp(-i\omega_- t). \tag{65}$$

By using the symmetry properties of the response function, i.e $\hat{\chi}(-\omega_-) = \hat{\chi}^*(\omega_-)$ and the above results the sensor signal is given by

$$\begin{aligned} V(t) &= \beta^{-1}(x_l(t) - x(t)) = \frac{\beta^{-1}}{1 + k\,\hat{\chi}(\omega)}\frac{x_1}{2}\exp(i\omega t) \\ &+ \frac{\beta^{-1}}{1 + k\,\hat{\chi}(\omega_+)}\frac{x_1}{4i}\exp(i\omega_+ t) - \frac{\beta^{-1}}{1 + k\,\hat{\chi}^*(\omega_-)}\frac{x_1}{4i}\exp(-i\omega_- t) \\ &= \frac{x_1}{2\beta}\exp(i\omega t)\left[\frac{1}{1 + k\,\hat{\chi}(\omega)} + \frac{1}{1 + k\,\hat{\chi}(\omega_+)}\frac{\exp(i\omega_t t)}{2i} - \frac{1}{1 + k\,\hat{\chi}^*(\omega_-)}\frac{\exp(-i\omega_t t)}{2i}\right]. \end{aligned} \tag{66}$$

To make shorter the notation the following quantity is introduced

$$\hat{Q}(\omega) = \frac{1}{1 + k\,\hat{\chi}(\omega)} \tag{67}$$

so that

$$V(t) = \frac{x_1}{2\beta}\exp(i\omega t)\left[\hat{Q}(\omega) + \hat{Q}(\omega_+)\frac{\exp(i\omega_t t)}{2i} - \hat{Q}^*(\omega_-)\frac{\exp(-i\omega_t t)}{2i}\right] \tag{68}$$

##### 2.3 Sampling of $V(t)$: the signals sequence $V_1(n)$ and $V_2(n)$

$V_1(n)$ is obtained by sampling the signal $V(t)$ at the time points $t \in \{t_{1,n}\} = \{t_{1,n} = \tau_t \cdot (n + 1/4)|\, n \in \mathbb{N}\}$. For this time values

$$\begin{aligned} \exp(-i\omega_t t_{1,n}) &= \exp(-i\omega_t\tau_t \cdot (n + 1/4)) = \exp(-i2\pi(n + 1/4)) = \exp(-i\pi/2) = -i \\ \exp(i\omega_t t_{1,n}) &= \exp(i\omega_t\tau_t \cdot (n + 1/4)) = \exp(i2\pi(n + 1/4)) = \exp(i\pi/2) = i \end{aligned} \tag{69}$$

then

$$V_1(n) = V(t_{1,n}) = \frac{x_1}{2\beta}\exp(i\omega t_{1,n})\left\{\hat{Q}(\omega) + \frac{1}{2}\left[\hat{Q}(\omega_+) + \hat{Q}^*(\omega_-)\right]\right\} \tag{70}$$

Similarly, $V_2(t_{1,n})$ is obtained by sampling the signal $V(t)$ at the time points $t \in \{t_{2,n}\} = \{t_{2,n} = \tau_t \cdot (n - 1/4) |\, n \in \mathbb{N}\}$. For this time values

$$\begin{aligned} \exp(-i\omega_t t_{2,n}) &= \exp(-i\omega_t\tau_t \cdot (n-1/4)) = \exp(-i2\pi(n-1/4)) = \exp(i\pi/2) = i \\ \exp(i\omega_t t_{2,n}) &= \exp(i\omega_t\tau_t \cdot (n-1/4)) = \exp(i2\pi(n-1/4)) = \exp(-i\pi/2) = -i \end{aligned} \tag{71}$$

and

$$V_2(n) = V(t_{2,n}) = \frac{x_1}{2\beta}\exp(i\omega t_{2,n})\left\{\hat{Q}(\omega) - \frac{1}{2}\left[\hat{Q}(\omega_+) + \hat{Q}^*(\omega_-)\right]\right\} \tag{72}$$

#### 2.4 Their Fourier spectra $v_1(\tilde{\omega})$ and $v_2(\tilde{\omega})$

$\hat{v}_1(\tilde{\omega})$ and $\hat{v}_2(\tilde{\omega})$ are obtained by discrete-time Fourier transform (DTFT) of the signals sequences $V_1(n)$ and $V_1(n)$. The DTFT is defined by

$$DTFT(F(n)) \quad = \textstyle\sum_{n=-\infty}^{\infty} \tau_t F(n)\exp(-i\tilde{\omega}\tau_t n). \tag{73}$$

As already explained for the FDE simulation where the FFT has been used to find numerically $\hat{V}_1(\omega_j)$ and $\hat{V}_2(\omega_j)$, the phase of the sequence transformation of $V_2(n)$ has to be corrected to take into account of the delay existing between the sample $V_1(n)$ and $V_2(n)$, see Eq. 24. Analogously

$$\begin{aligned} \hat{v}_1(\tilde{\omega}) &= DTFT\,(V_1(n)) \\ \hat{v}_2(\tilde{\omega}) &= e^{i\tilde{\omega}\tau_t/2} DTFT\,(V_2(n)) \end{aligned} \tag{74}$$

The attentive reader can note that the sign of the phase compensation of $\hat{V}_2(\tilde{\omega})$ is different from the one of Eq. 24. This because now $t_1(n) = t_2(n) + \tau_t/2$ while for the FDE simulation

$t_2(i) = t_1(i) + \tau_t/2$. Said that, it follows

$$\hat{v}_1(\tilde{\omega}) = \frac{\tau_t x_1}{2\beta} \left\{ \hat{Q}(\omega) + \frac{1}{2} \left[ \hat{Q}(\omega_+) + \hat{Q}^*(\omega_-) \right] \right\} \sum_{n=-\infty}^{\infty} \exp(i(\omega - \tilde{\omega})\tau_t n) \exp(i\omega\tau_t/4) \tag{75}$$

and by using the Poisson summation Formula

$$\hat{v}_1(\tilde{\omega}) = \frac{\tau_t x_1}{2\beta} \left\{ \hat{Q}(\omega) + \frac{1}{2} \left[ \hat{Q}(\omega_+) + \hat{Q}^*(\omega_-) \right] \right\} \exp(i\omega\tau_t/4) \sum_{m=-\infty}^{\infty} \delta(\omega - \tilde{\omega} - m\omega_t). \tag{76}$$

In the same way

$$DTFT\,(V_2(n)) = \frac{\tau_t x_1}{2\beta} \left\{ \hat{Q}(\omega) - \frac{1}{2} \left[ \hat{Q}(\omega_+) + \hat{Q}^*(\omega_-) \right] \right\} \exp(-i\omega\tau_t/4) \sum_{m=-\infty}^{\infty} \delta(\omega - \tilde{\omega} - m\omega_t) \tag{77}$$

and using Eq. 74

$$\hat{v}_2(\tilde{\omega}) = \frac{\tau_t x_1}{2\beta} \left\{ \hat{Q}(\omega) - \frac{1}{2} \left[ \hat{Q}(\omega_+) + \hat{Q}^*(\omega_-) \right] \right\} \exp(-i(\omega - 2\tilde{\omega})\tau_t/4) \sum_{m=-\infty}^{\infty} \delta(\omega - \tilde{\omega} - m\omega_t). \tag{78}$$

#### 2.5 The time-sharing linear response function $\chi_t(\omega)$

To evaluate Eq. 25 the Fourier components $\hat{V}_1(\omega)$, $\hat{V}_2(\omega)$ need to be extracted from their spectrum. For this purpose an operator with an arbitrary small finite spectral window of width $2\epsilon > 0$ is used

$$\hat{V}(\omega) = \frac{1}{2\epsilon} \int_{\omega-\epsilon}^{\omega+\epsilon} \hat{v}(\tilde{\omega}) d\tilde{\omega} \tag{79}$$

this gives

$$\hat{V}_1(\omega) = \frac{\tau_t x_1}{4\beta\epsilon} \left\{ \hat{Q}(\omega) + \frac{1}{2} \left[ \hat{Q}(\omega_+) + \hat{Q}^*(\omega_-) \right] \right\} \exp(i\omega\tau_t/4), \tag{80}$$

$$\hat{V}_2(\omega) = \frac{\tau_t x_1}{4\beta\epsilon} \left\{ \hat{Q}(\omega) - \frac{1}{2} \left[ \hat{Q}(\omega_+) + \hat{Q}^*(\omega_-) \right] \right\} \exp(i\omega\tau_t/4). \tag{81}$$

From Eq. 25 the linear response function $\chi_t(\omega)$ is estimated thought the formula

$$\hat{\chi}_t(\omega) = -\frac{2}{k}\frac{\hat{V}_2(\omega)}{\hat{V}_1(\omega)+\hat{V}_2(\omega)}. \tag{82}$$

By replacing the Eqs. 80 and 81 in Eq 82, it follows

$$\hat{\chi}_t(\omega) = -\frac{1}{k}\frac{\hat{Q}(\omega)-\frac{1}{2}\left[\hat{Q}(\omega_+)+\hat{Q}^*(\omega_-)\right]}{\hat{Q}(\omega)}. \tag{83}$$

Writing $\hat{Q}(\omega)$, $\hat{Q}(\omega_+)$, $\hat{Q}(\omega_-)$ by mean of a comun denominator,

$$\begin{aligned}\hat{Q}(\omega) &= \frac{1}{1+k\,\hat{\chi}(\omega)} = \frac{(1+k\,\hat{\chi}(\omega_+))(1+k\,\hat{\chi}^*(\omega_-))}{(1+k\,\hat{\chi}(\omega))(1+k\,\hat{\chi}(\omega_+))(1+k\,\hat{\chi}^*(\omega_-))} \\ &= \frac{1+k\,\hat{\chi}(\omega_+)+k\,\hat{\chi}^*(\omega_-)+k^2\,\hat{\chi}(\omega_+)\hat{\chi}^*(\omega_-)}{(1+k\,\hat{\chi}(\omega))(1+k\,\hat{\chi}(\omega_+))(1+k\,\hat{\chi}^*(\omega_-))},\end{aligned} \tag{84}$$

$$\begin{aligned}\hat{Q}(\omega_+) &= \frac{1}{1+k\,\hat{\chi}(\omega_+)} = \frac{(1+k\,\hat{\chi}(\omega))(1+k\,\hat{\chi}^*(\omega_-))}{(1+k\,\hat{\chi}(\omega))(1+k\,\hat{\chi}(\omega_+))(1+k\,\hat{\chi}^*(\omega_-))} \\ &= \frac{1+k\,\hat{\chi}(\omega)+k\,\hat{\chi}^*(\omega_-)+k^2\,\hat{\chi}(\omega)\hat{\chi}^*(\omega_-)}{(1+k\,\hat{\chi}(\omega))(1+k\,\hat{\chi}(\omega_+))(1+k\,\hat{\chi}^*(\omega_-))},\end{aligned} \tag{85}$$

$$\begin{aligned}\hat{Q}^*(\omega_-) &= \frac{1}{1+k\,\hat{\chi}^*(\omega_-)} = \frac{(1+k\,\hat{\chi}(\omega))(1+k\,\hat{\chi}(\omega_+))}{(1+k\,\hat{\chi}(\omega))(1+k\,\hat{\chi}(\omega_+))(1+k\,\hat{\chi}^*(\omega_-))} \\ &= \frac{1+k\,\hat{\chi}(\omega)+k\,\hat{\chi}(\omega_+)+k^2\,\hat{\chi}(\omega)\hat{\chi}(\omega_+)}{(1+k\,\hat{\chi}(\omega))(1+k\,\hat{\chi}(\omega_+))(1+k\,\hat{\chi}^*(\omega_-))},\end{aligned} \tag{86}$$

the numerator of Eq. 83 becomes

$$\begin{aligned}\hat{Q}(\omega)-\frac{1}{2}\left[\hat{Q}(\omega_+)+\hat{Q}^*(\omega_-)\right] &= \frac{-k\,\hat{\chi}(\omega)+k\frac{1}{2}[\hat{\chi}(\omega_+)+\ \hat{\chi}^*(\omega_-)]+k^2\,\hat{\chi}(\omega_+)\hat{\chi}(\omega_-)}{(1+k\,\hat{\chi}(\omega))(1+k\,\hat{\chi}(\omega_+))(1+k\,\hat{\chi}^*(\omega_-))} \\ &\quad \frac{-k^2\,\hat{\chi}(\omega)\frac{1}{2}\left[\hat{\chi}(\omega_+)+\hat{\chi}^*(\omega_-)\right]}{},\end{aligned} \tag{87}$$

then by using the definition of $\hat{\chi}_1(\omega)$, $\hat{\chi}_+(\omega)$, $\hat{\chi}^*_-(\omega)$ introduced in Chapter 1.8,

$$\hat{\chi}_1(\omega) = \frac{1}{2}\left[\hat{\chi}_+(\omega) + \hat{\chi}^*_-(\omega)\right], \tag{31}$$

$$\hat{\chi}_+(\omega) = \hat{\chi}(\omega_t + \omega) = \hat{\chi}(\omega_+), \tag{32}$$

$$\hat{\chi}^*_-(\omega) = \hat{\chi}^*(\omega_t - \omega) = \hat{\chi}^*(\omega_-), \tag{33}$$

and replacing Eq. 87 and Eq. 84 in Eq. 83 the FHA expression of $\chi_t(\omega)$ follows

$$\hat{\chi}_t(\omega) = \frac{\hat{\chi}(\omega) - \hat{\chi}_1(\omega) + k\left[\hat{\chi}(\omega)\hat{\chi}_1(\omega) - \hat{\chi}_+(\omega)\hat{\chi}^*_-(\omega)\right]}{1 + 2\,k\hat{\chi}_1(\omega) + k^2\hat{\chi}_+(\omega)\hat{\chi}^*_-(\omega)} \tag{30}$$

### 3 Supplementary Text 3: Interpretation of the parameters derived from the cytoplasmic rheology

The prediction of the rheological properties based on their constituent protein expression has been difficult if not impossible to date, especially *in vivo*. Thus, these properties need to be measured in order to understand how cells and their organelles react to mechanical stresses[5]. The traditional rheological models such as the Kelvin-Voigt (KV) and Maxwell models have been used extensively to describe the essence of the mechanical properties and indicate whether a material predominantly behaves as a solid or a liquid, respectively. However, their assumptions do not hold for complex fluids or solids such as the cytoplasm and diverse cellular organelles, or are only valid over a narrow range of frequencies. A large number of experiments in the past decades suggest that rheology of biological systems is described by a power law or a combination of power laws [24]. A frequency-dependent power law results from the fractional rheological element, the springpot [24], for which the G modulus reads $\hat{G} = C_\alpha(i\omega)^\alpha$ (see Section 1.6), where $\alpha$ captures all viscoelastic responses from purely elastic ($\alpha \to 0$) to purely dissipative ($\alpha \to 1$) and $C_\alpha$ is the strength of the response. Interestingly, the usage of the fractional formalism allows us to describe the rheological spectrum with fewer parameters: two per springpot, i.e. power law unit. It has been shown that the fractional Kelvin-Voigt model describes the rheological frequency response of living cells very well over the frequency range that is explored in an active optical tweezer microrheology experiment[29, 30, 1, 31, 32]. Similar to the conventional KV model, this model can be conceptualized as a parallel arrangement of two springpots, described by parameters ($C_\alpha$, $\alpha$) and ($C_\beta$, $\beta$), whose response dominates at low and high-frequency stimulations, respectively. For clarity and convenience, we will solely regard two springpot elements and refer the reader to the Supplementary Text 1 for the mathematical treatment of higher order models.

The two exponents $\alpha$ and $\beta$ in the model describing the Kelvin-Voigt material define the frequency scaling of the mechanical properties within the low and high frequency regimes, respectively. Herein, $\alpha$, which tends to zero for a perfect solid material without visco-elastic behavior, indicates how well the material responds conservative to low frequency stimulations. This is because the perturbation is performed much slower than the viscous relaxation timescale, thus the viscous contribution to the mechanical response has no effect. Note, that the opposite is true for a Maxwell material (and no elastic contribution is measured due to the 'flow' of the dashpot). On the other hand, $\beta$, which tends to 1 for a perfect liquid without restoring forces, describes how well the material responds as a liquid for high frequencies. The prefactors to both exponents, $C_\alpha$ and $C_\beta$, are thus the magnitude of the elastic and liquid behavior and can be conceptualized as an elastic or viscous modulus. Thus, a rheological spectrum affords insights into the forces that stabilize the cell's interior with only four parameters. Our notion that the fractional Kelvin-Voigt model fits the rheological spectrum of living cells best is not specific to *C. elegans* and zebrafish but has been previously used with the majority of cell types [33, 1, 29, 32]. Recently, this parameterization, together with the analysis of the out-of-equilibrium fluctuational energy, has been used to extract a rheological properties of different cultured cell lines[29]. Here we show how these properties changes between different cellular organelles and with age of an organism.

#### 3.1 Mutations of the nuclear envelope sensitize *C. elegans* intestinal cells to age-related changes in viscosity

With the aim to understand how cytoplasmic mechanics changes with the age of an organism, we applied the fractional Kelvin-Voigt model to the rheological spectrum derived from optical tweezer measurements directly inside intestinal cells of living *C. elegans*. We compared two different time-points and four different genotypes in order to understand how the cytoplasmic properties change with age and in mutants of the nuclear lamina. We found that control cells do not experience any significant rheological changes during the first eight days of adulthood, as indicated by all four rheological parameters (Supplementary Table 2). For all measurements, we observed a much 'stiffer' cytoplasm compared to zebrafish cells, indicated by larger values of $C_\alpha$ and $C_\beta$ but also a low exponent $\beta \approx 0.5$. This exponent of the high frequency component is closer to published values of living cells[34] rather than that of semiflexible polymers *in vitro* ($\alpha \approx 0.75$).

Lamin is a protein that supports the nuclear envelope and anchors chromatin to the nuclear membrane[35]. When we compared control cells to cells from animals expressing a dominant negative GFP::lmn-1 construct, we found a strong decrease in their cytoplasmic viscosity ($C_\beta$, p=0.049) and modest, insignificant increase in elasticity (decrease in $\alpha$, p=0.06; Extended Data Fig. 10f). Their overall solid and liquid character remained largely unchanged, even though we noticed a subtle decrease in $\alpha$, which could be interpreted as the emergence of a more solid-like character during age. Interestingly, in presence of the dnLMN-1, we begin to observe an elastic plateau at low frequencies (Extended Data Fig. 10a), similar to what was observed in cytoskeletal networks prepared in vitro[36]. The emergence of that plateau has previously been observed in ATP depleted cell and thus might hint towards a potential mechanism of how dnLMN-1 interferes with metabolic activity that maintains a fluid character of cytoplasmic mechanics[34]. The expression of the lamin construct itself had no further consequences on the mechanical properties during age and remained unchanged during the first eight days of adulthood.

We then performed rheology in mutants with defects in EMR-1/Emerin and LEM2/LEM-2, integral membrane proteins of the nuclear envelope that integrate the lamin nucleoskeleton and chromatin. For both mutants, after 8 days we observed a strong decrease in $C_\beta$, inducing a shift in the crossover frequency between the storage and the loss modulus to higher frequencies. This suggests that the aging process results in a tissue that is more compliant to mechanical stress, even for fast insults. In general, in *lem-2* mutants, we did not detect significant changes in rheology in young adults compared to control cells (p=0.3), and unchanged parameters in old animals. Viscosity slightly decreased during age ($C_\beta$, p=0.07) indicating that *lem-2* has minor roles in stabilizing the parameters of cytoplasmic rheology during age. This indicates that *lem-2* has little to no effect on stabilizing cytoplasmic rheology.

This picture slightly changed when we studied *emr-1*. Whereas we did not observe a significant change of $C_\beta$ between *emr-1* mutants and control cells in young and old adults, *emr-1* had a strong effect on cytoplasmic rheology in 8 day old adults, especially on $\alpha$, suggesting a more fluid cytoplasm, (p=0.003). Like wildtype animals in general, we observed a consistent decrease in viscosity ($C_\beta$, p=0.002), a modest decrease in elasticity ($C_\alpha$, p=0.08) and a strong and significant increase in both exponents $\alpha$ and $\beta$ (p=0.002 and p=0.03, respectively) with age. This suggests that *emr-1* loss causes a loss in conservative forces in the cytoplasm and leads to a more liquid-like material when stimulated at high frequencies ($\beta$ closer to 1) and more dissipative behavior at low frequency (alpha>0).

Taken together, we observed a strong defect in cytoplasmic rheology of the premature ageing mutants *emr-1* and the dominant negative GFP::LMN-1 construct, leading to an overall more dissipative and less viscous (more fluid) material. Contrary to intuition, we also observed the emergence of an elastic plateau at low frequencies in the *lmn-1* construct, whose biological significance needs to be addressed in future studies. Future work also needs to address how proteins of the nuclear envelope influence cytoplasmic mechanics during age.

#### 3.2 Rheology of the zebrafish cytoplasm is unaffected by bead insertion into the nucleus

Our surprising findings that optically trapped microsphere can be inserted into the nucleus to probe the internal rheology of the nucleoplasm motivated us to verify that cells remained viable and healthy during the measurement and are not immediately affected by the procedure. To do so, we performed paired measurements on cells that internalized two beads - a first measurement with one bead in the cytoplasm and at the nuclear interface before insertion and a second measurement after bead insertion. Using this procedure, we identified little to no changes in cytoplasmic and nuclear rheology. All except the $C_\alpha$ parameter are seemingly unchanged (Supplementary Figure S25a). Next, we verified that the nuclear envelope remained intact after the procedure or if the envelope ruptured irreversible during the procedure. To do so, we expressed GFP with a nuclear localization signal and stained the DNA with Hoechst and reasoned that envelope rupture would be visible by GFP and DNA leakage. Neither was observed in any of the experiments conducted (Suppl. Fig. 25b). Previously, we have shown that nuclear deformation precedes activation of a prominent mechanotransduction pathway[37] visible as an activation and relocalization of myosin II to the cell surface. Thus, we asked whether the insertion and the large strain associated to it activates myosin II (using transgenic Tg(actb2:Myl12.1-eGFP)). Compared to untreated control cells, we were not able to observe a substantial relocalization of myosin II (Suppl. Fig. 25c), indicating that the procedure does not activate intracellular mechanotransduction pathways and an associated change in rheological properties. Lastly, we verified that the cells remained viable and healthy for up to 1h after the insertion. Neither did the cells accumulate the apoptotic cell marker Annexin V at the cell surface, nor did the cell have a substantial defect in undergoing mitosis (Suppl. Fig. 25d). Taken together, the bead insertion into the nucleus did not affect cell viability and mechanical state immediately and for up to 1h after completion of the procedure.

### 4 Supplementary Text 4: Aberrations of the optical microscopy and optical trapping performance inside *C. elegans*

Our microscope objective (Plan Apo, Nikon WI 1.2) is infinity-corrected for water. However, being that both visible and IR light go through a thick layer of biological tissue that has a refractive index slightly different than 1.33, we wondered about the effect of optical aberrations. Here we assume that the *C. elegans* tissue does not have significant optical dispersion compared to that of water, for which the confocal imaging plane and the optical trapping plane still coincide deep inside the animal.

First, we addressed the question of whether spherical aberration due to the refractive index mismatch between the *C. elegans* tissue and the objective immersion medium (water), would lead to a significant drop in the optical trapping stiffness. To do so, we estimated the curvature of the cuticle of different age-matched, immobilized animals after staining them with DiI with slight modifications as previously described[38]. In short, *C. elegans* at Day 1 and Day 8 adults were collected and washed using a solution of 0.5% Triton X-100 and M9. Then, animals were incubated in 0.33 mg/mL DiI solution (Sigma Aldrich, 42364) for three hours, in agitation and in the dark. Three washes with M9 were made prior to animal mounting. Then animals were

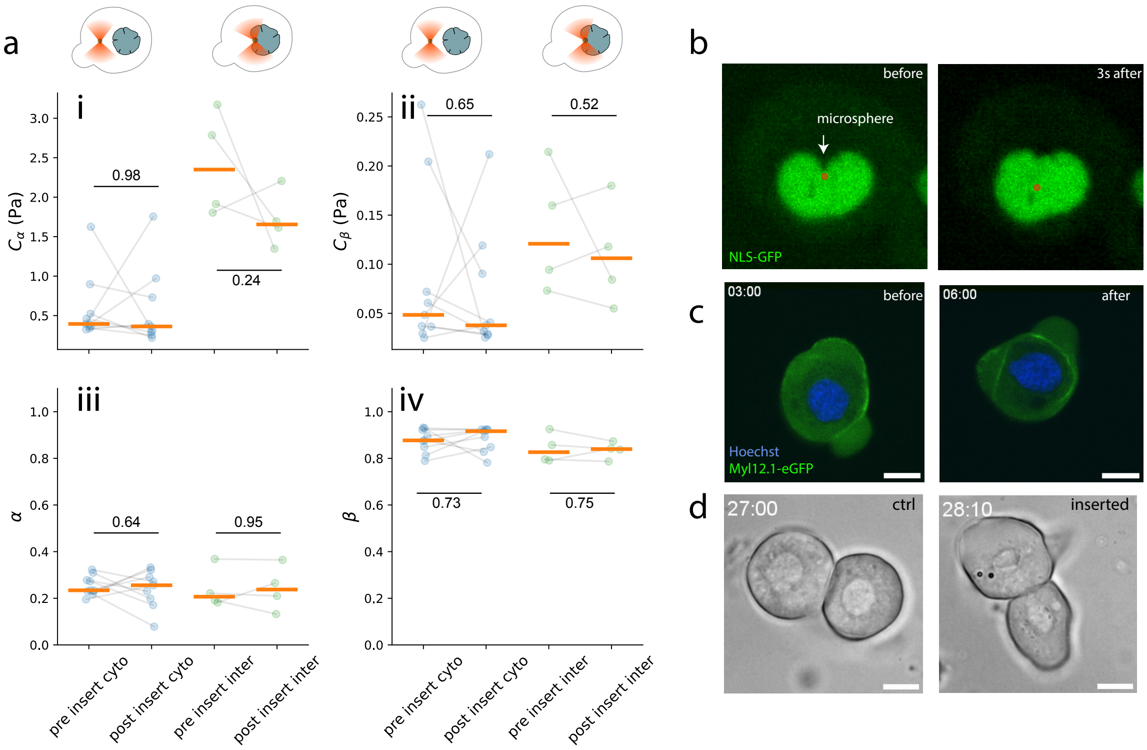

**Supplementary Fig. 25**: **a,** Rheology of the cytoplasm and interface remain unchanged by large nuclear deformations. The low-frequency component of the fractional Kelvin-Voigt fit to the rheological spectrum obtained from two different microspheres in the same cell before and after insertion into the nucleus. i) prefactor $C_\alpha$; ii) prefactor $C_\beta$; iii) Exponent $\alpha$; iv) Exponent $\beta$. Dots connected by lines correspond to sequentially paired measurements in the same cell. Numbers on the horizontal lines indicate p-value derived from a paired t-test on the indicated data pairs. **b,** Snapshots of a cell nucleus labeled with soluble NLS-GFP right before and after insertion. No GFP leakage is observed. Images from Video S5 **c,** Representative picture of a cell expressing Myl12.1-eGFP. Left: before insertion; right: 6 min after insertion. No myosin II activation is observed compared to untreated controls. **d,** Representative cells with an nucleus-inserted microsphere are able to successfully undergo mitosis. Endpoint from Video S6 (ctr) and S7 (inserted). Scale bars $=$ of $10\mu$ m

immobilized under the same conditions as for TimSOM experiment in the confocal microscope to record multiple Z-stacks at different parts of the worm body. From orthogonal sections, the width and height of each portion were measured and width/height ratios were calculated for each condition.

Interestingly, we found that the worm was considerably flattened in the optical trapping chamber (see Methods) and that the surface of the worm cuticle facing the microscope objective was completely flat (Supplementary Fig. 26). Simulations in Zemax carried out by the manufacturer showed no effect for an axial movement of the trap, though a substantial decrease in the trapping efficiency was noticed when laterally approaching the edge of the worm. The trap stiffness was reduced by 20% when placed at the very edge of the nematode, i.e. at the cuticle. (Data not shown.)

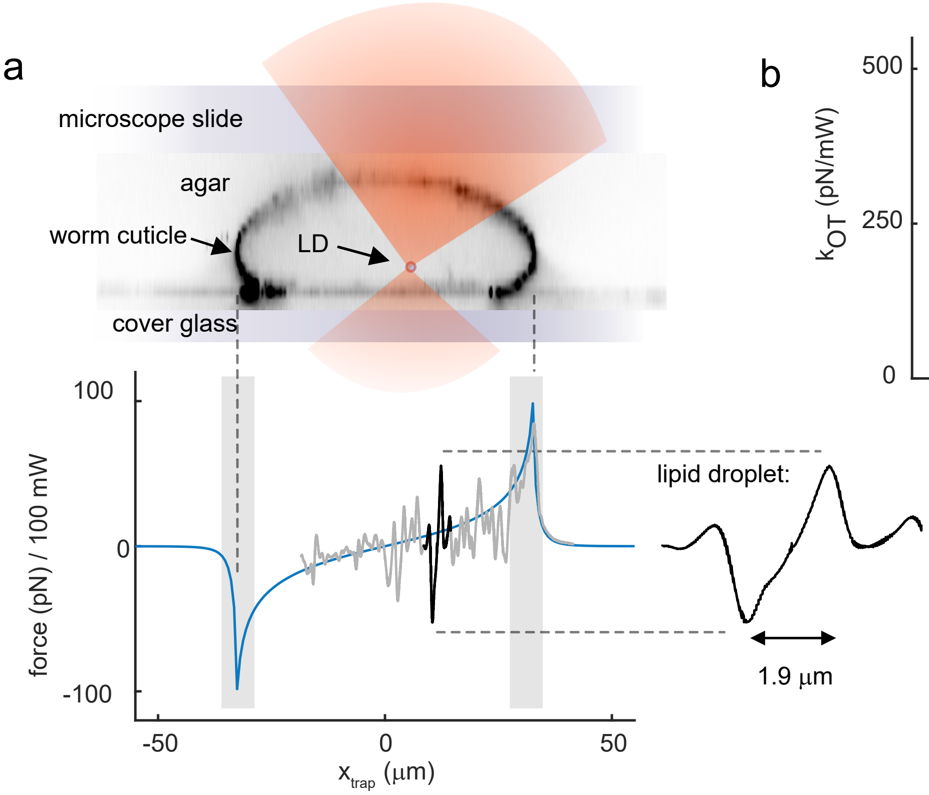

**Supplementary Fig. 26**: Variation of light momentum across the *C. elegans* section. **a:** Top - Cross section of a worm sample embedded into an agar pad and mounted between a microscope slide and a cover glass. The worm cuticle is tagged with DiIC18(3) (known as DiI; see below). Bottom - Optical force for a 100-mW laser trap. The blue line indicates the simulation under the geometrical optics approximation (see text below) and the gray line shows the force measured while scanning the optical trap across the animal. In black, the force profile around the trapped lipid droplet, with a stiffness of $k_{\rm LD} = 50.3 \rm pN/\mu m$, is indicated. **b:** Trap stiffness on a microsphere probing the cytoplasm, nuclear interface and nuclear interior. P-values are derived from a two-sided t-test. Boxes indicates the central 50% of the data around the median (horizontal line), and the whisker delimit the 10 and 90th percentile.

Second, we tackled the variation in the light momentum of the trapping laser through the quasi-cylindrical shape of the worm. The aforementioned mismatch between the refractive index of the worm and that of the water immersion medium, and between the worm and the agar medium layer in which the worm is embedded, lets us think there might be a variation in the momentum of the incoming beam of light, in particular, when moving it transverse to the cylindrical axis. Such change in light momentum could be misinterpreted as contributing to the actual trapping force emerging from the photon momentum exchange with the trapped lipid droplet. From 3D stacks of worms mounted into our agar pads (Supplementary Fig. 26a), we measured the length of the long sectional axis and did a simplistic approximation by considering them as a cylinder. Using the Optical Trapping in Geometrical Optics Toolbox (OTGO, see Ref. [39]), we calculated the force profile with a refractive index for the worm of $n_{\text{worm}} = 1.379$ [40] and an RI for the agar medium of $n_{\text{agar}} = 1.336$ [41]. As expected for a sample with radius much larger than the wavelength, $R >> \lambda$, the force varied smoothly around the central position of the cylindrical worm and steepened when approaching the edges. With a laser power of $P = 100$ mW, we measured a slope of 0.94 pN/µm which was 50-fold smaller than the optical trapping stiffness of an exemplary lipid droplet (Supplementary Fig. 26a, black line), for which we can state that the variation in light momentum measured with our direct force sensor mostly corresponds to the actual force the trap is exerting onto the lipid droplet probing tissue microrheology. However, the distance to the worm cuticle was always kept above 5 µm. Below this point, the slope of the light momentum baseline steepens upwards from 10% of the average trap stiffness of the lipid droplet probes (shaded area in Supplementary Fig. 26a).

Finally, we checked for trap stiffness variability of the polystyrene microspheres used in our experiments in living zebrafish embryonic stem cells (Supplementary Fig . 26b). For the same bead trapped in the cytoplasm, nuclear interface and nuclear interior (see Fig. 3a-c in Main Text), no significant variation of the optical tapping stiffness was observed.

### 5 Supplementary Text 5: General consideration for intracellular rheology with optical tweezers

#### 5.1 Heating

A focussed infrared laser light, depending on the power and the trapped particle, may lead to local heating in the sample, which may affect enzymatic functions, stress response, and cell rheological properties [42, 43, 44]. To characterize the effect of the trapping laser on cell mechanics we have systematically varied the trapping power ranging from 50-250mW on the sample plane while performing the standard rheology routine in zebrafish progenitor cells. We found a change in rheology, most notably in the low frequency elastic part, described by $C_\alpha$ and $\alpha$ for high laser powers (higher than 120mW, Extended Data Figure 4e). Importantly, this increase is most likely due to the laser power and not a history effect of a repetitive rheology routine, as a similar protocol with a constant 90mW laser power did not lead to a similar effect (Extended Data Figure 4d). Note, the relation between the change in these parameters and the laser power may strongly depend on the sample and cell type under investigation. Our values of 60-90mW are on the lower end of the spectrum of laser powers that were used in the literature to measure intracellular mechanics in living systems, which range from 30mW-150mW in tissue culture cell[1, 45] over 200mW in *Drosophila* [46] and up to 750mW in chick embryos[47]. We also noticed a strong dependence on the maximum laser power that can be used on different biomolecular condensates. For MEC-2 droplets, the maximum laser power was 90mW. Beyond, the droplet started to visibly change refractive index, visible as Schlieren throughout the droplets. For CPEB4 droplets, the maximum laser power that could be applied without this effect was 50mW.

#### 5.2 Sensitivity of the technique

Even though the force measurements of optical tweezers are limited to $\approx$ 200pN, relatively large stresses can be measured due to the exceptional sensitivity of the BFPI to detect nanometer and even subnanometer displacements. Thus, a main factor limiting the detection is mechanical noise, and of course cellular activity. In order to assess the limits of our setup, we casted PAA gels with different stiffnesses up to 30kPa using published recipes as described in the methods section. We then performed the rheology routine using 300mW at the sample plane and recorded the motion of the bead using the static detection laser while driving it actively with the trapping laser. As expected, the gel could be characterized as a Kelvin-Voigt viscoelastic solid material (Supplementary Fig. 27). The resulting signals were Fourier transformed to obtain a peak in the frequency of the driving laser. Using the Fourier-transformed force and displacement has the advantage that only oscillations at the applied frequency are evaluated[1]. Importantly, for all frequencies tested, we observed a corresponding peak in the static, detection trap, indicating that the displacement of the microsphere could be recorded. We note that for lower power or stiffer gels, the amplitude of the peaks measured with the detection trap decreases. Importantly, if the data points did not exceed the noise level, they remained undetected and were omitted from the analysis. With this analysis we were able to measure the ≈20kPa low frequency modulus of the PAA gel which reached up to 100kPa shear modulus at the highest driving frequency. As these measurements were perform close to maximum available laser power, the maximum value inside cells is likely a somewhat lower. However, all measurements of the interior of living cells yielded shear moduli in the range of 0.1-100Pa and are thus several orders of magnitude below our resolution limit. We should mention at this point that our setup suffers from mechanical noise in the 0.5-1kHz range, which originated from the ventilation in the Biosafety level S2 laboratory environment and coupled into the measurement of the stiffest gel (performing the measurement in damped media does not show this effect). Thus, several data points needed to be omitted in the analysis that did not exceed the noise levels.

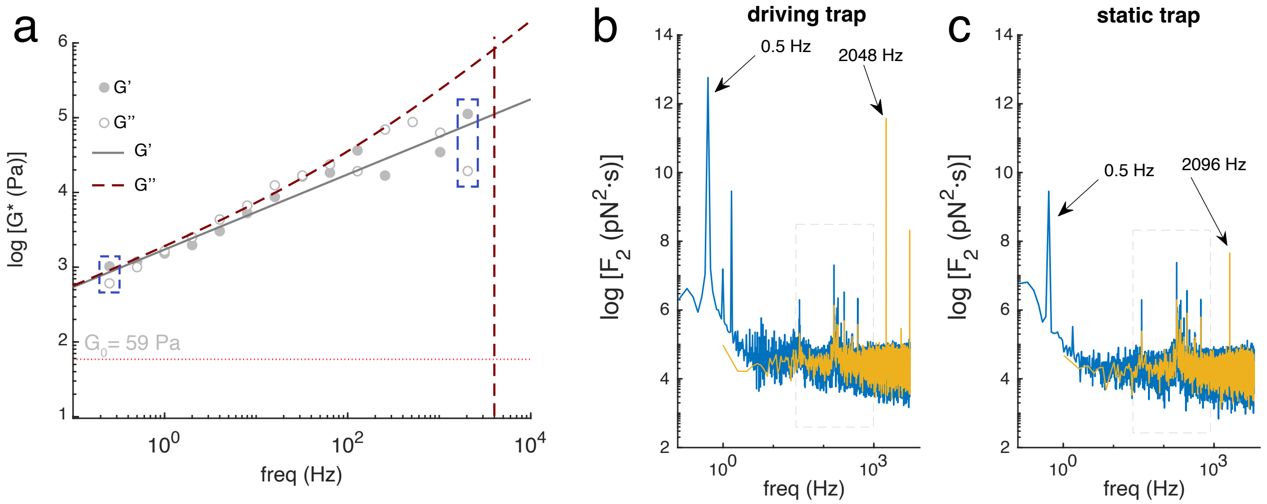

**Supplementary Fig. 27**: Noise, not force limits the detection sensitivity of the technique. **a,** Representative rheological spectrum of a PAA gel measured with TimSOM. Solid and dashed line indicate fractional Kelvin-Voigt fit to the storage and loss curves. Some data points were omitted due to mechanical noise. **b, c,** Power spectral density of the signal from the (b) driving trap and the (c) static detection trap at the lowest (0.5Hz) and highest (4096Hz) frequency yielding the right-most 100kPa data point. Note, the mechanical noise in the central region due to the ventilation of the biosafety level 2 lab environment.

#### 5.3 Dependence of the rheological results on probe size

Assuming a continuous viscoelastic material, the size of the probe determines the resistance of the material against the movement of the bead driven by the trap. This is taken care of in the calculation of the complex shear modulus from the response function according to Eq. 1b. However, in reality, the cytosol is not a homogeneous, isotropic continuum material but consists of cytoskeletal meshwork infiltrated by a viscous fluid, with membraneless compartments, endosomes, polyribosome, etc. Thus, one would expect a different response for microspheres smaller than the mesh size compared to the response measured with microspheres that are larger than the pore size of the mesh[48]. In particular, one would expect a decrease of the scaling exponent $\alpha$ concomitant with the particle radius $a$, given a particular mesh size[49]. Recent estimates suggest that the cytoplasmic pore size is in the order of few nanometers (Ref. [50], hydraulic pore size = 15nm), a reason why we assume that the cytoplasm can be modeled as a continuum with our experimental approach.

More recently, magnetic tweezers have been used to drag probes of the size approaching the scale of the entire cell (up to 12.5% volume fraction) and found a strong effect of mechanical signatures on probe size and displacement amplitude. This size-dependent viscoelasticicty was found to emerge from hydrodynamic coupling between the object and the cell boundary[51] (see also discussion below).

#### 5.4 Rheology measurements in confined volumes.

TimSOM has no particular limit in the probe size that can be used to explore intracellular rheology. The limits for stable trapping as a function of bead size and refractive index can be conveniently explored using one of the many computational toolboxes[39, 52]. We note though, that for larger bead size a larger movements the linear stress/strain relation of the surrounding elastic material may be violated, thus requiring new models to analyze the data. Further, larger beads (and larger displacement amplitudes $x_0$) will move more cytosol, and its resistance to movement may be influenced by the spatial constraints such as the static cell boundary. This may require to take wall effects into account. Such wall effects are well studied in fluid mechanics and can lead to an increase in effective viscosity, if the particle is closer than 5 radii from the boundary[53], or if the particle size approaches that of the confining volume. The effects of probe size on the rheological measurements have been recently explored in large sea urchin oocytes[51], which do not contain visible cytoplasmic compartmentalizations, actin cortex or cytosol movement that would influence a rheological measurement. Whereas the movement of small probes did not create detectable cytoplasmic flows, larger probes did. Consequently, the fluid flow induced by larger moving probes generates shear flows from hydrodynamic interactions between the object and cell boundaries, which effectively increases the drag force and thus viscosity of the measurement. This can be understood with a simple correction factor modifying Stoke's law that depends on the ratio $\lambda$ between probe size $a$ and the size of the container $R$ in which the measurement was performed (in our case the cell or nucleus):

$$\gamma(\lambda) = 6\pi\eta \, aC(\lambda), \qquad \text{with} \tag{88}$$

$$C(\lambda) = \frac{4(1-\lambda^5)}{4 - 9\lambda + 10\lambda^3 - 9\lambda^5 + 4\lambda^6} \tag{89}$$

where $\eta$ is the bulk fluid viscosity of the Newtonian liquid in an infinitely large container. For $\lambda << 1$, e.g. small probes but also for small displacements, this factor approaches 1 and thus does not markedly change the rheological measurement in the confined volume compared to an infinite container. With our microspheres $a = 0.5$µm, inside the cell $R = 15.4 \pm 0.004$µm, $\lambda$ becomes 0.03 and thus $C(\lambda) = 1.07$ for the cytoplasm and $C(\lambda) = 1.18$ inside the nucleus ($R = 7.1 \pm 0.2$ µm). We thus conclude that the wall effect does not have a strong influence on our measurements.

One may argue that the initial position of the microsphere closer to the cell membrane or the nuclear interface may induce a similar effect. Indeed, for large objects, this hydrodynamic coupling existed and both drag and stiffness increased if the probes were closer to the cell surface. However, for small probes, this effect was in-existent[51].

### 6 Supplementary Data Tables

**Supplementary Data Table 1: Statistics of the rheology in zebrafish** P-values derived from a Mann Whitney U test, comparing the parameters $\alpha$, $\beta$, $C_\alpha$ and $C_\beta$ extracted from the fractional Kelvin Voigt fit to the rheological spectrum in Fig. 3 and Extended Data Fig. 5. Some combinations were acquired 'paired' with the same microsphere inside the same cell, which are indicated in the figure legends of Fig. 3 and as blue fields in the tables.

**Supplementary Data Table 2: Statistics of the rheology in *C. elegans.*** P-values derived from a Kruskal-Wallis test followed by a Dunn test for pairwise comparison of the indicated combinations, comparing the parameters $\alpha$, $\beta$, $C_\alpha$ and $C_\beta$ extracted from the fractional Kelvin Voigt fit to the rheological spectrum in Fig. 4. Includes number of measurements on different droplets (N) and replicates (n, independent experimental days).

**Supplementary Data Table 3: Rheological routine for water and glycerol** Set of parameters used for the active microrheology measurement in water and glycerol.

**Supplementary Data Table 4: Recipes for the preparation of elastic poly-acrylamide hyrogels** Tables describing the content of acrylamide and bisacrylamide to produce hydrogels with varying stiffness.

**Supplementary Data Table 5: Parameters of the rheological routine for protein condensates, zebrafish cells and *C. elegans* animals** Set of parameters used for the active microrheology measurement in zebrafish progenitor cells and the active microrheology measurement on cytoplasm of intestinal cells in *C. elegans.* From left to right, columns correspond to oscillating frequency, amplitude, duration and measurement duration. We systematically took, at least, eight cycles for the measurement and the measurement setpoint took place 1 s after the oscillation started. Trap power is always $P_{trap} = 60 - 90$ mW.

**Supplementary Data Table 6: List of DNA sequences** Oligonucleotides for cloning the RFP-Lamin A (zebrafish) and CRISPR reagents to produce *lmn-1::GFP11* (*C. elegans*). crRNA and single-stranded oligonucleotide repair construct were being used.

**Supplementary Data Table 7: List of *C. elegans* strains used in this study**

### 7 Supplementary Videos

#### Supplementary Video 1

Representative video of the dual trap assay to measure surface tension and mechanical properties of protein droplet. Scale bar = 5um.

#### Supplementary Video 2

Representative video of the TimSOM assay to measure bulk mechanical properties of protein droplet. Scale bar = 5um.

#### Supplementary Video 3

Representative video of the complete rheology routine inside a zebrafish progenitor cell. Scale bar = 10um.

#### Supplementary Video 4

Representative video of a zebrafish progenitor cell subjected to a bead insertion routine, after applying a 100 pN force clamp for $\approx$ 5s. Force clamp started after 5s and entered the nucleus after 11s, after which the fc routine is abruptly stopped. Scale bar = 10µm. Green = lap2beta; blue= Hoechst 33342. Time label in seconds.

#### Supplementary Video 5

Representative video of a zebrafish progenitor cell stained with Hoechst 33342 to highlight nuclear DNA, which expresses nuclear-localized, soluble GFP (NLS-GFP). No obvious leakages of DNA and soluble GFP from the nucleus into the cytosol after bead insertion is visible, indicating no significant damage to the nuclear envelope. Video representative for N=3 cells. Scale bar = 10µm. Blue=Hoechst 33342; Green=NLS-GFP.

#### Supplementary Video 6

Representative video of a dividing zebrafish progenitor cell. Scale bar = 10µm.

#### Supplementary Video 7

Representative video of a dividing zebrafish progenitor cell after insertion of a microsphere into the nucleus. Scale bar = 10µm.
